# The small GTPase ARF3 controls metastasis and invasion modality by regulating N-cadherin levels

**DOI:** 10.1101/2022.04.25.489355

**Authors:** Emma Sandilands, Eva C. Freckmann, Alvaro Román-Fernández, Lynn McGarry, Laura Galbraith, Susan Mason, Rachana Patel, Jayanthi Anand, Jared Cartwright, Hing Y. Leung, Karen Blyth, David M. Bryant

**Author notes:** Correspondence: DMB.

## Abstract

ARF GTPases are central regulators of membrane trafficking that act by controlling local membrane identity and remodelling to facilitate vesicle formation. Unravelling ARF GTPase function is complicated by the overlapping association of ARFs with guanine nucleotide exchange factors (GEFs), GTPase-activating proteins (GAPs), and numerous interactors. The extent to which redundancy is a major factor in ARF function or whether individual ARF GTPases make unique contributions to cellular behaviour remains unclear. Through a functional genomic screen of 3-Dimensional (3D) prostate cancer cell behaviour we explore the contribution of all known ARF GTPases, GEFs, GAPs, and a large selection of interactors to collective morphogenesis. This revealed that the ARF3 GTPase regulates the modality of invasion, acting as a switch between leader cell-led chains of invasion or collective sheet movement. Functionally, the ability of ARF3 to control invasion modality is dependent on association and subsequent control of the junctional adhesion molecule N-cadherin. *In vivo*, ARF3 levels acted as a rheostat for metastasis from intraprostatic tumour transplants and ARF3:N-cadherin expression can be used to identify prostate cancer patients with metastatic, poor-outcome disease. Our analysis defines a unique function for the ARF3 GTPase in controlling how cells collectively organise during invasion and metastasis.

## Main

ARF GTPases are highly evolutionarily conserved regulators of membrane trafficking^1, 2^. ARF proteins co-ordinate membrane trafficking by regulating the local identity of the membrane to which they are recruited, such as through modulation of phospholipid composition via phosphatidylinositol kinases^1, 3^. This allows the recruitment of adaptor and coat proteins, facilitating membrane protein clustering and membrane deformation and ultimately leading to vesicle budding of encapsulated cargoes^1^. ARF GTPases are therefore central players in the localisation of most membrane proteins and have emerged as key regulators of polarised cell behaviours underpinning cancer cell growth and metastasis^4, 5^.

ARF GTPases cycle between GDP- or GTP-bound forms with the assistance of guanine nucleotide exchange factors (GEFs) and GTPase-activating proteins (GAPs)^6^. Rather than consideration of ARF-GTP as active, and ARF-GDP as inactive, the full cycle of GTP-loading of an ARF by a GEF to allow recruitment of effectors, followed by nucleotide hydrolysis by a GAP to return to GDP-ARF is required for ARF function^2^. Therein, an ARF GAP acts as both terminator and effector of the ARF GTPase cycle. In humans, five ARF GTPases are divided into three classes based on homology: Class I (ARF1, ARF3), Class II (ARF4, ARF5), and Class III (ARF6).

A complication in unravelling ARF GTPase function is their high degree of similarity in sequence and consequently their overlapping ability to associate with GEFs, GAPs and interactors^2^. For instance, Class I ARFs (ARF1, ARF3) differ by seven amino acids in their N- and C-terminal regions, while their core ARF domain regions are identical. Moreover, of the 17 GEF and 23 GAP proteins, many of these share the ability to modulate nucleotide association on most ARFs *in vitro*. ARF GTPases can also act in amplifying loops, with a GEF acting as an ARF-GTP effector to activate further ARFs^7, 8^. This complexity makes it difficult to predict how ARFs and their regulators contribute to cellular behaviour from individual single interactions and sets the stage for a systems-level analysis to identify how these potentially overlapping components functionally contribute to morphogenesis.

Here, we present a system-level characterisation of ARF GTPase function in collective cellular behaviours using large-scale timelapse imaging of the morphogenesis of prostate cancer cells in 3-Dimensional (3D) culture, machine learning to identify distinct resulting phenotypes, and molecular characterisation of behaviours. This work identifies a key role for the poorly studied ARF3 GTPase in controlling how cells collectively organise into distinct phenotypes. ARF3 controls the modality of invasion, between leader cell-led chains of invasion versus collective sheet movement, by associating with and controlling levels of the junctional protein N-cadherin. ARF3 therefore acts as a rheostat for the modality of invasion, which regulates metastasis *in vivo* and can be used to identify prostate cancer patients with metastatic, poor-outcome disease. Our approach therefore allows elucidation of distinct functions of ARF GTPases in collective morphogenesis.

## Results

### A 3D screen for ARF GTPase contribution to collective cancer cell behaviour

We interrogated the functional contribution of ARF GTPases, their GEFs, GAPs and known interactors and effectors, which we term the ‘ARFome’, to cancer cell morphogenesis (Fig. 1a). We engineered a lentiviral system that co-encodes an shRNA and membrane-targeted mVenus (mem:Venus) fluorescent protein to transduced cells (Fig. 1b), and generated a highly validated library targeting all ARFs, GEFs, GAPs and 72 known interactors (Fig. 1c; Supplementary Table S1). Examination of ARF GTPase expression across 9 prostate cancer cell lines indicated that metastatic PC3 cancer cells showed high expression levels of all ARF GTPases compared to normal prostate cells (RWPE-1, PRECLH; Fig. S1a-h), particularly in 3D acini compared to 2D culture (Fig. S1g-j). PC3 cells also expressed almost all components of the ARFome (Fig. S1k). We therefore examined ARFome contribution to 3D morphogenesis in PC3 cells.

**Figure 1:**
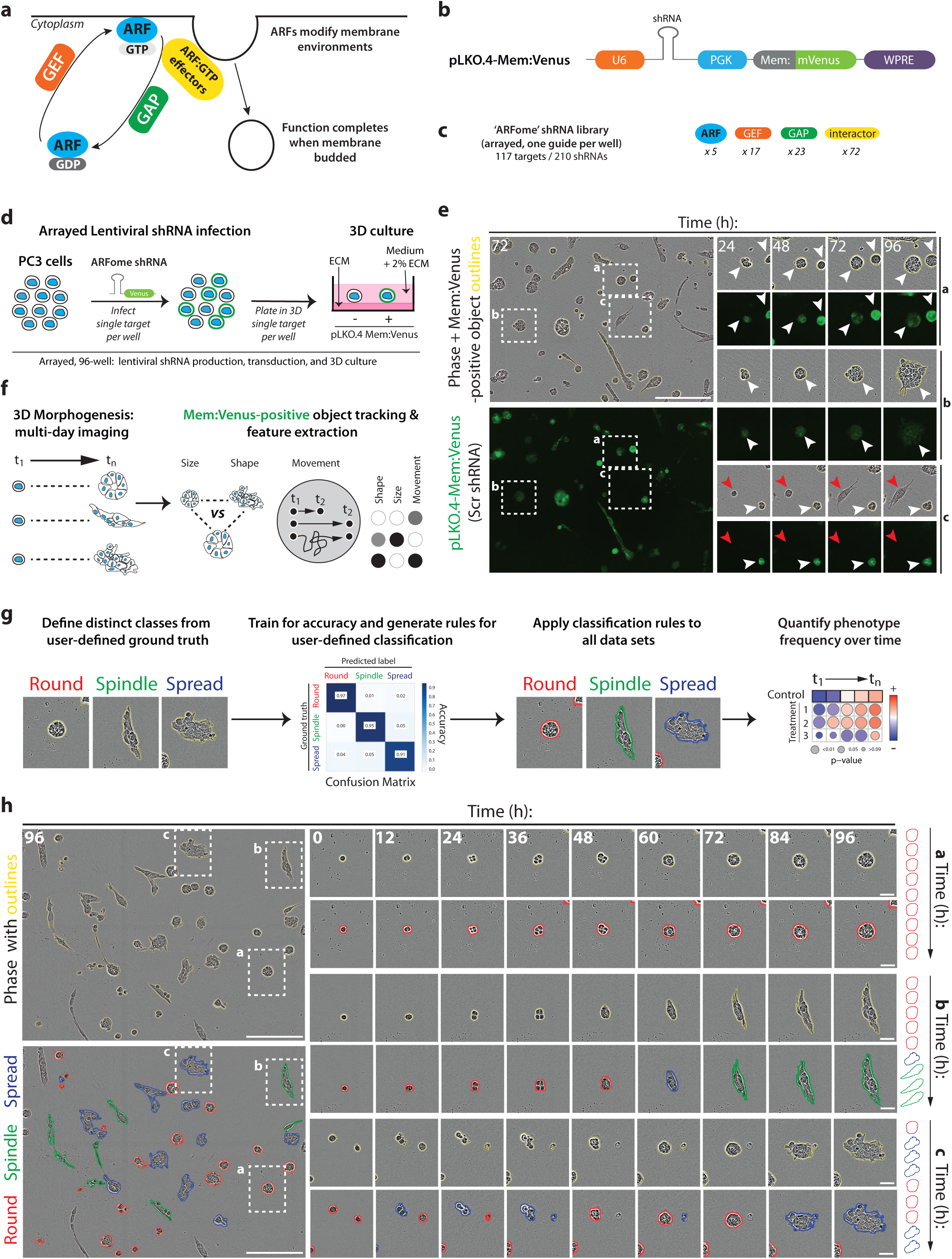
Development of a 3D functional genomic screen to examine ARF GTPase contribution to collective cancer cell behaviour. **a.** Cartoon, ARF GTPase cycle. **b.** Cartoon, pLKO.4-mem:Venus shRNA lentiviral vector. **c.** Schema, distribution of ‘ARFome’ components in shRNA library. **d.** Schema, 96-well based lentiviral infection of ‘ARFome’ shRNA into PC3 cells that were then cultured as heterogeneous 3D acini in extracellular matrix (ECM). **e.** Images of PC3 acini expressing pLKO.4-mem:Venus Scr shRNA. Yellow outlines, mem:Venus-positive acini. Magnified images of boxed regions (a-c) show acini at various times. White or red arrowheads, presence or absence of mem:Venus respectively. **f.** Schema, heterogeneous PC3 acini imaged over time vary in size, shape and movement characteristics. Properties measured and information extracted for thousands of mem:Venus-positive acini. **g.** Schema, phase images of 3 acini (yellow outlines) exhibiting morphological heterogeneity (Round, Spindle, Spread). Machine learning used to classify phenotypic states, train for accuracy and generate user-defined rules. Rules then applied to all data sets; Round, Spindle or Spread (red, green, blue outlines respectively) and changes in global state frequency tracked over time. **h.** PC3 acini (yellow outlines) and their user-defined classifications, Round, Spindle or Spread (red, green, blue outlines respectively). Magnified images of boxed regions (a-c) show classification of heterogeneous acini at various times. Cartoon, changes in user-defined classification over time.

We developed a high-throughput, arrayed, live imaging-based screening approach to determine the 3D phenotype of ARFome component depletion on multi-day morphogenesis. Control (Scramble shRNA-expressing, Scr) 3D acini could be distinguished from non-shRNA-expressing acini by the presence (Fig. 1d,e, white arrowheads) or absence (Fig. 1d,e, red arrowheads) of mem:Venus fluorescence, respectively. 3D acini were imaged every hour for 96 hours (Supplementary Movies S1, S2) and size, shape and movement features were extracted for thousands of mem:Venus-positive acini per manipulation (Fig. 1e,f; Supplementary Table S1). This live imaging approach revealed that multiple distinct 3D phenotypes occur in these cells in parallel, confirming our previous observations^3^. To detect these alternate phenotypes, we generated a machine learning classifier based on the Fast Gentle Boosting algorithm to define three morphogenesis classes with high fidelity to true user classification (91-97%): acini that are spherical (‘Round’), acini that are elongated (‘Spindle’), and those that are locally invading (‘Spread’), which were then applied to classify and quantify the phenotype of all acini (Fig. 1g). Application of these classes to timelapse sequences indicated that distinct phenotypes arise from single cells, and that cells can stay in the same cell state throughout observation or cycle between states to give rise to alternate phenotypes (Fig. 1h).

To identify the phenotypes of individual ARFome component depletion, we compared the relative fold-change in frequency of Round, Spindle and Spread phenotypes within each shRNA-expressing condition to control cells (Scr shRNA) over 96 hours of observation (Fig. 2a; Fig. S2a). The resulting relative change in each phenotype across time allowed division of shRNAs against ARFome components into 7 distinct Phenotype Groups based on clustering, including highly round (Group 3), increased local spreading (Group 1), or increased spindle-type behaviours (Group 2) (Fig. 2a,b; Fig. S2a). Application of these groupings to network analysis of ARFome interactions from literature and publicly available databases indicated phenotypic clusters centred around different ARF GTPases (Fig. 2c,d), which could not be easily appreciated based on connections between nodes alone due to the highly interconnected nature of the ARFome. ARF4 and ARF5 associated with Phenotype Groups 4 and 6 that are characterised by minimal change relative to control cells (Fig. 2c,d). While ARF6 was associated solely with Phenotype Group 1, both ARF1 and ARF3 had one shRNA in each of Phenotype Groups 5 and 1, which displayed a modest but robust increase in Spindle and Spread behaviours, respectively. We therefore subsequently focused on exploring how these two highly similar Class I ARFs contribute to 3D multicellular morphogenesis.

**Figure 2:**
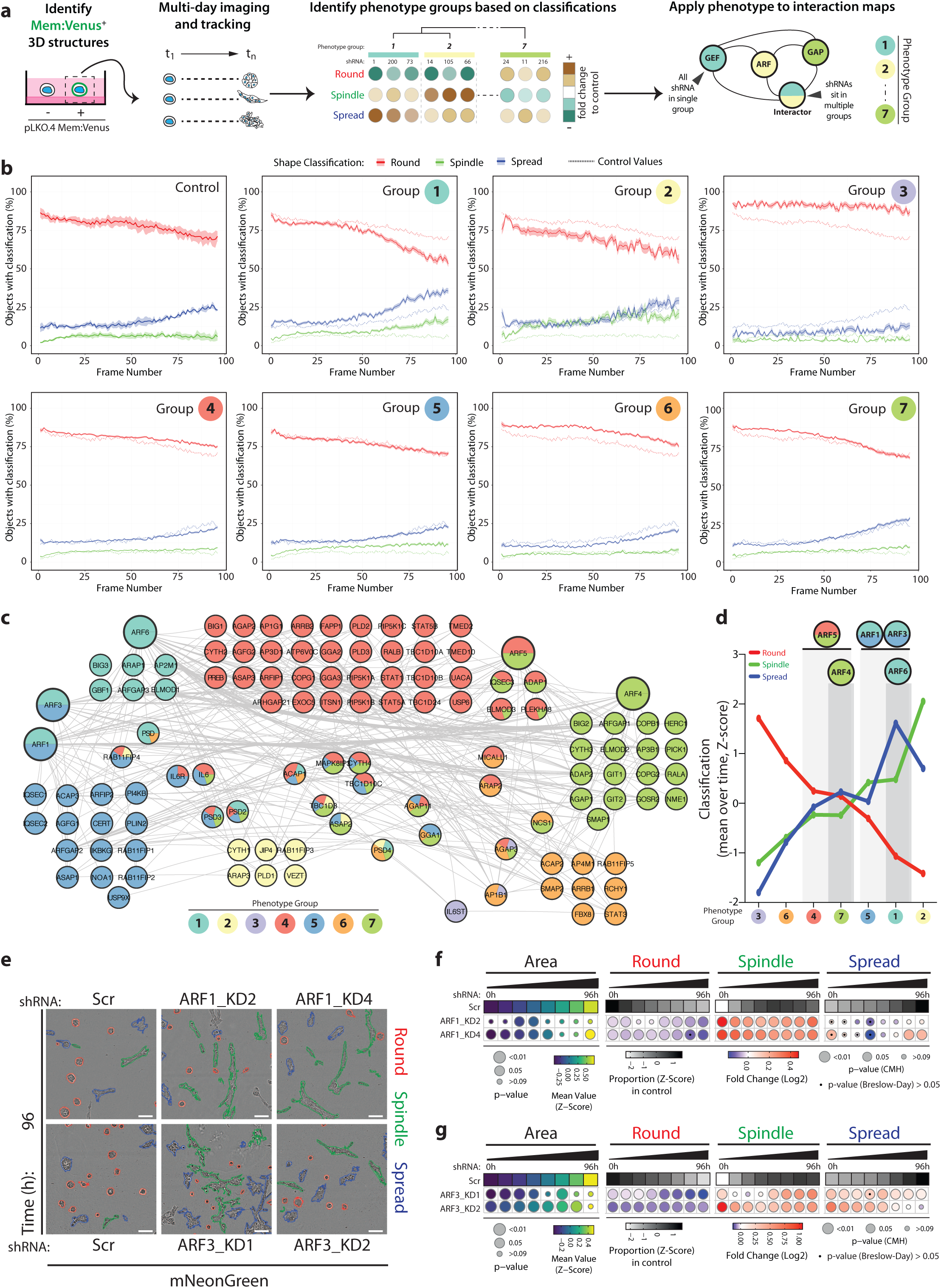
Contribution of each component of the ARFome to collective cancer cell behaviour was examined by individual depletion. **a.** Schema, PC3 acini expressing mem:Venus shRNAs were imaged, tracked and classified. Phenotype Group 1 – 7 identified based on frequency of classification into Round, Spindle and Spread over time. Interaction map shown, shRNAs colour coded by Phenotype Group. **b.** Graphs show percentage of acini classified as Round (red), Spindle (green) and Spread (blue) for each Phenotype Group and Scr shRNA (Control, on each graph). Data are mean, shaded regions represent s.e.m. Viral infections and live 3D spheroid assays carried out 3 independent times. Each experimental replicate consisted of 18 technical replicates of Scr (170,674 acini in total) and 1 replicate of 210 ARFome shRNAs (Supplementary Table S1). **c.** STRING network analysis of acini visualised using Cytoscape. Phenotype Groups 1 - 7 identified by frequency of acini classification into Round, Spindle and Spread. Colours indicate Phenotype Group and the proportion of shRNAs for each target that sit in different groups is shown. **d.** Graph is mean percentage of acini, across all timepoints, classified into Round, Spindle and Spread per Phenotype Group. **e.** PC3 acini expressing mNeonGreen (mNG) and Scr, *ARF1* or *ARF3* shRNA. Outlines; Round (red), Spindle (green) and Spread (blue). Scale bar, 100 μm n=4 and 6 experimental replicates for *ARF1* and *ARF3* shRNA respectively each with 4 technical replicates/condition. 16,760 (Scr), 21,086 (*ARF1_*KD2), 19,424 (*ARF1_*KD4) and 31,414 (Scr), 40,135 (*ARF3_*KD1), 30,460 (*ARF3_*KD2) acini quantified in total. **f-g.** Quantitation of **(e).** Heatmaps, Area is mean of Z-score normalised values (purple to yellow). P-values, Student’s t-test, Bonferroni adjustment, represented by size of bubble. Heatmaps, Round, Spindle or Spread is Log_2_ Fold Change from control (Scr) (blue to red). Proportion of control at each time is Z-score normalised (white to black). P-values, Cochran-Mantel-Haenszel (CMH) test, Bonferroni adjusted, represented by size of bubble. Dot indicates p-value (Breslow-Day test, Bonferroni-adjusted) for consistent effect magnitude.

We independently validated depletion of each Class I ARF GTPase using an orthogonal approach of lentiviral shRNA expression and stable antibiotic selection (Fig. S2b,c). Analysis of 3D phenotypes revealed that Area, as an indirect measure of growth, was unaffected by ARF1 or ARF3 depletion (Fig. 2e-g). ARF1 depletion induced Spindle-type behaviours in acini at the expense of Round phenotype (Fig. 2e,f). In contrast, ARF3 depletion induced Spindle behaviours at the expense of Round phenotype, but also induced Spread phenotypes (Fig. 2e,g), indicating that upon longer-term selection for stable depletion these highly similar ARFs do not share identical phenotypes upon depletion. Independent validation of multiple ARFome members displaying phenotypes resembling ARF1/3/6 from our screen revealed that a) the ARFGEF PSD/EFA6A, and b) the dual RAB11-GTP and ARF-GTP effector RAB11FIP4/Arfophilin-2 displayed phenotypes mirroring ARF3, not ARF1 (Fig. S2d-i). This suggests that RAB11FIP4 and PSD are key ARF3 interactors normally suppressing invasion. This indicates successful identification from our screen of potentially co-acting partnerships in the ARFome that regulate collective morphogenesis.

### Contextual regulation of shape and invasion by Class I ARF GTPases

We examined Class I ARF contribution to a range of cellular behaviours. Depletion of neither ARF1 nor ARF3 affected cell proliferation in 2D or 3D culture (Fig. S3a-e), corroborating a lack of effect on Area measurements in 3D culture (Fig. 2f,g). The effect of Class I ARFs on individual cell shape in 2D culture was variably aligned with their respective collective 3D phenotypes upon ARF depletion (Fig. S3e-h). ARF1 depletion increased the frequency of Spindle shape of single cells in 2D (Fig. S3e-g), mirroring the collective Spindle phenotype induced in 3D upon ARF1 depletion (Fig. 2e-f). In contrast, despite inducing both Spindle and Spread collective 3D behaviours, ARF3 depletion in 2D culture robustly induced Round single cell shape (Fig. S3e,f,h). This was mirrored in 2D culture by PSD depletion (Fig. S3i), but not RAB11FIP4 depletion (Fig. S3j). Therefore, the effect of ARF3 on collective morphogenesis is specific to a 3D environment, not single cells. This emphasises the requirement to examine ARF function in 3D systems that allow assessment of collective behaviours.

To determine the effect of Class I ARFs on collective behaviours, we examined the ability of wounded monolayers to invade, which can occur through wound repair via single cell invasion, movement as a sheet, or as a leader cell-led chain of cells (Fig. 3a, Supplementary Movies S3-S5). In the absence of exogenous ECM addition, this approach assays 2D migration. Plating of monolayers onto ECM and overlay of cells and wound with further ECM allows examination of collective invasion. Despite their differences in single cell shape effects, depletion of either ARF1 or ARF3 increased 2D migration ability, largely through the movement of single cells (Fig. 3b,c). In 3D invasion contexts, either ARF1 or ARF3 depletion resulted in chain-type invasion mechanisms (Fig. 3d,e; arrowheads), mirroring the induction of Spindle chains in 3D acinus culture in both conditions (Fig 2e-g). Co-depletion of ARF1 and ARF3 induced increased Spindle and Spread behaviours in both 2D and 3D culture and increased spindle-type invasion from wounded monolayers (Fig. 3f-j), resulting in a phenotype midway between ARF1 and ARF3 individual depletion. Depletion of PSD and RAB11FIP4, also induced an increase in spindle-type monolayer invasion, particularly for RAB11FIP4 (Fig. S3k,l). These data indicate individual and key roles for each of the Class I ARFs in suppressing invasion in cells and emphasise that the phenotype of ARF depletion is contextual on whether cells are assayed individually or collectively. This suggests that a function of Class I ARFs may be to regulate molecules that control collective behaviours.

**Figure 3:**
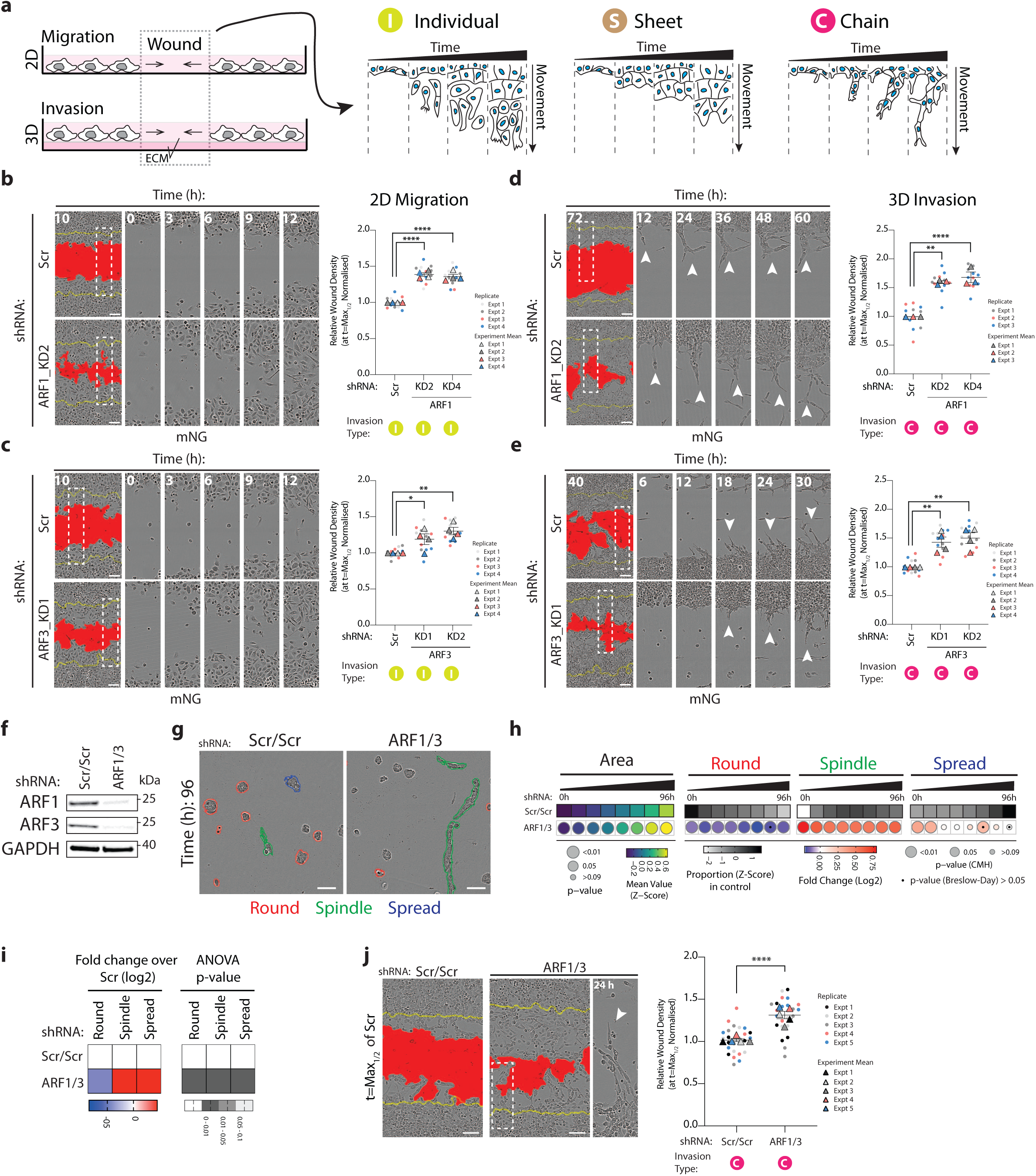
Class I ARF GTPases regulate migration and invasion of prostate cancer cells. **a.** Schema, 2D migration or 3D invasion (+ ECM) of wounded PC3 monolayer. Three modes of movement observed, cells moving individually (I), as a sheet (S) or as chains (C). **b-c.** Phase images of cells expressing mNG and Scr or **(b)** *ARF1* or **(c)** *ARF3* shRNA in 2D migration assay. Yellow lines, initial wound and red pseudo colour, wound at t=Max_1/2_. Scale bars, 100 μm. Magnified images of boxed regions at different times shown. Graph is Relative Wound Density (RWD) at t=Max_1/2_ (Scr = 50% closed). Data is mean ± s.e.m. (4 experimental replicates, triangles, 2-4 technical replicates, circles). P-values (Student’s 2-tailed t-test), *p ≤0.05, **p≤0.01 and ****p≤0.0001. **d-e.** Cells in a wounded monolayer overlaid with 25% ECM were imaged to observe 3D invasion. Phase images of cells expressing mNG and Scr, **(d)** *ARF1* or **(e)** *ARF3* shRNA shown. Yellow lines, initial wound and red pseudo colour, wound at t=Max_1/2._ Scale bars, 100 μm. Magnified phase images of boxed regions at different times shown. White arrowheads, invasive chains. Graph is RWD at t=Max_1/2_, normalised to Scr. Data is mean ± s.e.m. (3-4 experimental replicates, triangles, 3-5 technical replicates, circles). P-values (Student’s 2-tailed t-test), **p≤0.01 and ****p≤0.0001 **f.** Western blot analysis of PC3 cells expressing Scr/Scr or *ARF1/3* shRNA for ARF1 or ARF3. GAPDH is loading control for ARF3 and sample control for ARF1. Panels shown are representative of 3 independent lysate preparations. **g.** Phase images of acini expressing Scr/Scr or *ARF1/3* shRNA. Outlines; Round (red), Spindle (green) and Spread (blue). Scale bar, 100 μm n=5 experimental replicates each with 3-4 technical replicates/condition. 20,645 (Scr/Scr), 8,601 (*ARF1/3*) mem:Venus-positive acini quantified in total. **h.** Quantitation of **(g)**. Heatmaps, Area is mean of Z-score normalised values (purple to yellow). P-values, Student’s t-test, Bonferroni adjustment, represented by size of bubble. Heatmaps, Round, Spindle or Spread is Log_2_ Fold Change from control (Scr/Scr) (blue to red). Proportion of control at each time is also Z-score normalised (white to black). P-values, Cochran-Mantel-Haenszel (CMH) test, Bonferroni adjusted, represented by size of bubble. Dot indicates p-value (Breslow-Day test, Bonferroni-adjusted) for consistent effect magnitude. **i.** 2D PC3 cells expressing Scr/Scr or *ARF1/3* shRNA classified into Round, Spindle and Spread. Heatmaps, Log_2_ Fold Change over Scr/Scr. P-values, one-way ANOVA, greyscale values as indicated. n=2 experimental replicates with 4 technical replicates/condition. 3,323 (Scr/Scr) and 1,847 (*ARF1/3*) mem:Venus-positive cells quantified in total. **j.** Phase images of cells expressing Scr/Scr or *ARF1/3* shRNA in 3D invasion assay shown. Yellow lines, initial wound and red pseudo colour, wound at t=Max_1/2_. Scale bars, 100 μm. Magnified image of boxed region shown. White arrowhead, invasive chain. Graph is RWD at t=Max_1/2_, normalised to Scr/Scr. Data is mean ± s.e.m. (5 experimental replicates, triangles, 4-5 technical replicates, circles). P-values (Student’s 2-tailed t-test), ****p ≤ 0.0001.

### ARF3 is a rheostat for the modality of collective invasion

Given the induction of Class I ARF expression in 3D culture versus 2D culture (Fig. S1i,j), we examined whether Class I ARFs may be rheostats for regulation of invasive activity. Overexpression of mNeonGreen (mNG)-tagged ARF1 (ARF1-mNG) or ARF3 (ARF3-mNG), both of which localised to intracellular puncta compared to cytoplasmic mNG alone, did not affect cell growth in either 2D or 3D contexts (Fig. S4a-e), similar to depletion of these ARFs. The shape of 2D single cells was modulated by ARF1-mNG or ARF3-mNG in the converse fashion to depletion of each ARF: ARF1-mNG overexpression increased the Round single cell phenotype, while ARF3-mNG induced single cells to undergo spreading (compare Fig. S4f to S3g,h). This confirms distinct effects of ARF1 and ARF3 on 2D cell shape.

When examining the effects on cell movement we observed that while ARF1- mNG expression had no effect on 2D migration (Fig. S4g,h) or 3D invasion (Fig. 4a,b; white arrowheads demarcating chain-led invasion), ARF3 drastically affected cell behaviours. ARF3-mNG overexpression increased both 2D wound closure and 3D invasion, but did so by inducing sheet-like movement of the cell monolayer (Fig. 4c,d; black arrowheads denoting sheet movement; Fig. S4i,j). In 3D acinus culture, ARF1-mNG overexpression displayed largely no phenotypic alteration (Fig. 4e; S4a). In contrast, ARF3-mNG overexpression induced Spindle phenotypes at early timepoints that decreased over time, relative to control, while cell spreading was robustly increased at all timepoints (Fig. 4f; S4a), mirroring the sheet like invasion of monolayers (Fig. 4c; black arrowheads). This indicates a unique function of ARF3 as a rheostat that controls the modality of collective invasion; low ARF3 levels result in leader cell-led chain-type invasion, while elevated ARF3 levels switch cells to a collective sheet-movement invasive activity. It is important to note that these phenotypes manifest in 3D culture where collective activity is assayed for.

**Figure 4:**
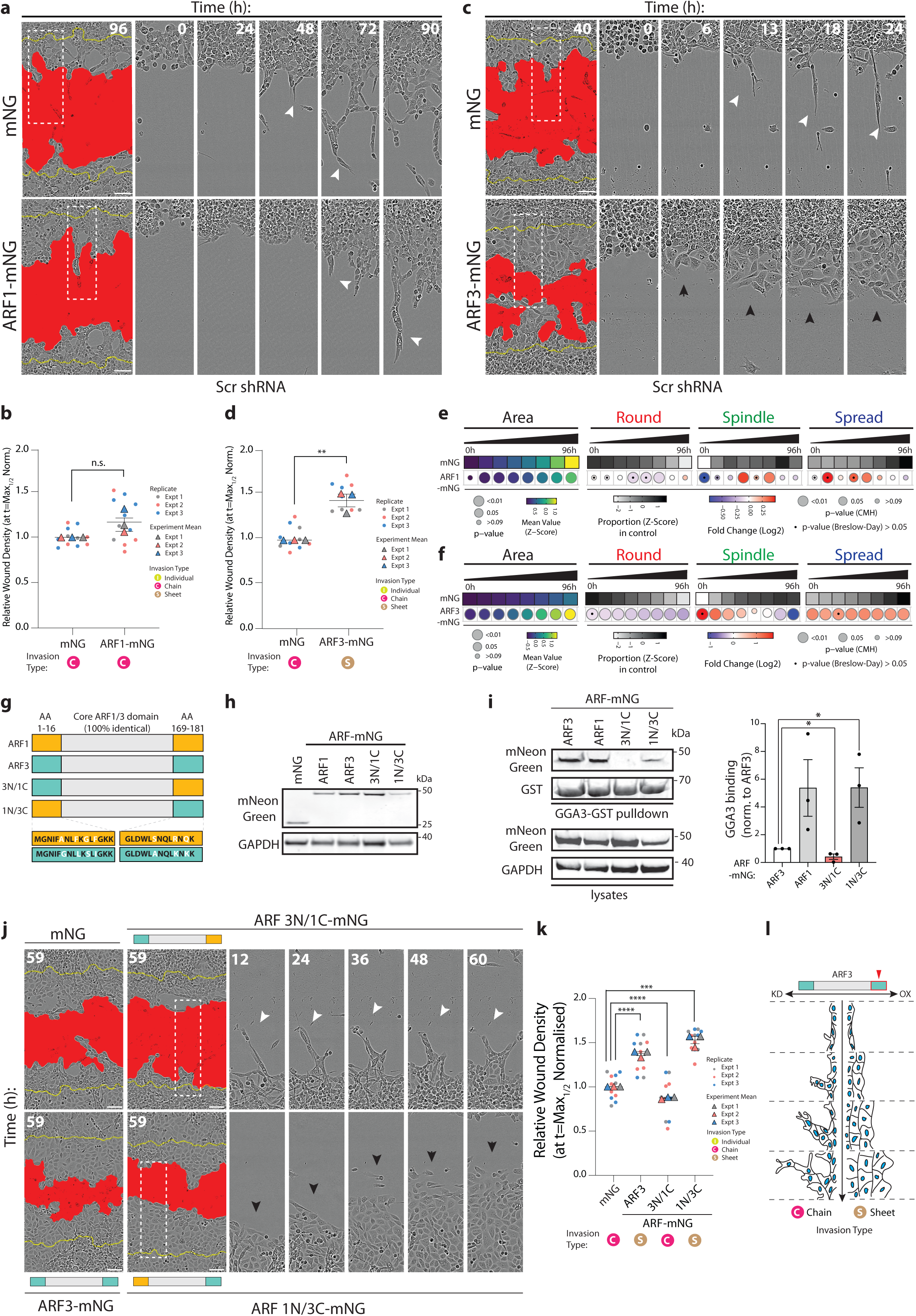
ARF3 is a rheostat for the modality of collective invasion. **a-d.** PC3 cells expressing mNG, **(a)** ARF1-mNG or **(c)** ARF3-mNG and Scr shRNA in 3D invasion assay. Yellow lines, initial wound and red pseudo colour, wound at t=Max_1/2_. Scale bars, 100μm. Magnified images of boxed regions shown. White and black arrowheads, invasive chain or sheet respectively. RWD at t=Max_1/2_, normalised to mNG is shown in graphs **(b, d).** Data is mean ± s.e.m. (3 experimental replicates, triangles, 2-5 technical replicates, circles. P-values (Student’s 2-tailed t-test), n.s. not significant and **p≤0.01. **e-f.** PC3 acini expressing mNG, **(e)** ARF1-mNG or **(f)** ARF3-mNG and Scr shRNA were classified into Round, Spindle and Spread. n=6 and 4 experimental replicates for ARF1-mNG and ARF3-mNG respectively each with 2-4 technical replicates/condition. **(e)** 5,005 (mNG), 1,938 (ARF1-mNG) and **(f)** 9,320 (mNG), 8,699 (ARF3-mNG) mNG-positive acini quantified in total. Heatmaps, Area is mean of Z-score normalised values (purple to yellow). P-values, Student’s t-test, Bonferroni adjustment, represented by size of bubble. Heatmaps, Round, Spindle or Spread is Log_2_ Fold Change from control (mNG) (blue to red). Proportion of control at each time is Z-score normalised (white to black). P-values, Cochran-Mantel-Haenszel (CMH) test, Bonferroni adjusted, represented by size of bubble. Dot indicates p-value (Breslow-Day test, Bonferroni-adjusted) for consistent effect magnitude. **g.** Schema, Class 1 ARFs share 100% identical core region but differ in 7 amino acids (AA) in N and C termini. ARF chimeras with ARF3 N-terminal and ARF1 C-terminal (3N/1C) and ARF3 C-terminal and ARF1 N-terminal (1N/3C) created. **h.** Western blot of PC3 cells expressing mNG, ARF1-mNG, ARF3-mNG and ARF-mNG chimeras for mNeonGreen and GAPDH, as loading control. Panels shown are representative of 3 independent lysate preparations. **i.** ARF-GTP pulldown and representative western blot for mNeonGreen, GST and GAPDH, as loading control for mNeonGreen. n=3 independent lysate preparations and pulldowns. Graphs show mean GGA3 binding ± s.e.m. normalised to ARF3. P-values (Student’s 2-tailed t-test), *p ≤0.05. **j-k.** PC3 cells expressing mNG, ARF3-mNG or ARF-mNG chimeras plated in 3D invasion assay. Yellow lines, initial wound and red pseudo colour, wound at t=Max_1/2_. Scale bars, 100μm. Magnified images of boxed regions shown. White and black arrowheads, invasive chain or sheet respectively. RWD at t=Max_1/2_, normalised to mNG shown in **(k).** Data is mean ± s.e.m. (3 experimental replicates, triangles, 2-5 technical replicates, circles). P-values (Student’s 2-tailed t-test), ***p ≤ 0.001 and ****p ≤ 0.0001. **l.** Schema, ARF3 expression levels affect mode of invasion.

We mapped the unique ability of ARF3 to induce a collective sheet-type invasion phenotype by creating chimeras between ARF1 and ARF3, which only differ by 4 amino acids in their N-termini and 3 amino acids in their C-termini (Fig. 4g). This revealed that the C-terminus of ARF3 is required for sheet-type invasive activity. Examination of GTP-loading of ARF1, ARF3, and chimeras revealed that both ARF1-mNG and ARF3-mNG are GTP-loaded, with ARF1 displaying increased GTP levels compared to ARF3 (Fig. 4g-i) and indicating that a lack of effect of ARF1 overexpression is not simply due to lack of GTP-loading of the tagged ARF1. ARF chimera with an ARF3 N-terminus and ARF1 C-terminus (3N/1C) displayed poor GTP loading despite robust expression, while the converse ARF chimera with ARF1 N-terminus and ARF3 C-terminus (1N/3C) showed increased GTP-loading compared to ARF3 alone (Fig. 4g-i). Compared to the cytoplasmic and nuclear fluorescence of mNG alone, ARF3-mNG localised to intracellular puncta in 2D single cells and 3D acini (Fig. S4k; white arrows). ARF 3N/1C-mNG chimera abrogated the ARF3- induced increase in Spread phenotype observed in 2D (Fig S4l) and resulted in clustering of fluorescent puncta towards the cell periphery in 2D single cells (Fig. S4k,m; black arrows), and near cell-cell contacts in 3D acini (Fig. S4k). The ARF 1N/3C-mNG chimera, in contrast, displayed enlarged puncta that were nonetheless reminiscent of the distribution of ARF3 in 2D single cells and 3D acini (Fig. S4k,m). Phenotypically, the ARF3 N-terminus was dispensable, and C-terminus indispensable, to maintain sheet-type invasion (Fig 4j,k), and Spread-type acinus formation (Fig. S4k,n) to levels reminiscent of ARF3 wild-type. These data indicate that ARF3 acts as a rheostat for the modality of invasion and that this function is dictated to the Class I ARFs by three unique residues in the ARF3 C-terminus (A174/K178/K180) (Fig. 4l).

### N-cadherin is a key interactor of ARF3 that controls morphogenesis

As ARF3 levels controlled the modality of collective movement in 3D, we examined whether ARF3 also contributed to junctional organisation between cells. Compared to mNG-expressing acini alone, ARF3-depleted acini displayed lowered overall F-actin intensity and a robust decrease of F-actin specifically at cell-cell, but not cell-ECM, junctions (Fig. 5a, arrowheads; Fig. S5a,b). In contrast, ARF3-mNG overexpression resulted in increased overall F-actin intensity, which was observed at the cell cortex (Fig. 5a, arrows; Fig. S5a,b). Analysis of the major cell-cell adhesion molecules E-cadherin and N-cadherin, which are co-expressed in PC3 cells, revealed that ARF3 levels associated with altered N-cadherin, but not E-cadherin, levels; ARF3 depletion decreased N-cadherin levels, while ARF3-mNG overexpression strongly increased N-cadherin levels (Fig. 5b-d). Moreover, N-cadherin co-localised with a sub-set of ARF3-positive intracellular puncta in 3D acini (Fig. 5e) and N-cadherin could be recovered in ARF3 immunoprecipitants (Fig. 5f). ARF3 has been reported as part of the N-cadherin interactome^9^ and our data indicate that N-cadherin is regulated by ARF3 in our system.

**Figure 5:**
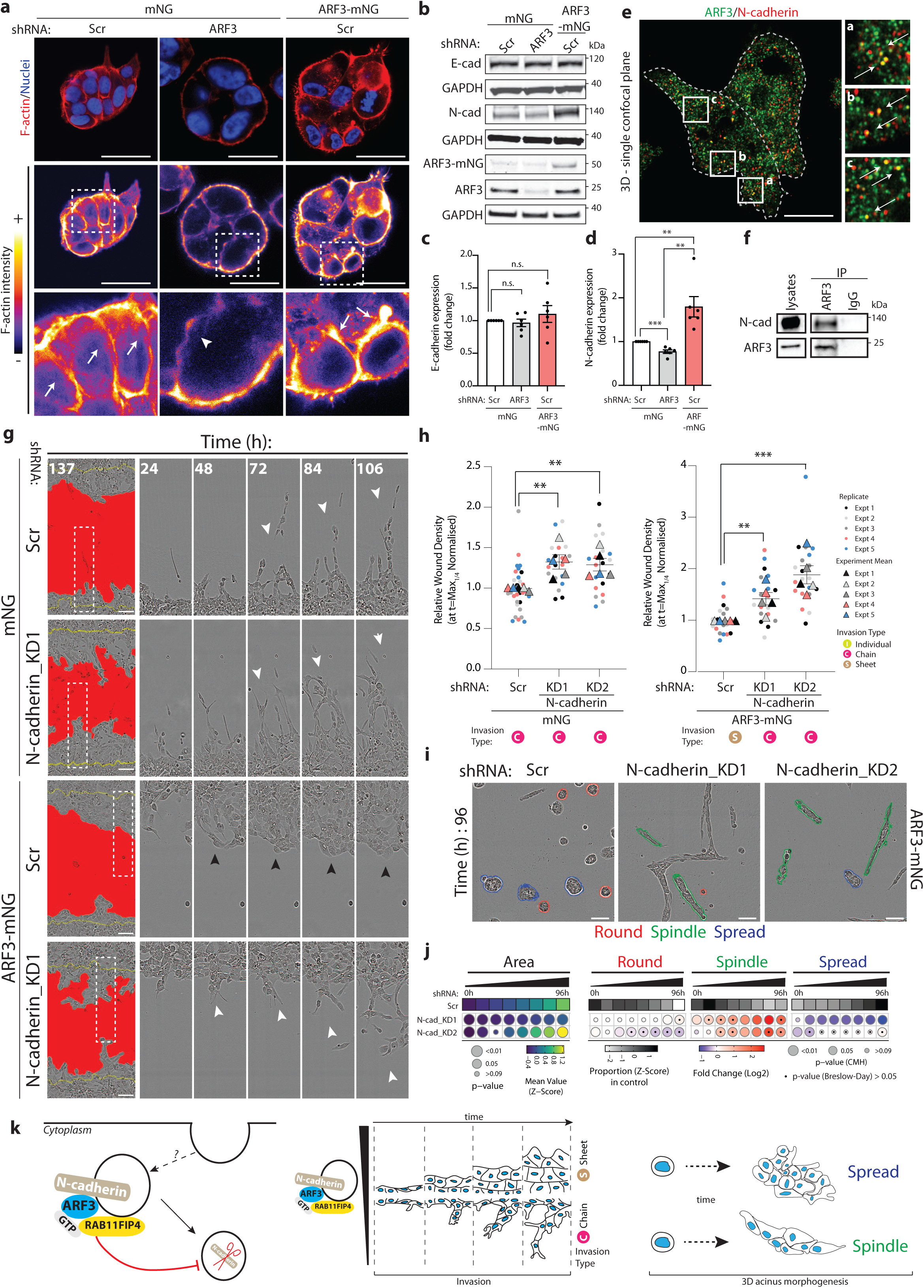
N-cadherin is a key interactor of ARF3 that controls morphogenesis. **a.** Confocal images of PC3 acini expressing mNG or ARF3-mNG and either Scr or ARF3 shRNA stained with F-actin (red) and Hoechst (nuclei, blue). F-actin intensity, FIRE LUT. Magnified images of boxed regions shown. White arrows or arrowheads, presence or absence of intense F-actin staining in junctions respectively. Scale bars, 20 μm. Images representative of phenotypes observed in 3 experimental replicates. **b-d.** Representative western blot of PC3 cells **(b)** expressing mNG or ARF3-mNG and Scr or *ARF3* shRNA for E-cadherin, N-cadherin and ARF3. GAPDH loading control is shown for each blot. n=6 independent lysate preparations. Data is presented in **(c,d)** as mean fold change ± s.e.m. normalized to Scr. P-values (Student’s 2-tailed t-test), n.s. not significant and **p≤0.01. **e.** Confocal image of PC3 acini stained for ARF3 (green) and N-cadherin (red). Magnified images of boxed regions shown (a-c). White arrows, co-localization in subset of puncta. Scale bars, 20μm. Image is representative of co-localization observed in cells in 3 experimental replicates. **f.** Immunoprecipitation (IP) was performed using an anti-ARF3 antibody or mouse IgG and samples immunoblotted for N-cadherin and ARF3. Panels shown are representative of 3 IPs from 3 independent lysate preparations. **g-h.** PC3 cells in 3D invasion assay. Yellow lines, initial wound and red pseudo colour, wound at t=Max_1/4._ Scale bars, 100μm. Magnified images of boxed regions shown. White and black arrowheads, invasive chain or sheet respectively. RWD at t=Max_1/4_, normalised to Scr is shown in graphs **(h).** Data is mean ± s.e.m. (5 experimental replicates, triangles, 3-8 technical replicates, circles. P-values (Student’s 2-tailed t-test), **p≤0.01 and ***p≤0.001. **i.** Phase images of PC3 acini expressing ARF3-mNG and Scr or *N-cadherin* shRNA. Outlines; Round (red), Spindle (green) and Spread (blue). Scale bar, 100 μm n=3 experimental replicates each with 3 technical replicates/condition. 4,039 (Scr), 4,814 (*N-cadherin_*KD1) and 3,454 (*N-cadherin_*KD2) mNG acini quantified in total. **j.** Quantitation of **(i)**. Heatmaps, Area is mean of Z-score normalised values (purple to yellow). P-values, Student’s t-test, Bonferroni adjustment, represented by size of bubble. Heatmaps, Round, Spindle or Spread is Log_2_ Fold Change from control (Scr) (blue to red). Proportion of control at each time is Z-score normalised (white to black). P-values, Cochran-Mantel-Haenszel (CMH) test, Bonferroni adjusted, represented by size of bubble. Dot indicates p-value (Breslow-Day test, Bonferroni-adjusted) for consistent effect magnitude. **k.** Model, ARF3-RAB11FIP4 complex interacts with and regulates levels of N-cadherin in intracellular vesicles to control different modes of invasion.

ARF3 and N-cadherin acted to mutually stabilise each other’s level; N-cadherin levels decreased or increased upon ARF3 depletion or overexpression, respectively (Fig. 5b,d), while depletion of N-cadherin also decreased endogenous ARF3 levels (Fig. S5c-f). In contrast, E-cadherin levels were not consistently changed upon alteration of either ARF3 or N-cadherin (Fig. 5b,c; Fig. S5c). N-cadherin was essential for the switch between chain-type and sheet-type invasion, as depletion of N-cadherin increased 2D Spindle shape, chain-type invasion of 3D monolayers expressing mNG alone and completely reversed the sheet-type invasion of ARF3-mNG-expressing monolayers (Fig. S5g; Fig. 5g,h). This was not simply due to a decrease in ARF3 levels upon N-cadherin depletion, as total levels of ARF3 were maintained upon ARF3-mNG expression in N-cadherin knockdown cells (Fig. S5d,f). Moreover, N-cadherin depletion in control mNG-expressing cells phenocopied ARF3 depletion in 3D acini phenotypes (adoption of Spindle and Spread phenotypes; compare Fig. S5h,i to Fig. 2e,g), and completely reversed the spread-type phenotype of ARF3-mNG-expressing 3D acini (compare Fig. 4f to Fig 5i,j). This effect on N-cadherin levels was also mirrored by depletion of the ARF3 effector RAB11FIP4 (Fig. S5j). Taken together, these data indicate that ARF3, possibly in concert with the RAB11FIP4 regulator of endosomal recycling, act as a rheostat to control cellular levels of N-cadherin to influence the modality of invasion (Fig. 5k).

### ARF3 regulates metastasis in vivo

We examined the function of ARF3 in tumourigenesis *in vivo* through orthotopic intraprostatic xenograft of PC3 cells in control (mNG and Scr shRNA), ARF3- depleted (mNG and ARF3 shRNA) or ARF3-elevated (ARF3-mNG and Scr shRNA) conditions (Fig. 6a). Mice were examined at timed endpoint of 8 weeks, which allows for examination of effects on both primary tumour formation and metastasis^3^. No difference in cell engraftment or prostate weights at timed endpoint were detected between any conditions (Fig. 6b,c) suggesting no effect on primary tumour growth, similar to a lack of effect on 2D or 3D proliferation *in vitro* upon ARF3 manipulation (Fig. S3a-d; S4b-e). Mirroring the effect on switching collective movement modality *in vitro*, ARF3 depletion versus overexpression showed robust and alternate effects on metastasis *in vivo*. ARF3 depletion induced a fully penetrant metastatic incidence (the number of mice with a primary tumour that also possessed at least one metastasis) compared to a reduction in metastatic incidence in ARF3-overexpressing cells (100% in ARF3 knockdown, 67% in ARF3 overexpression, compared to 75% in control; Fig. 6d, p=0.0308). Moreover, ARF3 depletion also increased the number of organs with metastasis per mouse compared to ARF3 overexpression (Fig. 6e, p=0.01), as well as expanded the metastatic tropism to all organs examined, bar the stomach, while ARF3 overexpression resulted in metastasis to only very proximal organs (lumbar lymph nodes, mesentery, and spleen) (Fig. 6f). These data reveal that ARF3 is an *in vivo* regulator of metastasis, not primary tumour formation. Moreover, this suggests that of the alternate modalities of collective movement that ARF3 can influence *in vitro*, while sheet-type movement conditions may allow local metastasis, only the spindle-type chain-based invasive modality is able to induce distant and widespread metastasis.

**Figure 6:**
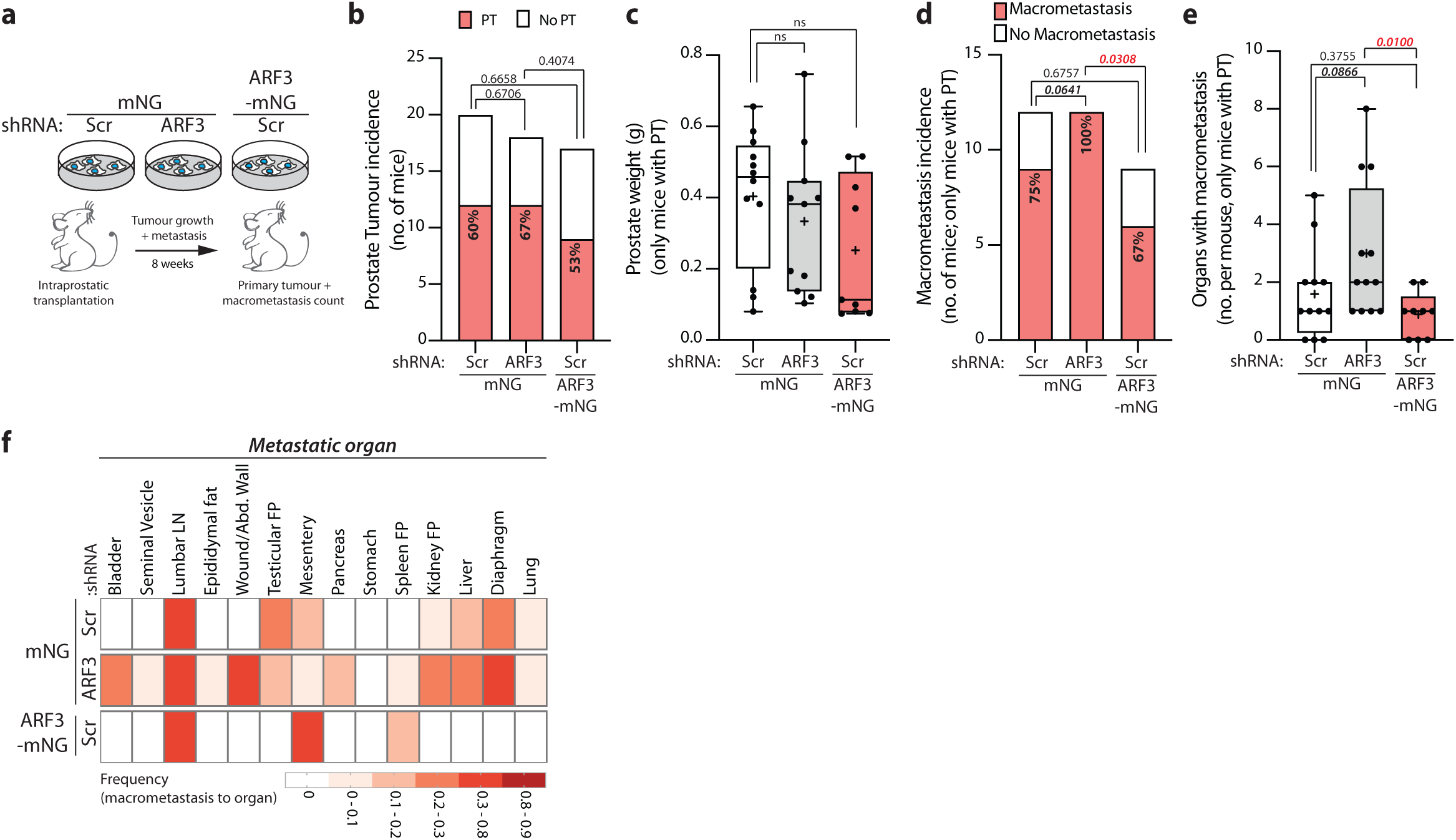
ARF3 regulates metastasis in vivo. **a.** Schema, intraprostatic transplantation of PC3 cells expressing mNG or ARF3-mNG and Scr or *ARF3* shRNA into CD-1 nude male mice. Primary tumour (PT) formation and incidence and location of macrometastasis determined after 8 weeks. **b.** PT incidence in mice (total number) transplanted with PC3 cells expressing mNG, Scr shRNA (control, 20 mice), mNG, *ARF3* shRNA (18 mice) or ARF3-mNG, Scr shRNA (17 mice). P-values (Chi-squared test), indicated. **c.** Prostate weight in mice (with PT only) transplanted with PC3 cells expressing mNG, Scr shRNA (control, 12 mice), mNG, *ARF3* shRNA (12 mice, (weight of 1 mouse prostate not recorded)) or ARF3-mNG, Scr shRNA (9 mice). Box and whiskers plot, min–max percentile; +, mean; dots, outliers; midline, median; boundaries, quartiles. P-values (Student’s 2-tailed t-test), n.s. not significant. **d.** Macrometastasis incidence in mice (with PT only) transplanted with PC3 cells expressing mNG, Scr shRNA (control, 12 mice), mNG, *ARF3* shRNA (12 mice) or ARF3-mNG, Scr shRNA (9 mice). P-values (Chi-squared test), indicated. **e.** Macrometastasis count/mouse in mice (with PTs only) transplanted with PC3 cells expressing mNG, Scr shRNA (control, 12 mice), mNG, *ARF3* shRNA (12 mice) or ARF3-mNG, Scr shRNA (9 mice). Box and whiskers plot, min–max percentile; +, mean; dots, outliers; midline, median; boundaries, quartiles. P-values (Mann-Whitney test (2-tailed)), indicated on graph. **f.** Macrometastasis frequency to indicated organs for mice (with PT only) transplanted with PC3 cells expressing mNG, Scr shRNA (control, 12 mice), mNG, *ARF3* shRNA (12 mice) or ARF3-mNG, Scr shRNA (9 mice).

### N-cadherin and ARF3 expression identify poor-outcome prostate cancer patients

As the ARF3-N-cadherin module controls invasion modality *in vitro* and metastasis *in vivo*, we examined whether levels of ARF3 and/or N-cadherin may identify patients with poor outcome or metastatic disease. *ARF3* mRNA levels across prostate normal tissue, primary tumour or metastasis displayed inconsistent trends across independent patient cohorts (Fig. 7a-e). Examination of two larger patient datasets indicated that while no difference in *ARF3* mRNA levels was observed in The Cancer Genome Atlas (TCGA) between normal prostate tissue versus primary tumour (PRAD, Prostate Adenocarcinoma; normal, n=52; tumour, n=497; Fig. 7f; Fig. S6a) or across Gleason Grade (Fig. 7g), a modest but significant decrease in *ARF3* mRNA level was observed in tumours when a number of independent datasets were combined (GENT2 database^10^) (Prostate; normal, n=86; tumour, n=323; Fig. S6b). Across tumour types with sufficient normal and tumour tissue for comparison, in the TCGA dataset (22 tumour types) both colon adenocarcinoma (COAD) and kidney renal clear cell carcinoma (KIRC) displayed decreased *ARF3* levels in the tumour, while nine tumour types displayed higher *ARF3* mRNA levels (Fig. S6a). In the combined datasets, colon and kidney again displayed lower tumour *ARF3* levels, as did 4 additional tumour types (blood, brain, prostate, and skin; Fig. S6b). In contrast, compared to normal tissue, increased expression of *ARF3* was observed in tumours in 9/22 (TCGA; Fig. S6a) or 12/30 (combined datasets; Fig S6b) tumour types. This indicates that while *ARF3* levels are widely altered in tumour versus normal tissue, the directionality of alternation of *ARF3* levels in tumours is dependent on tissue type and that *ARF3* expression alone is not a consistent indicator of clinical characteristics.

**Figure 7:**
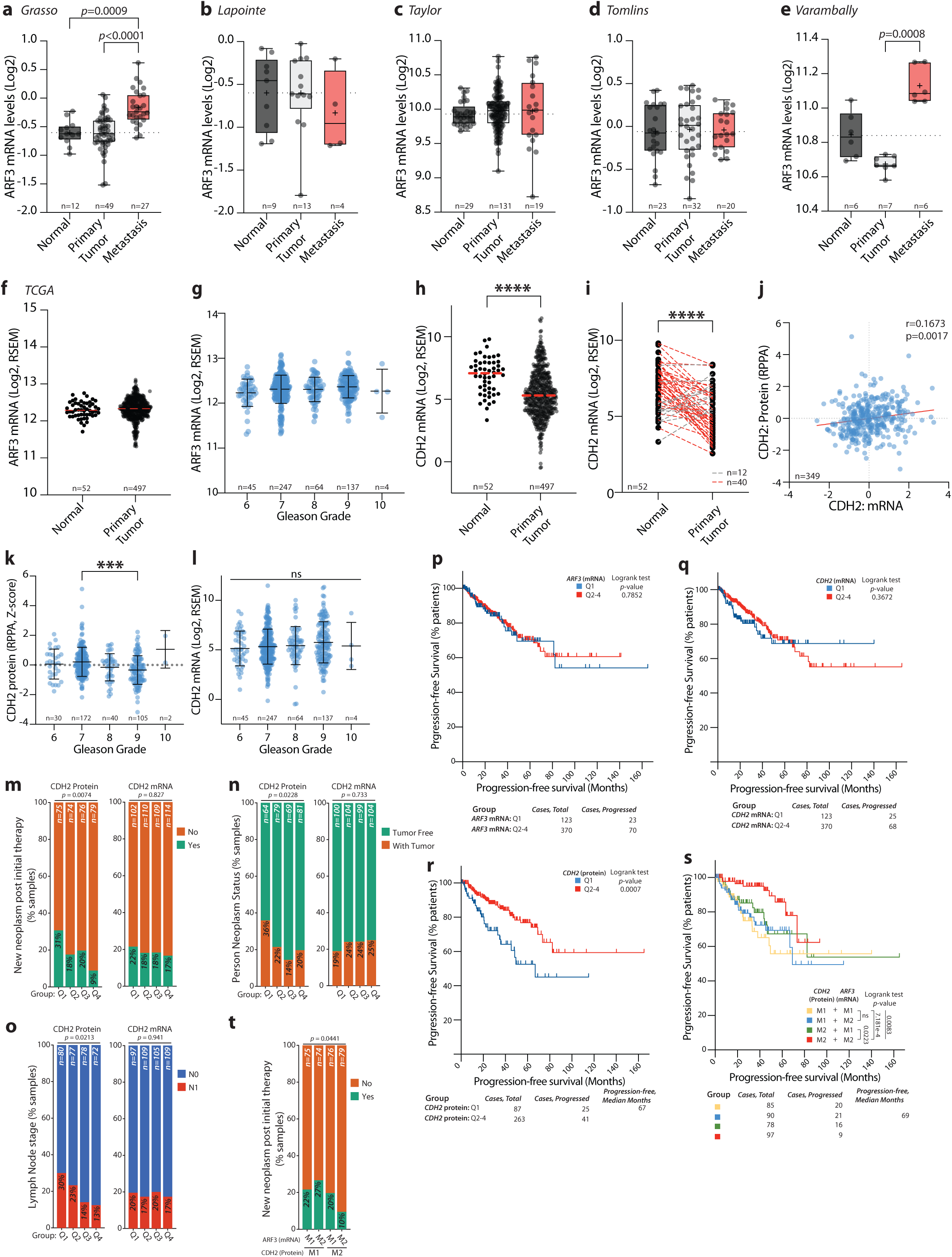
N-cadherin and ARF3 expression identify poor-outcome prostate cancer patients. **a-e.** *ARF3* mRNA expression in normal prostate, primary tumour and metastasis samples from prostate cancer patients. Patient numbers, on graph. Glinsky^37^, Grasso (GSE35988), Lapointe (GSE3933), Taylor (GSE211032), TCGA, Tomlins (GSE6099), Varambally (GSE3325). P-values, Kruskal-Wallis test with Dunn’s multiple comparison test. **f.** *ARF3* mRNA (Log2, RSEM) from Normal vs Primary Tumour prostate samples (dataset, TCGA). Patient numbers, on graph. P-values, two-tailed unpaired t-test with Welch’s correction. **g.** *ARF3* mRNA (Log2, RSEM) from prostate tumour samples with different Gleason Grade scores (dataset, TCGA). Patient numbers, on graph. P-values, one-way ANOVA with Tukey corrections for multiple testing. **h-i.** *CDH2* mRNA (Log2, RSEM) from Normal vs Primary Tumour prostate samples (dataset, TCGA). Patient numbers, on graph. (**h)**, all samples; (**i)**, matched samples from same patients. Lines, directionality of change in normal compared to tumour; red, elevated in tumour; grey, elevated in normal. P-values, two-tailed unpaired t-test with Welch’s correction, ****p≤0.0001. **j.** Correlation (Pearson) between *CDH2* mRNA and N-cadherin protein in prostate tumours (dataset, TCGA). **k-l.** CDH2/N-cadherin protein **(k)** or mRNA levels (Log2, RSEM) **(l)** from prostate tumour samples with different Gleason Grade scores (dataset, TCGA). Patient numbers, on graph. P-value, one-way ANOVA with Tukey corrections for multiple testing, n.s. not significant and ***p≤0.001. **m-o.** Clinical parameters upon grouping of patients based on quartiles of expression (Q1-4) for N-cadherin protein or *CDH2* mRNA in prostate tumour samples. **(m)** New Neoplasm post initial therapy. **(n)** Neoplasm status. **(o)** Lymph Node Stage. Dataset, TCGA. Patient numbers, on graph. Data presented as % of samples in each quartile grouping in presented categories. P-values, chi-squared test. **p-r.** Progression-free survival of prostate cancer patient groups based on quartiles (Q1 versus Q2-4 combined) of expression in tumour samples of **(p)** *ARF3* mRNA, **(q)** *CDH2* mRNA, or **(r)** CDH2/N-cadherin protein. P-values, log rank test. Dataset, TCGA. Patient numbers, on graph. **s.** Progression-free survival of prostate cancer patient groups based on median split (M1, low versus M2, high) of CDH2/N-cadherin protein and *ARF3* mRNA expression in prostate tumours. P-values, log rank test. Dataset, TCGA. Patient numbers, on graph. **t.** Neoplasm status upon grouping of prostate cancer patients based on median split (M1, low versus M2, high) of CDH2/N-cadherin protein and *ARF3* mRNA expression in prostate tumours. Dataset, TCGA. Patient numbers, on graph. Data presented as % of samples in each quartile grouping in presented categories. P-values, chi-squared test.

Given the requirement of N-cadherin for ARF3 function *in vitro*, we examined whether the effect of ARF3 requires consideration of N-cadherin. Only four tumour types showed a consistent alteration in mRNA levels of N-cadherin (gene: *CDH2*) across the independent datasets: increased *CDH2* levels in thyroid (THCA) and kidney (KIRP), decreased *CDH2* mRNA levels in colon (COAD) and prostate (PRAD) adenocarcinoma (Fig. 7h; S6c,d). Decreased *CDH2* mRNA levels were also observed when comparing normal and tumour prostate tissue samples matched from the same patients (Fig. 7i). As N-cadherin (but not ARF3) protein expression was also surveyed in Reverse Phase Protein Array (RPPA) analysis of the TCGA cohort, this allowed examination of both mRNA and protein of CDH2 in prostate cancer patients. Notably, *CDH2* mRNA only modestly correlated with N-cadherin protein (Fig. 7j). We therefore examined whether *CDH2* mRNA or N-cadherin protein levels could be used to identify groups of prostate cancer patients with different clinical characteristics. N-cadherin protein levels showed a modest decrease with progressive Gleason Grade, while mRNA did not show significant differences (Fig. 7k,l). When divided into quartiles of expression, ascending N-cadherin protein expression, but not *CDH2* mRNA levels, showed a significant, inverse decrease in the frequency of patients presenting a new neoplasm following initial therapy (protein, *p*=0.0074; mRNA, not significant; Fig. 7m), whether patients were with or without tumour (protein, *p*=0.0228; mRNA, not significant; Fig. 7n), and lymph node metastasis positivity (protein, *p*=0.0213; mRNA, not significant; Fig. 7o). This indicates that N-cadherin protein is partially uncoupled from *CDH2* mRNA levels, and that low N-cadherin levels identifies patients with recurrent, metastatic tumours.

Analysis of progression-free survival of prostate cancer patients (TCGA) indicated that neither *ARF3* nor *CDH2* mRNA levels could stratify patient groups with altered survival (Fig. 7p,q). In contrast, patients with lowest N-cadherin protein levels showed drastically reduced progression-free survival (compare lowest quartile (Q1) to all other patients, Q2-4; p=0.0007; Fig. 7r). We examined whether combining N-cadherin protein expression with *ARF3* mRNA levels would further stratify patient survival by comparing patient groups segregated by expression based on a median split (M1, low; M2, high; Fig. 7s). Patients with low N-cadherin protein (M1) showed similar survival regardless of *ARF3* mRNA levels (yellow and blue lines), whereas in high N-cadherin expressing patients (M2) the mRNA levels of *ARF3* divided survival patterns; having low *ARF3* (M1) mRNA despite high N-cadherin expression (M2) (green line) reduced survival to levels mirroring low N-cadherin protein, while conversely having both high N-cadherin protein (M2) and *ARF3* mRNA (M2) (red line) identified patients with best progression-free survival. Similarly, high levels of CDH2 protein and ARF3 mRNA identified a patient group with lowest levels of new tumour formation after initial therapy, while all other groups showed similar rates (Fig. 7t). This clinical data is consistent with our *in vitro* data identifying a co-operation between N-cadherin and ARF3, wherein ARF3 and N-cadherin mutually control each other’s levels and function in tumourigenesis, with reduced N-cadherin protein associated with metastatic, recurrent prostate cancer. These data also indicate ARF3 as a contextual regulator of N-cadherin protein levels during tumourigenesis in prostate cancer patients.

## Discussion

The use of 3D culture allows the assessment of how individual genes or entire pathways contribute to collective cell behaviours. The application of a screening approach to 3D requires a number of adaptions not directly transferrable from the screening of 2D cell cultures. First, collective morphogenesis occurs over multiple days, requiring stable genetic manipulations to which siRNA transfections are poorly suited. Moreover, 3D phenotypes can be stereotyped, but asynchronous, or multiple phenotypes can occur in parallel in the same well. This requires live-imaging of 3D morphogenesis to capture and quantify these considerations.

A bottleneck in the systematic screening of collective morphogenesis (arrayed, one manipulation per well) is the plating of multiple parallel manipulations into 3D culture at the same starting density. This is essential to ensure that phenotypes quantified do not simply represent morphogenesis from a different starting point, such as altered density. If the genetic manipulation of interest alters proliferation before plating into 3D culture, this is a technical challenge in assuring similar plating. We have overcome these obstacles using an arrayed shRNA library vector that co-encodes membrane-targeted mVenus fluorescent protein (mem:Venus) to allow plating at similar density in 3D culture. Through the use of phase contrast and fluorescent imaging of 96-well plates of 3D culture, tracking of morphogenesis of mem:Venus-positive acini, and machine learning-based classifications of distinct phenotypes, we were able to perform a functional genomic characterisation of the ARFome contribution to collective morphogenesis.

It is important to note that despite using a library with high independent validation of target depletion from shRNAs, our approach is not an exhaustive analysis of every ARFome member. Rather, shRNAs are assigned into Phenotype Groups based on their relative change in Round, Spindle or Spread acinus phenotype over time compared to a control (Scr) shRNA (Fig. S2a). While this can detect phenotypes such as being highly Round (Group 3, e.g. *IL6ST*) or highly Spindle (Group 2, e.g. *RAB11FIP3*), phenotypes with modest change to control (Group 7, e.g. *ARF4*) were also identified. Modest changes can occur due to a *bona fide* lack of strong phenotype or could be due to inconsistent effect across the three independent instances we performed the screen. Therefore, lack of robust effect should be interpreted through the lens of such limitations of large-scale screens, rather than definitive demonstration of a lack of function of such ARFome members.

Of those ARFome members that exhibited notable phenotypes, our screen identified the Class I ARFs, the GEF PSD, and the effector RAB11FIP4 as repressors of Spindle- and Spread-type collective invasion. Particularly, we identify that loss of ARF3, not ARF1, in 3D culture phenocopies PSD and RAB11FIP4 loss of function. That depletion of Class I ARFs induced invasive activity was somewhat unexpected, as numerous studies, particularly in breast cancer cells, report a pro-invasive and pro-tumourigenic function of ARF1^11–19^.

In our studies, we define that ARF3 has a function distinct from that of ARF1. Notably, these 3D phenotypes, which by definition require multicellular collective function, were not recapitulated when looking at the shape of single cells in 2D, which may explain some of the differences to observations using 2D culture. Moreover, ARF3 expression was strongly induced in 3D culture, suggesting a requirement for collective function. That we identify such co-acting modules using a functional morphogenesis perturbation approach is notable as this would be difficult to predict from studies of single cells in the literature. Of ARFome expression in PC3 cells, PSD was the lowest expressed GEF, while RAB11FIP4 sits at approximately the mid-point of the ARFome interactors screened, which makes selection of these candidates non-obvious. PSD is also known as Exchange Factor for ARF6 (EFA6)- A, which represents its prior consideration as a GEF for mostly ARF6. However, most ARF GEFs can have broad action, and PSD can efficiently catalyse GTP-loading of ARF1 but was reported to only modestly GTP-load ARF3 ^8, 20^. However, PSD GEFs can also act as effectors of GTP-loaded ARF proteins ^8^. Whether PSD acts as an ARF3 GEF or ARF3 effector remains to be demonstrated.

We identified that RAB11FIP4 is likely a key ARF3 effector, and that N-cadherin is a key cargo protein of ARF3-RAB11FIP4 complex on endosomes. RAB11FIP4, also known as Arfophilin-2, is a dual Rab11- and ARF-binding protein that controls the organisation of, and trafficking through, the Rab11 recycling endosome^21–23^. In our screen, the depletion of the related protein RAB11FIP3/Arfophilin-1 also induces a Spindle-type 3D phenotype, suggesting that the RAB11 endosome might be a major target of ARF3 function. Indeed, ARF1 and ARF3 have been reported to control recycling endosome function independent to effects on the Golgi apparatus^24^. The related protein RAB11FIP1 regulates recycling of endocytosed N-cadherin to promote cell migration^25^. Similarly, knockdown of RAB11FIP3 in neuronal cells causes the intracellular retention of N-cadherin, which can only be rescued by expression of RAB11FIP3 with functional ARF-binding capacity^26^. Therefore, the endosomal traffic of N-cadherin appears to be a major target of ARF3-RAB11FIP3/4 in our system. It is unexpected to identify RAB11FIP3/4 proteins as phenocopying ARF3 function, as previous reports suggest Arfophilins to be Class II or Class III ARF effectors, with only modest binding of Class I ARFs ^27, 28^. Given the widespread overlap in binding capabilities within the ARFome, this emphasises the need for phenotypic screening to identify co-acting modules within the ARFome.

In our system, ARF3 interacted with N-cadherin and acted as a rheostat to control N-cadherin levels in cells without affecting E-cadherin levels. Reciprocally, decreased N-cadherin levels also reduced ARF3 levels, suggesting a mutual regulation of the complex. Notably, in cardiomyocytes ARF3 interacts with N-cadherin but not E-cadherin, while conversely ARF4 interacts with E-cadherin but not N-cadherin ^9^, identifying specificity of different ARFs for different cadherins. Given that RAB11FIP4 regulates N-cadherin levels, and that ARF3 and N-cadherin co-localise in endosomal compartments, regulation of recycling of N-cadherin by ARF3-RAB11FIP4 may be a mechanism by which the latter complex acts as a rheostat to control levels of the former. How N-cadherin in turn controls ARF3 levels is unclear.

The functional consequence of altering ARF3 was to control the modality of invasion in cells through influencing junctional F-actin levels and N-cadherin-dependent collective sheet-type movement. This activity unique to ARF3, compared to ARF1, required three residues in the ARF3 C-terminus (A174/K178/K180). How these residues uniquely couple ARF3 to N-cadherin function is the focus of future work. Although conceptually it is understandable that increased levels of a cadherin could increase junctional stability and induce sheet-type movement, that this was conferred by N-cadherin was unexpected. Conversely to our findings, N-cadherin overexpression has been reported as a targetable and prominent poor prognostic indicator of prostate cancer outcome and metastasis^29^. Some of this discrepancy may be due to experimental system; some of these studies involve ectopic overexpression of N-cadherin in E-cadherin-negative cells. Notably, in our studies both N-cadherin and E-cadherin are expressed in PC3 cells and alteration of N-cadherin levels occurred without changing E-cadherin levels, indicating that this was not a transcriptional change between cadherin types (i.e. a ‘cadherin switch’). Effects on N-cadherin on collective behaviours are likely to differ greatly if other Type-I classical cadherins are not present. Despite this, examination of prostate cancer patients indicated that low protein levels of N-cadherin in prostate tumours were associated with metastatic tumours with poor progression-free survival. Notably, when N-cadherin levels were high in patients, combining this with ARF3 levels further stratified patients, wherein ARF3^HI^-N-cad^HI^ patients presented with the best clinical outcome. Moreover, the ability of N-cadherin to stratify survival was specific to protein, not mRNA, levels, indicating that the trafficking of N-cadherin, such as by the ARF3-RAB11FIP4 rheostat identified here, is essential for N-cadherin influence on tumorigenesis.

Our studies indicate that ARF3-RAB11FIP3/4-N-cadherin function to control the modality of invasion *in vitro*. By using a murine model of intraprostatic xenograft of ARF3-manipulated cells, we could examine which modality was required for metastasis *in vivo*. While both ARF3 depletion and overexpression conditions still grew tumours and metastasised, the type and frequency of metastasis was dramatically altered. ARF3 depletion resulted in completely penetrant, widespread metastasis, while ARF3-overexpressing cells manifested low efficiency, local metastases. This was mirrored in analysis of prostate cancer patients who possessed metastatic disease when the key ARF3-RAB11FIP4 cargo N-cadherin protein levels were low. In breast cancers, ARF3 upregulation was reported as associated with pro-proliferative functions^30^, while in gastric cancer, similar to our findings in prostate cells, ARF3 suppressed proliferative function and *in vivo* tumourigenesis^31^. Indeed, in our own analysis while ARF3 expression was commonly altered in tumour compared to normal tissue, the directionality of expression change was dependent on tumour type. Given that control of N-cadherin levels was a major function of ARF3, this seeming incongruent effect of ARF3 may be due to a differential effect of N-cadherin in alternate tissue types. Therefore, rather than being a generalised good or bad indicator of tumourigenesis, the ARF3-N-cadherin rheostat may be a contextual regulator of tumourigenesis in different tumour types. Collectively, this indicates that our screening approach to identify co-acting ARFome modules using 3D live-imaging approaches identifies an unexpected function for the Class I ARF, ARF3, in regulation of N-cadherin influence on collective invasion dynamics during metastasis.

## Methods

### Cell Culture

HEK293-FT cells (Thermo Fisher Scientific) were cultured in Dulbecco’s Modified Eagle Medium (DMEM) with 10% Fetal Bovine Serum (FBS), 2mM L-glutamine and 0.1mM Non-Essential Amino Acids (NEAA) (all Gibco). PC3 cells (ATCC) were maintained in Ham’s F-12K (Kaighn’s) Medium (Gibco) supplemented with 10% FBS. RWPE-1, RWPE-2, WPE-NB14 and CA-HPV-10 cell lines (ATCC) were grown in Keratinocyte Serum Free Medium (K-SFM) supplemented with 50 μg/ml Bovine Pituitary Extract (BPE) and 5ng/ml Epidermal Growth Factor (EGF) (all Gibco). LNCaP and VCaP cells (ATCC) were cultured in RPMI-1640 or DMEM respectively (Gibco) containing 10% FBS and 2mM L-glutamine. DU145 cells (ATCC) were maintained in Minimum Essential Medium (MEM) (Gibco) with 10% FBS and 2mM L-glutamine. 22Rv1 cells (ATCC) were grown in phenol free RPMI-1640 containing 10% charcoal stripped FBS and 2mM L-glutamine (all Gibco). Cells were routinely checked for mycoplasma contamination and HEK293-FT, PC3, RWPE-1, RWPE-2, LNCaP and DU145 cells were authenticated using short tandem repeat (STR) profiling.

### ARFome shRNA screen

#### Generation of shRNA

Oligonucleotides for shRNAs were synthesised for 210 ARFs, GEFs, GAPs and effectors (Thermo Fisher Scientific) based on short hairpin sequences available from the RNAi Consortium (Broad Institute). Sequences and validation data are shown in Supplementary Table S1. shRNA oligonucleotides were cloned into a modified lentiviral vector, pLKO.4-Membrane-Venus (mem:Venus). This was constructed from pLKO.1-puro and the puromycin-resistance cassette replaced with an mVenus fluorescent protein with an N-terminal membrane-targeting domain of GAP43 (derived from approach used in ^32^), and was also modified to contain a single XhoI site. This allowed for screening of successful ligation of shRNA cassettes into the vector, which introduced a second XhoI site in the form of the shRNA loop (5’-CTCGAG-3’). The library generated was therefore improved by allowing for XhoI restriction digest screening of shRNA-containing sequences prior to plasmid sequencing (University of York). The resultant plasmid DNA was mini-prepped and plated into 96 well shRNA source plates at 100ng/μl for use in transient transfections.

#### Lentiviral infection of PC3

HEK293-FT cells were washed in PBS then dissociated by addition of 0.25% Trypsin in PBS/EDTA. Detached cells were collected in medium, cell number determined and 13,000 cells added per well of 96 well plates. Plates were then incubated at 37°C, 5% CO_2_ overnight. Lentiviral packaging vectors psPAX2 (Addgene plasmid 12260) (100ng/well) and VSVG (pMD2.G; Addgene plasmid 12259) (10ng/well) were diluted in OPTI-MEM (10μl/well, Thermo Fisher Scientific) and dispensed into 96 well plates. Lipofectamine 2000 (Thermo Fisher Scientific) was also diluted in OPTI-MEM (0.6μl and 10μl per well) and dispensed into 96 well plates. After 5 minutes at room temperature the contents of these plates were combined and 1.25μl of each shRNA added (final concentration of 125ng/well). Plates were mixed briefly using a shaker, incubated at room temperature for 5 minutes and then added to HEK293-FT cells overnight at 37°C, 5% CO_2._ Supernatant was removed and replaced with HEK293- FT medium containing an additional 10% FBS and plates incubated for a further 24 hours at 37°C, 5% CO_2._ Viral supernatant was then removed and replaced again after 24 hours. The viral supernatant collected after 24 and 48 hours was combined in 96 well plates and centrifuged at 300g for 4 minutes. 150μl of supernatant was decanted from each well and added to 96 well plates containing 50μl Lenti-X Concentrator (Clontech) at 4°C for 1 hour. Plates were then centrifuged at 1,100g for 45 minutes at 4°C and the resultant pellets re-suspended in PC3 medium and added to PC3 cells plated in 96 well plates (at 6,000 cells/well) 24 hours prior to infection.

Additional PC3 medium (50μl/well) was added after 24 hours and the plates incubated for a further 48 hours at 37°C, 5% CO_2._ PC3 cells were then washed in PBS and 50μl of 0.25% Trypsin in PBS/EDTA added slowly to each well. This was removed immediately and replaced with 5μl of Trypsin per well for 7 minutes with gentle shaking. A further 5μl of Trypsin was added to the centre of each well for 1 minute, without shaking, to help break up any cell clumps. PC3 medium was added, and the cells split into 4 96 well TC treated plates (Greiner, 655090) which were incubated at 37°C, 5% CO_2_ for use in assays.

#### Live imaging of PC3 in 3D

A visual inspection was carried out after 24 hours to identify plates with consistent confluence across wells. Cells were washed in PBS, trypsinised as described above and re-suspended in PC3 medium. Typically, 1/2 to 2/3 of cells from each well were used for subsequent 3D cultures depending upon the initial confluence and effectiveness of trypsinisation. Medium was supplemented with 2% Growth Factor Reduced Matrigel (GFRM; BD Biosciences) and added to 96 well ImageLock plates (Satorius) pre-coated with 10μl of GFRM for 15 minutes at 37°C. For live imaging ImageLock Plates were incubated at 37°C for 4 hours, then imaged using an IncuCyte ZOOM (Satorius) with IncuCyte ZOOM Live Cell Analysis System Software 2018A. Phase and GFP images were taken every hour for 4 days at 2 positions per well with x10 objective lens.

#### Analysis

CellProfiler (Version 3.1.8) was used to design a pipeline to process the phase images acquired on the IncuCyte, including retention of only mem:Venus-positive objects. This pipeline identified and tracked acini at each timepoint and generated a database containing size, shape and movement measurements. CellProfiler Analyst (Version 2.2.0) was then used to apply iterative, user-supervised machine learning to spheroid measurements in the resulting database. Specifically, PC3 acini were classified into bins based on their morphology: Round, Spindle or Spread. Machine learning to differentiate between these classes, using a maximum of 20 rules with Fast Gentle Boosting, was used until accuracy for each class was >90%, as assessed using a confusion matrix. Once generated, these classification rules were saved as a .txt file and imported into CellProfiler for classification without need for re-training of new data sets.

Custom pipelines in KNIME Data Analytics Platform (Version 3.3.1) were then used to collate data from experimental and technical replicates, filter datasets and apply tracking label corrections. For user-defined phenotypes, heatmaps show phenotype frequency over time in comparison to control samples. The relative proportion of the total acini in each sample is presented as Z-score normalised data in 12-hour time intervals using ggpot2 R package. P values correspond to circle size. Statistical comparison was performed using the Cochran-Mantel-Haenszel test, which takes into account experimental replicates and is only statistically significant when the effect was present across all replicates. We also used the Breslow-Day statistic to test whether the magnitude of effect was homogeneous across all experimental replicates – a non-significant p-value indicated homogeneity and is represented by a black dot in the heatmap. For this, we used the DescTools R package. In both statistical tests, a Bonferroni adjustment was applied to account for multiple testing. For detection of phenotype classes, shRNAs were grouped into 7 groups based on a dendrogram from clustering the fold change over time for each shRNA in Round, Spindle, and Spread phenotypes compared to the control (Scr) value. The dendrogram was generated using hierarchical clustering of heatmap data by complete linkage of Euclidian distances between samples. Line plots are presented which show the mean ± s.e.m. of Round, Spindle and Spread phenotypes over time for each group and for Scr shRNA controls. The average proportion of acini classified as Round, Spindle or Spread across all time points is also shown for each group.

The entire shRNA screen, all the way from virus production to phenotype quantitation, was performed three independent times. Each replicate consisted of 18 technical replicates (3 per 96 well plate) of control (Scr) shRNA and 1 replicate of each of the 210 ARFome shRNAs. 170,674 Scr mem:Venus shRNA-expressing acini were quantified in total. Supplementary Table S1 shows the total number of acini quantified for each mem:Venus shRNA.

### Generation of stable cell lines

Short hairpin sequences in pLKO.1-puromycin lentiviral vector (Sigma Aldrich) were used to generate stable knockdown. The specific shRNA sequences used are shown in Supplementary Table S2. GFP-tagged ARFs were kind gifts from P. Melançon (University of Alberta) and alternate fluorescent tags, mutations and chimeras were generated by sub-cloning. All RNAi-resistant variants and chimeras were made by mutagenesis or sub-cloning using fragment synthesis (GeneArt). Plasmids will be deposited with Addgene upon publication.

Stable cell lines were made by co-transfecting lentiviral packaging vectors VSVG (pMD2.G; Addgene plasmid 12259) and psPAX2 (Addgene plasmid 12260) with plasmid of interest, into HEK293-FT using Lipofectamine 2000 (Thermo Fisher Scientific) as per manufacturer’s instructions. Viral supernatants were filtered using PES 0.45μm syringe filters (Starlab) and concentrated using Lenti-X Concentrator (Clontech) as per the manufacturer’s instructions. PC3 cells were transduced with lentivirus for at least 3 days prior to FACS sorting or selection with 300μg/ml G418, 2.5μg/ml puromycin (both Thermo Fisher Scientific) or 10μg/ml blasticidin (InvivoGen). To allow direct comparison PC3 cells expressing mNG were used for shRNA expression and Scr shRNA was added to cells over-expressing mNG-tagged plasmids.

### Live 3D culture and analysis

Stable cells lines were used to form acini with 2,250 cells plated per well in a 96-well ImageLock plate pre-coated with 10 μl of GFRM for 15 minutes at 37°C. Medium was supplemented with 2% GFRM. Plates were incubated at 37°C for 4 hours, then imaged using an IncuCyte ZOOM or IncuCyte S3 (Satorius). Images were taken every hour for 4 days at 2 positions per well using a x10 objective lens.

The CellProfiler pipeline and classification rules generated in CellProfiler Analyst previously described for the ARFome shRNA screen were also used to analyse these experiments. A Custom pipeline in KNIME Data Analytics Platform (Version 3.3.1) was used to collate data from experimental and technical replicates, filter datasets, apply tracking label corrections and overlay coloured outlines onto phase images. This pipeline then generated a heatmap of mean features over time (i.e. area) with statistical comparison to control sample. Data are presented in heatmaps as Z-score normalised in 12-hour time intervals using ggpot2 R package. P values correspond to circle size. Statistical comparison was performed by Student’s t-test, two-tailed, and a Bonferroni adjustment was applied to account for multiple testing. For user-defined phenotypes, heatmaps show phenotype frequency over time in comparison to control samples as described for ARFome shRNA screen. Number of experimental replicates (n), the number of technical replicates per experiment and the number of acini quantified in total per condition are stated in Figure Legends.

### Fixed 3D culture and analysis

PC3 acini were cultured in GFRM as described in ‘Live 3D culture and analysis’ section and incubated for 3 days at 37°C, 5% CO_2._ Acini were gently washed with PBS prior to addition of 4% paraformaldehyde for 15 minutes. Samples were blocked in PFS (0.7% fish skin gelatin/0.025% saponin/PBS) for 1 hour then stained with primary antibodies overnight at 4°C with gentle agitation. After 3 x 5 minute washes in PFS, secondary antibodies (all Thermo Fisher Scientific) were added for 45 with gentle agitation at room temperature. Acini were washed 3x in PBS for 5 minutes each and maintained in PBS at 4°C until imaging was carried out. Antibodies used described in Supplementary Table S3.

Confocal images were taken using A1R microscope (Nikon) with x40 objective, exported as TIFF files and processed in Fiji. Other images were taken using an Opera Phenix™ High Content Screening System (Perkin Elmer). 35 images and 15 planes were taken per well with a 20x objective for PC3 acini. Columbus Image Data Storage and Analysis System (PerkinElmer, version 2.9.1) was used to design a custom pipeline to measure F-actin intensity per acini. F-actin staining was used to detect individual acini in maximum projection images of all planes and any acini touching the image border were excluded from further analysis. Data is presented in box and whiskers plot as total F-actin intensity per acinus. The percentage of acini with visibly reduced F-actin intensity at junctions in maximum projection images is also shown. Values are mean ± s.e.m. and P values and statistical test used are described in Figure Legends.

Number of experimental replicates (n), number of technical replicates per experiment and the number of acini quantified in total per condition are stated in Figure Legends.

### 2D Morphology Assays

Method described previously^3^, briefly cells were plated on 96 well plates for 48 hours, fixed and then stained, as described in ‘Fixed 3D culture and analysis section’, using Hoechst 34580, Alexa Fluor 568 Phalloidin and HCS CellMask™ Deep Red Stain. Plates were imaged using an Opera Phenix™ High Content Screening System (x10 or x63 objective) and Columbus Image Data Storage and Analysis System (PerkinElmer, version 2.9.1) used to design a custom pipeline for analysis.

Cells were detected based on nucleus localization (Hoechst) and the shape of each cell defined by either F-actin or HCS CellMask staining. Any cells touching the image border were excluded from further analysis. Where appropriate, cells expressing mNG-tagged proteins were detected using fluorescence intensity properties and any cells not expressing protein of interest were filtered from further analysis. Morphology properties of each object were calculated to classify them into three different categories (Round, Spindle and Spread) using machine learning following manual training. A custom pipeline was generated using KNIME Data Analytics Platform (Version 3.3.1) to collate data from independent and technical replicates, calculate the log2 fold change of each phenotype over control and to calculate statistical significance using one-way ANOVA. Number of experimental replicates (n), number of technical replicates per experiment and total number of cells quantified per condition are stated in each Figure Legend.

### 2D Proliferation Assays

1,000 PC3 cells were plated per well in a 96-well ImageLock plate then imaged using an IncuCyte ZOOM (10x objective, 2 images per well) each hour for 96 hours. Confluence per well was then measured using the IncuCyte ZOOM analysis software. Mean confluence for each condition at each time point (from 4 technical replicates) was calculated and normalised to time 0 (t = 0) for 3 experimental replicates. Mean data ± s.e.m. is presented in a line graph (PRISM 7, GraphPad) as confluence normalised to t = 0 at 12 hour time points. P values and the statistical test used are described in Figure Legends.

### 3D Proliferation Assays

PC3 acini were set up, as described in ‘Live 3D culture and analysis’ section, on four 96-well TC-treated plates with 2,250 cells plated per well. Plates were maintained at 37°C, 5% CO_2_ for 4, 24, 48 and 72 hours. At these time points 100 μl CellTiter-Glow (Promega) was added to each well and plates placed on an orbital shaker at low speed for 5 minutes. Plates were then removed and incubated at room temperature for a further 20 minutes. Luminescence was measured using a Tecan SPARK Microplate Reader (Tecan). Mean luminescence for each condition at each time point (from 3 technical replicates) was calculated and normalised to time 0 (4 hours) for 4 experimental replicates. Mean data ± s.e.m. is presented in a line graph (PRISM 7, GraphPad) as luminescence fold change to t = 0 at 24 hour time points. P values and the statistical test used are described in Figure Legends.

### Invasion and Migration Assays

Method described previously^3^, briefly ImageLock plates were coated with 20μl of 10% GFRM diluted in medium overnight at 37°C. 70,000 PC3 cells in 100μl medium were plated in each well for 4 hours at 37°C. The resultant monolayer was wounded using a wound making tool (Satorius), washed three times with medium and overlaid with either 50μl of 25% GFRM for invasion assays or 100μl medium for migration assays. After incubation at 37°C for an hour 100μl medium was added to each well of invasion assay and plates imaged every hour for 4-6 days using the IncuCyte ZOOM.

For each experimental replicate the average Relative Wound Density (RWD) of all control samples (Scr shRNA) was calculated and each technical replicate normalized to this value. Results are presented as RWD at the time point at which the average RWD of the control samples is 50% (t=Max_1/2_) or 25% (t=Max_1/4_). Mean values ± s.e.m. (triangles) and technical replicates (circles) for each independent experiment are presented. The number of experimental replicate (n) and technical replicates per experiment is also stated in the appropriate Figure Legend. P values and the statistical test used are described in Figure Legends.

### Immunoblotting

2x10^5^ cells were plated in 6 well plates for 48 hours. Plates were washed twice with ice cold PBS then RIPA lysis buffer added for 15 minutes on ice (50mM Tris-HCl, pH 7.4, 150mM NaCl, 0.5mM MgCl2, 0.2mM EGTA, and 1% Triton X-100 with cOmplete protease inhibitor cocktail and PhosSTOP tablets (Roche)). Cells were scraped and lysates clarified by centrifugation at 13,000 rpm for 15 minutes at 4°C. BCA Protein Assay kit (Pierce) was used to determine protein concentration as per manufacturer’s instructions.

RWPE-1 or PC3 acini were cultured in 3D as described above by plating 2x10^5^ cells supplemented with 2% GFRM into 6 well plates pre-coated with 180 μl of GFRM for 30 minutes at 37°C. After 2-3 days acini were lysed in RIPA as described for 2D samples then slowly passaged 10x through a 25-27G needle. After centrifugation, 13,000 rpm at 4°C for 15 minutes, the supernatant was separated from the lower layer of GFRM and debris and used for SDS PAGE.

SDS-PAGE was performed using Bolt 4-12% Bis-Tris gels and buffers (Thermo Fisher Scientific, as per the manufacturer’s instructions, typically 20μg per sample) and proteins transferred to PVDF membranes using the iBlot 2 transfer system (Thermo Fisher Scientific). Membranes were incubated for 1 hour in Rockland blocking buffer (Rockland) and primary antibodies added overnight at 4°C. Antibodies used described in Supplementary Table S3. After addition of appropriate secondary antibodies for 1 hour, membranes were washed in TBST three times and imaged using a ChemiDoc Imager (BioRad) or Odyssey Imaging System (LI-COR Biosciences). Bands were quantified using Image Lab 6.1 (BioRad) or Image Studio Software 6.0 (LI-COR Biosciences). The number of independent lysate preparations (n) is stated in the appropriate Figure Legend and quantitation of fold change of protein expression is shown as mean ± s.e.m. P values and the statistical test used are described in Figure Legends. GAPDH was used as a loading control for each blot and a representative blot for each sample set is shown where appropriate.

### Immunoprecipitation

1mg of cell lysate was immunoprecipitated with 2μg anti-ARF3 antibody (BD, 610784) overnight at 4°C with gentle rotation. Either anti-mouse agarose or mouse agarose (IgG control) (both Sigma) were added for 1 hour at 4^0^C with rotation. Samples were washed three times in RIPA lysis buffer, separated by SDS–PAGE, and immunoblotted. n=3 experiments from independent lysate preparations.

### ARF-GTP Pulldown

Method described previously^3^, briefly PC3 cells were incubated on 120mm plates for 48 hours at 37°C, 5% CO_2._Cells were then lysed on ice in pulldown-lysis buffer (50mM Tris, 100mM NaCl, 2mM MgCl2, 0.1% SDS, 0.5% Na-deoxycholate, 1% Triton X-100, 10% glycerol). Lysates were syringed 5x using a 25-27G needle and centrifuged at 4°C 14,000g for 1 minute. Spin columns were equilibrated with 50μl of Glutathione Agarose resin and washed with pulldown-column wash buffer (1:1 pulldown-lysis buffer and 1xTBS). 80μg of GST-GGA3-GAT recombinant fusion protein was immobilised on the agarose resin by incubation at 4°C with gentle rocking. After 1 hour 1 mg of each lysate was added onto spin columns and incubated again at 4°C for 2 hours with rocking. Pulldown wash buffer (50mM Tris, 100mM NaCl, 2mM MgCl2, 1% NP-40, 10% glycerol) was used to wash unbound proteins off the column. 60μl of pulldown-elution buffer (10mM Glutathione in 1xTBS) was added to each spin column and incubated for 5 minutes at room temperature. Eluted protein was collected at 1,250g for 1 minute and samples prepared for SDSPAGE and immunoblotting as described above. n = 3 experimental replicates. Data is presented as GGA3 binding normalised to ARF3 levels for each experiment. Values are mean ± s.e.m.

### RNA-sequencing

Briefly, RNA was extracted using an RNeasy kit, incorporating a DNase digestion step using RNase-Free DNase Set (both Qiagen), followed by reverse transcription using High-Capacity cDNA Reverse Transcription kit (Thermo Fisher Scientific) all per the manufacturer’s instructions. Quality control of all RNA samples was performed using a 2200 Tapestation and High-sensitivity RNA screentape (Agilent) and only samples with RIN values >7.9 were processed. Libraries were then prepared using 1ug of total RNA as per the manufacturer’s instructions (Illumina TruSeq stranded mRNA). Libraries were sequenced using an Illumina NextSeq 500, on a High-Output 75 cycle run with paired-end 36bp read length.

For RNA-sequencing analysis, quality checks and trimming on the raw RNA-seq data files were performed using FastQC version 0.11.8, FastP version 0.20 and FastQ Screen version 0.13.0. RNA-seq paired-end reads were aligned to the GRCm38 version of the human genome and annotated using HiSat2 version 2.1.0. Expression levels were determined and analysed using a combination of HTSeq version 0.9.1, the R environment version 3.6.1, utilizing packages from the Bioconductor data analysis suite and differential gene expression analysis based on the negative binomial distribution using the DESeq2 package version 1.22.2. Pathway Analysis was preformed using MetaCore from Clarivate Analytics (https://portal.genego.com/).

### Animal studies

Animal experiments were performed in compliance with all relevant ethical regulations and approvals of the relevant UK Home Office Project Licence (70/8645 and P5EE22AEE) and carried out with ethical approval from the Beatson Institute for Cancer Research and the University of Glasgow under the Animal (Scientific Procedures) Act 1986 and the EU directive 2010 and sanctioned by Local Ethical Review Process (University of Glasgow).

7-week-old CD1-nude male mice were obtained from Charles River (UK) and acclimatised for at least 7 days. Mice were kept in a barriered facility at 19-22 °C and 45-65% humidity in 12 h light/darkness cycles with access to food and water ad lib and environmental enrichment. 2 x 10^6^ PC3 cells stably expressing mNG and either Scr shRNA (20 mice) or *ARF3* KD1 shRNA (18 mice) or expressing ARF3-mNG and Scr shRNA (17 mice) were surgically implanted into one of the anterior prostate lobes of each mouse (under anaesthesia and with analgesia). The mice were continually assessed for signs of tumour development (including by palpation) and humanely sacrificed at an 8-week timepoint, prior to tumour burden becoming restrictive. Primary tumour (PT) and macrometastasis (MM) incidence were analysed by gross observation and are presented as number of mice with PT or MM incidence only in mice with a PT. Macrometastasis count per mouse and weight of prostate is also shown in box and whiskers plots only for mice with PTs (Scr shRNA, 12/20 mice, *ARF3* KD1 shRNA, 12/18 mice and ARF3-mNG and Scr shRNA 9/17 mice). P values and the statistical test used are described in Figure Legends.

### Analysis of patient cohorts

Patient data i.e. copy number and RNAseq data from Cancer Cell Line Encyclopedia (CCLE) was accessed, analysed and downloaded using in-platform cBioportal.org tools ^33, 34^. Copy number and mRNA levels (RNAseq) for prostate normal and cancer cell lines were downloaded from the Cancer Cell Line Encyclopedia (CCLE; https://sites.broadinstitute.org/ccle/; ^35^). Patient datasets for normal and tumour samples from The Cancer Genome Atlas (TCGA, pan-cancer) were downloaded from the TCGA Splicing Variants database (TSVdb.com, ^36^). Data for the combination of normal and tumour were downloaded from the Gene Expression for Normal and Tumour database (GENT2, http://gent2.appex.kr/gent2/, ^10^), with sample numbers and study IDs in Supplementary Tables S4 and S5. Datasets for Normal, Tumour and Metastasis (Glinsky ^37^), Grasso (GSE35988), Lapointe (GSE3933), Taylor (GSE211032), TCGA (obtained from cBioportal.org), Tomlins (GSE6099) and Varambally (GSE3325)) were downloaded from CANCERTOOL ^38^ and analysed using GraphPad Prism 9.

## Data Availability

RNA-seq data will be deposited in the Short Read Archive database. Any other data that supports the findings of this study will be available from corresponding author.

## Acknowledgements

This work was supported by the following grants: D.M.B. NIH K99CA163535, CRUK (C596/A19481) E.S., AR.-F, and L.M. CRUK C596A17196 and A31287. E.C.F. was supported by a University of Glasgow Industrial Partnership PhD scheme co-funded by Essen Bioscience, Sartorius Group. R.P. and H.L.Y funded by CRUK (A22904). S.M., J.A. L.G. and K.B. CRUK (A29799, A17196 and A31287).

We would like to thank the Core Services and Advanced Technologies at the Cancer Research UK Beatson Institute, with particular thanks to the Beatson Advanced Imaging Resource, Molecular Technologies and Biological Services Unit.

## Author Contributions

D.B conceived the project. Under supervision of D.B. lab work was performed by E.S., A.R.F., J.C. and L.M.G. E.C.F. and D.B. performed computational analyses. L.G., S.M., R.P. and J.A. performed *in vivo* murine experiments under supervision of K.B. and H.L.Y. Manuscript was written by D.B. and E.S. All authors read and commented on manuscript.

## Figure Legends

**Supplementary Figure 1: Expression levels of ARF GTPases vary in different prostate cancer cell lines in both 2D and 3D.**

**a-e.** Graphs generated using RNAseq data from the Cancer Cell Line Encyclopedia (CCLE) comparing **(a)** *ARF1,* **(b)** *ARF3,* **(c)** *ARF4,* **(d)** *ARF5* and **(e)** *ARF6* gene copy number and mRNA expression levels in multiple prostate cancer and non-transformed cell lines. Metastatic PC3 and normal prostate PLECLH cell lines, red and green respectively.

**f.** Western blot of androgen receptor (AR) proficient or deficient prostate cell lines for ARF1, ARF3, ARF4, ARF5 and ARF6. GAPDH is loading control for ARF6 and a sample control for all other blots. Panels shown are representative of 3 independent lysate preparations.

**g.** Western blot of RWPE-1 and PC3 acini, formed in GFRM (3D) for 2 days, for ARF1, ARF3, ARF4, ARF5 and ARF6. GAPDH is loading control for ARF3 and a sample control for all other blots. Panels shown are representative of 3 independent lysate preparations.

**h.** Graph is fold change of ARF expression in PC3 cells, normalized to RWPE-1 cells. Data presented as mean ± s.e.m. Panels shown are representative of 3 independent lysate preparations. P-values (Student’s 2-tailed t-test), n.s. not significant, *p 0.05 and **p ≤ 0.01.

**i-j.** Western blot of PC3 cells in 2D or in 3D for **(i)** ARF1 or **(j)** ARF3. GAPDH is loading control for each ARF blot. Graph, fold change of ARF expression, normalised to 2D samples. Dashed lines indicate blot was spliced. Data presented as mean ± s.e.m. and panels shown are representative of 3 independent lysate preparations. P-values (Student’s 2-tailed t-test), **p ≤ 0.01, ***p ≤ 0.001.

**k.** RNAseq data from PC3 cells shows mRNA expression (Log_2_) of genes encoding ARF GTPases, GEFs, GAPs, components of the IL6 signalling pathway and ARF effectors and interactors (Log_2_). n=4 mRNA samples prepared independently.

**Supplementary Figure 2: Effect of individual ARFome shRNA expression on collective cancer cell behaviour.**

**a.** PC3 cells expressing ARFome shRNA were plated on ECM with 2% ECM overlay and multi-day high-throughput imaging carried out live in 3D. Mem:Venus-positive acini classified into Round, Spindle or Spread phenotypes at each timepoint. Heatmap presents this classification, in 12- hour time intervals, as a Log_2_ Fold Change from control (Scr) (blue to red). The proportion of control at each timepoint is Z-score normalised for each class (white to black). P-values, Cochran-Mantel-Haenszel (CMH), Bonferroni adjusted, represented by the size of the bubble. Dot indicates p-value (Breslow-Day test, Bonferroni-adjusted) for consistent effect magnitude. shRNAs grouped into 7 groups (Phenotype Group 1 - 7) based on dendrogram generated using hierarchical clustering by complete linkage of Euclidian distances between samples. Viral infections and live 3D assays carried out 3 independent times. Each experimental replicate consisted of 18 technical replicates of Scr shRNA (170,674 acini in total) and 1 replicate of each of the 210 ARFome shRNAs (Supplementary Table S1).

**b-c.** Western blot of PC3 cells expressing mNG and Scr, **(b)** *ARF1* or **(c)** *ARF3* shRNA for ARF1, ARF3 and GAPDH as a loading control for each. Panels shown are representative of 3 independent lysate preparations. Graph is fold change of ARF expression normalized to Scr. Data is mean ± s.e.m. P-values (Student’s 2-tailed t-test), **p ≤ 0.01, ***p ≤ 0.001 and ****p ≤ 0.0001.

**d-e.** Western blot of PC3 cells expressing mNG and Scr, **(d)** *PSD* or **(e)** *RAB11FIP4* shRNA for PSD, RAB11FIP and GAPDH, as a loading control for each. Panels shown are representative of 2 independent lysate preparations.

**f.** Phase images of PC3 acini expressing mNG and Scr or *PSD* shRNA. Outlines; Round (red), Spindle (green) and Spread (blue). Scale bar, 100μm n=2 experimental replicates each with 4 technical replicates/condition. 10,723 (Scr), 13,920 (*PSD_*KD1), 15,157 (*PSD*_KD2) acini quantified in total.

**g.** Quantitation of **(f)**. Heatmaps, Area is mean of Z-score normalised values (purple to yellow). P-values, Student’s t-test, Bonferroni adjustment, represented by size of bubble. Heatmaps, Round, Spindle or Spread is Log_2_ Fold Change from control (Scr) (blue to red). Proportion of control at each time is Z-score normalised (white to black). P-values, Cochran-Mantel-Haenszel (CMH) test, Bonferroni adjusted, represented by size of bubble. Dot indicates p-value (Breslow-Day test, Bonferroni-adjusted) for consistent effect magnitude.

**h**. Phase images of PC3 acini expressing mNG and Scr or *RAB11FIP4* shRNA. Outlines; Round (red), Spindle (green) and Spread (blue). Scale bar, 100μm n=3 experimental replicates each with 3-4 technical replicates/condition. 14,551 (Scr), 16,435 (*RAB11FIP4_*KD1), 11,880 (*RAB11FIP4_*KD2) acini quantified in total.

**i.** Quantitation of **(h)**. Heatmaps, Area is mean of Z-score normalised values (purple to yellow). P-values, Student’s t-test, Bonferroni adjustment, represented by size of bubble. Heatmaps, Round, Spindle or Spread is Log_2_ Fold Change from control (Scr) (blue to red). Proportion of control at each time is Z-score normalised (white to black). P-values, Cochran-Mantel-Haenszel (CMH) test, Bonferroni adjusted, represented by size of bubble. Dot indicates p-value (Breslow-Day test, Bonferroni-adjusted) for consistent effect magnitude.

**Supplementary Figure 3: Depletion of Class I ARF GTPases affects 2D phenotype but not proliferation of prostate cancer cells.**

**a-b.** PC3 cells expressing mNG and Scr, **(a)** *ARF*1 or **(b)** *ARF*3 shRNA were plated at low density and imaged. Data is mean confluence ± s.e.m., normalized to time 0. n=3 experimental replicates with 4 technical replicates/condition. P-values (one-way ANOVA), n.s. not significant.

**c-d.** PC3 acini expressing mNG and Scr, **(c)** *ARF*1 or **(d)** *ARF*3 shRNA were plated for 4 to 72 hours. CellTiter-Glow was added and luminescence measured to assess ATP-based cell viability. Data is mean luminescence ± s.e.m., normalized to 4 hour time point. n=4 experimental replicates with 3 technical replicates/condition. P-values (one-way ANOVA), n.s. not significant.

**e.** Phase images of PC3 cells described in **(a-b)**. Scale bars, 100 μm. Representative of n=3 experimental replicates with 4 technical replicates/condition.

**f.** Schema, machine learning applied to classify and quantify 2D PC3 cells into three phenotypic categories. Upper panel, mem:Venus Scr shRNA (green), whole cell stain (WCS, red) and Hoechst (nuclei, blue). Lower panels, Round (blue), Spindle (green) and Spread (red). Scale bar, 100μm. Mean proportion of PC3 cells, expressing Scr shRNA, with each phenotype shown for n=3 experimental replicates each with 18 technical replicates. Total of 290,830 cells quantified.

**g-j.** PC3 cells in 2D expressing Scr and **(g)** *ARF1,* **(h)** *ARF3*, **(i)** *PSD* or **(j)** *RAB11FIP4* shRNA were classified into Round, Spindle and Spread. Heatmaps, Log_2_ Fold Change over Scr. P-values, one-way ANOVA, greyscale values as indicated. n=4, 2, 2 and 2 experimental replicates with 3-4 technical replicates/condition for *ARF1*, *ARF3*, *PSD* and *RAB11FIP4* shRNA respectively. **(g)** 15,392 (Scr), 26,486 (*ARF1_*KD2), 20,480 (*ARF1_*KD4), **(h)** 3,610 (Scr), 2,692 (*ARF3_*KD1), 2,882 (*ARF3_*KD2), **(i)** 6,769 (Scr), 16,953 (*PSD_*KD1), 24,041 (*PSD*_KD2) and **(j)** 18,432 (Scr), 30,568 (*RAB11FIP4_*KD1), 35,729 (*RAB11FIP4*_KD2) cells were quantified in total.

**k.** PC3 cells expressing mNG and Scr or *PSD* shRNA in 3D invasion assay. Yellow lines, initial wound and red pseudo colour, wound at t=Max_1/2._ Scale bars, 100μm. Magnified image of boxed region shown. White arrowhead, invasive chains. RWD at t=Max_1/2_ is shown, normalised to Scr. Data is mean ± s.e.m. (3 experimental replicates, triangles, 4-8 technical replicates, circles). P-values (Student’s 2-tailed t-test), *p 0.05, and ***p≤0.001.

**l.** PC3 cells expressing mNG and either Scr or *RAB11FIP4* shRNA in 3D invasion assay. Yellow lines, initial wound and red pseudo colour, wound at t=Max_1/4._ Scale bars, 100 μm. Magnified image of boxed region shown. White arrowhead, invasive chains. RWD at t=Max_1/4_ is shown, normalised to Scr. Data is mean ± s.e.m. (3 experimental replicates, triangles, 3-8 technical replicates, circles). P-values (Student’s 2-tailed t-test), ****p ≤ 0.0001.

**Supplementary Figure 4: ARF1 or ARF3 over-expression has different effects on migration and invasion of prostate cancer cells.**

**a.** Phase images of PC3 cells expressing mNG, ARF1-mNG or ARF3-mNG and Scr shRNA (upper panels). Scale bars, 100μ m. n=3 experimental replicates with 4 technical replicates/condition. Confocal images (middle panels) show localization of mNG constructs (black, inverted images) in 2D cells. Magnified images of boxed regions shown (a-f). White arrows, ARF mNG in puncta. Scale bars, 20μm. Images are representative of observations made in 3 experimental replicates. Also shown (lower panels) are phase images of PC3 acini, Round (red), Spindle (green) and Spread (blue). Scale bar, 100 μm. n=6 and 4 experimental replicates for ARF1-mNG and ARF3-mNG respectively each with 2-4 technical replicates/condition. Quantitation shown in **4e** and **4f**.

**b-c.** Confluence quantified in cells expressing mNG, **(b)** ARF1-mNG or **(c)** ARF3- mNG and Scr shRNA using phase images. Data is mean ± s.e.m., normalized to time 0. n=3 experimental replicates with 4 technical replicates/condition. P-values (one-way ANOVA), n.s. not significant.

**d-e.** PC3 acini expressing mNG, **(d)** ARF1-mNG or **(e)** ARF3-mNG and Scr shRNA were plated for 4 to 72 hours. CellTiter-Glow was added and luminescence measured to assess ATP-based cell viability. Data is mean ± s.e.m., normalized to 4 hour time point. n=4 experimental replicates with 3 technical replicates/condition. P-values (one-way ANOVA), n.s. not significant.

**f.** PC3 cells expressing mNG, ARF1-mNG or ARF3-mNG and Scr shRNA were classified into Round, Spindle and Spread. Heatmaps, Log_2_ Fold Change over mNG. P-values, one-way ANOVA, greyscale values as indicated. n=3 experimental replicates with 3-4 technical replicates/condition. 4,309 (mNG), 4,766 (ARF1) and 5,261 (mNG), 6,508 (ARF3) mNG-positive cells quantified in total.

**g-j.** Phase images of cells expressing mNG, **(g)** ARF1-mNG or **(i)** ARF3-mNG and Scr shRNA in 2D migration assay. Yellow lines, initial wound and red pseudo colour, wound at t=Max_1/2._ Scale bars, 100 μm. Magnified images of boxed regions shown. RWD at t=Max_1/2_, normalised to mNG is shown for **(h)** ARF1-mNG or **(j)** ARF3-mNG. Data is mean ± s.e.m. (4-5 experimental replicates, triangles, 3-4 technical replicates, circles). P-values (Student’s 2-tailed t-test), n.s. not significant and ***p ≤ 0.001.

**k.** Confocal images of PC3 cells expressing mNG, ARF3-mNG or ARF-mNG chimeras (black, inverted images in upper panels). Magnified images of boxed regions shown (a-h). White arrows, ARF mNG in discrete puncta, black arrows, areas of concentrated peri-nuclear staining. Scale bars, 20 μm. n=3 experimental replicates. Confocal images of PC3 acini stained with F-actin (red) and Hoechst (nuclei, blue) (middle panels). Intensity of F-actin staining can be appreciated using FIRE LUT. Scale bars, 20 μm. n=3 experimental replicates. Phase images of PC3 acini (lower panels); Round (red), Spindle (green) and Spread (blue). Scale bar, 100 μm n=3 experimental replicates each with 3 technical replicates/condition.

**l.** 2D PC3 cells expressing mNG, ARF3-mNG or ARF-mNG chimeras classified into Round, Spindle and Spread. Heatmaps, Log_2_ Fold Change over mNG. P-values, one-way ANOVA, greyscale values as indicated. n=2 experimental replicates with 4 technical replicates/condition. 4,746 (mNG), 7,342 (ARF3-mNG), 5,567 (3N/1C) and 4,717 (1N/3C) mNG-positive cells quantified in total.

**m.** Quantitation of **(k)** (upper panels). n=3 experimental replicates with 43 (ARF3), 97 (3N/1C), 76 (1N/3C) cells quantified in total. Graph is percentage of cells with mNG concentrated in the peri-nuclear region. Data is mean ± s.e.m. P-values (Student’s 2-tailed t-test), n.s. not significant and ***p ≤ 0.001.

**n.** Quantitation of **(k)** (lower panels). n=3 experimental replicates each with 3 technical replicates/condition. 3,765 (mNG), 2,218 (ARF3-mNG), 1,150 (3N/1C) and 3,067 (1N/3C) mNG-positive acini quantified in total. Heatmaps, Area is mean of Z-score normalised values (purple to yellow). P-values, Student’s t-test, Bonferroni adjustment, represented by size of bubble. Heatmaps, Round, Spindle or Spread is Log_2_ Fold Change from control (mNG) (blue to red). Proportion of control at each time is also Z-score normalised (white to black). P-values, Cochran-Mantel-Haenszel (CMH) test, Bonferroni adjusted, represented by size of bubble. Dot indicates p-value (Breslow-Day test, Bonferroni-adjusted) for consistent effect magnitude.

**Supplementary Figure 5: N-cadherin depletion mimics ARF3 depletion in 2D and 3D assays.**

**a.** Quantitation of total F-actin intensity/acini described in **(5a)**. Box and whiskers plot, 10–90 percentile; +, mean; dots, outliers; midline, median; boundaries, quartiles. n=2 experimental replicates with 643 (mNG, Scr shRNA), 712 (mNG, *ARF3_KD1* shRNA) and 383 (ARF3-mNG, Scr shRNA) cells quantified in total. P-values (Student’s 2-tailed t-test), ****p ≤ 0.0001.

**b.** Quantitation of percentage of PC3 acini described in **(5a)** with F-actin intensity visibly reduced in junctions. Data is mean ± s.e.m. n=3 experimental replicates with 121 (mNG, Scr shRNA), 108 (mNG, *ARF3_KD1* shRNA) and 121 (ARF3-mNG, Scr shRNA) cells quantified in total. P-values (Student’s 2-tailed t-test), n.s. not significant and ***p ≤ 0.001.

**c-f.** Representative western blots of PC3 cells expressing **(c)** mNG or **(d)** ARF3-mNG and Scr or *N-cadherin* shRNA for N-cadherin, E-cadherin and ARF3 antibodies. GAPDH is a loading control for both cadherin blots and a sample control for ARF3. Graphs are fold change, normalized to Scr. Data is mean ± s.e.m. for n=3 or 5 independent lysate preparations for mNG **(c,e)** and 3 independent preparations for ARF3-mNG **(d,f)**. P-values (Student’s 2-tailed t-test), n.s. not significant. *p ≤ 0.05, **p ≤ 0.01, ***p ≤ 0.001 and ****p ≤ 0.0001.

**g.** 2D PC3 cells expressing mNG or ARF3-mNG and Scr or *N-cadherin* shRNA were classified into Round, Spindle and Spread. Heatmaps, Log_2_ Fold Change over Scr. P-values, one-way ANOVA, greyscale values as indicated. n=3 experimental replicates with 3 technical replicates/condition. 19,184 (Scr), 37,230 (*N-cadherin* KD1), 26,284 (*N-cadherin* KD2) and 16,210 (Scr), 29,065 (*N-cadherin* KD1), 46,352 (*N-cadherin* KD2) cells were quantified for mNG or ARF3-mNG respectively.

**h.** Representative phase images of PC3 acini expressing mNG and Scr or *N-cadherin* shRNA. Outlines; Round (red), Spindle (green) and Spread (blue). Scale bar, 100μm n=5 experimental replicates each with 3-4 technical replicates/condition. 23,538 (Scr), 34,624 (*N-cadherin_*KD1) and 36,432 (*N-cadherin_*KD2) acini quantified in total.

**i.** Quantitation of **(h)**. Heatmaps, Area is mean of Z-score normalised values (purple to yellow). P-values, Student’s t-test, Bonferroni adjustment, represented by size of bubble. Heatmaps, Round, Spindle or Spread is Log_2_ Fold Change from control (Scr) (blue to red). Proportion of control at each time is Z-score normalised (white to black). P-values, Cochran-Mantel-Haenszel (CMH) test, Bonferroni adjusted, represented by size of bubble. Dot indicates p-value (Breslow-Day test, Bonferroni-adjusted) for consistent effect magnitude.

**j.** Western blot of PC3 cells expressing mNG and Scr or *RAB11FIP4* shRNA for N-cadherin and GAPDH, as a loading control. Graph is fold change, normalized to Scr. Data is mean ± s.e.m. Panels shown are representative of 3 independent lysate preparations. P-values (Student’s 2-tailed t-test), *p ≤ 1.5 and **p ≤ 0.01.

**Supplementary Figure 6: N-cadherin and ARF3 expression in prostate cancer patients.**

**a-d.** Analysis of Normal versus Tumour mRNA expression (Log2) for *ARF3* and *CDH2* from TCGA and GENT2 datasets (see methods for dataset IDs). Dots, expression per patient. Blue, normal; red, tumour. Circle next to Tumour Type indicates significant expression directionality change in tumour compared to normal tissue: Green, higher in tumour; magenta, lower in tumour. Patient numbers on graph. P-values, Brown-Forsythe and Welch ANOVA test with unpaired Welch’s correction, with individual variances compared for each comparison. *, 0.0332; **, 0.0021; ***, 0.0002; ****, <0.0001.

**Supplementary Table S1: ShRNA library information and result from screen.** ShRNA screen clone ID, The RNAi Consortium (TRC) clone ID, shRNA targeting region, Gene symbol, Gene ID, Species targeted, Reference sequence (RefSeq) ID, Validation Information (including methodology, target cell line and information, RNAi sequence (sense and antisense), match to human if designed to another species, the number of mem:Venus-positive acini quantified from the shRNA screen, and the Group ID assigned to the shRNA from the screen (‘Dendogram Group’).

**Supplementary Table S2: Non-screen RNAs used in this study.** ShRNA clone ID, The RNAi Consortium (TRC) clone ID, Gene target, and RNAi sequence (sense and antisense).

**Supplementary Table S3: Antibodies used in this study.** Antibodies utilised, supplier information, assay and concentration used.

**Supplementary Table S4: Combined normal and tumour dataset information for ARF3.** Tumour type, Dataset_ID, Normal Vs Tumour ID, and Sample count from GENT2 database for all combined expression data for ARF3 mRNA.

**Supplementary Table S5: Combined normal and tumour dataset information for CDH2.** Tumour type, Dataset_ID, Normal Vs Tumour ID, and Sample count from GENT2 database for all combined expression data for CDH2 mRNA.

**Supplementary Movie 1-2: Live imaging of acini from Fig 1e.** Live imaging in (Movie 1) phase and (Movie 2) mem:Venus of PC3 3D acini expressing control (Scr) shRNA from the mem:Venus pLKO.4 vector, imaged every hour for 96h.

**Supplementary Movie 3-5: Examples of alternate movement modalities from live imaging of wounded PC3 monolayers from Figure 3.** Live imaging in phase of PC3 monolayers displaying individual migration (Movie 3), sheet movement (Movie 4), and chain-lead movement (Movie 5), imaged every hour.

