## Supplementary material for "The small GTPase ARF3 controls metastasis and invasion modality by regulating N-cadherin levels": Supplemenary Table S1

| Screen ID | TRC ID | Region | Gene Symbol | Gene ID | Species | RefSeq Id | Validated | Cell line | Method | KD | Target Seq (sense) | Target Seq (anti-sense) | Human Match? | No. of acini quantified | Dendrogram Group |
|---|---|---|---|---|---|---|---|---|---|---|---|---|---|---|---|
| ACAP1_1 | TRCN0000122969 | CDS | ACAP1 | 9744 | Human | NM_014716 | | | | | GCTGATGTCAACTGGGTCAAT | ATTGACCCAGTTGACATCAGC | | 6434 | 4 |
| ACAP1_2 | TRCN0000122970 | CDS | ACAP1 | 9744 | Human | NM_014716 | | | | | GCAGGAGATGAGACGTATCTT | AAGATACGTCTCATCTCCTGC | | 11180 | 5 |
| ACAP1_3 | TRCN0000122971 | CDS | ACAP1 | 9744 | Human | NM_014716 | | | | | TCACGCTAAATACGTGGAGAA | TTCTCCACGTATTTAGCGTGA | | 6028 | 1 |
| ACAP1_4 | TRCN0000122972 | CDS | ACAP1 | 9744 | Human | NM_014716 | | | | | GCGCCTTCGTTGTCGGCATTT | AAATGCCGACAACGAAGGCGC | | 9009 | 4 |
| ACAP1_5 | TRCN0000122973 | CDS | ACAP1 | 9744 | Human | NM_014716 | | | | | CTGGTTCACCATTCAGAGCAA | TTGCTCTGAATGGTGAACCAG | | 7185 | 6 |
| ACAP2_2 | TRCN0000123038 | CDS | ACAP2 | 23527 | Human | NM_012287 | Yes | MCH58 | SYBR | 0.92 | CCAGTATTGCTACTGCTTATA | TATAAGCAGTAGCAATACTGG | | 8795 | 6 |
| ACAP3_1 | TRCN0000156390 | CDS | ACAP3 | 116983 | Human | NM_030649 | Yes | MCH58 | SYBR | 0.78 | CGATGAGTCCAAAGTGGAGTT | AACTCCACTTTGGACTCATCG | | 8083 | 5 |
| ADAP1_1 | TRCN0000148189 | CDS | ADAP1 | 11033 | Human | NM_006869 | | | | | GCTCTGAAGTATTTCAACAGA | TCTGTTGAAATACTTCAGAGC | | 12466 | 4 |
| ADAP1_2 | TRCN0000148260 | CDS | ADAP1 | 11033 | Human | NM_006869 | | | | | GCACTTCAAGCATAAACCTTA | TAAGGTTTATGCTTGAAGTGC | | 7494 | 7 |
| ADAP1_4 | TRCN0000181086 | CDS | ADAP1 | 11033 | Human | NM_006869 | | | | | CAGCACCCGTAACATCTTCAT | ATGAAGATGTTACGGGTGCTG | | 10315 | 7 |
| ADAP1_5 | TRCN0000446284 | 3UTR | ADAP1 | 11033 | Human | NM_006869 | | | | | ACGGACATTGGACTCACTGTG | CACAGTGAGTCCAATGTCCGT | | 8994 | 4 |
| ADAP2_1 | TRCN0000141961 | CDS | ADAP2 | 55803 | Human | NM_018404 | Yes | 293T/17 | SYBR | 0.88 | GCCTGGTCTTAAAGGAACAAT | ATTGTTCCTTTAAGACCAGGC | | 10462 | 7 |
| AGAP1_2 | TRCN0000158851 | 3UTR | AGAP1 | 116987 | Human | NM_014914 | Yes | 293T/17 | SYBR | 0.77 | GCTGTTTAATAGAACGTGATT | AATCACGTTCTATTAAACAGC | | 9586 | 7 |
| AGAP11_1 | TRCN0000253876 | CDS | AGAP11 | 119385 | Human | NM_133447 | | | | | GGACATGGAAGCTCTGCATAT | ATATGCAGAGCTTCCATGTCC | | 10426 | 5 |
| AGAP11_2 | TRCN0000253877 | CDS | AGAP11 | 119385 | Human | NM_133447 | | | | | ACATCATCATATCACGGAAAG | CTTTCCGTGATATGATGATGT | | 7781 | 4 |
| AGAP11_3 | TRCN0000253878 | CDS | AGAP11 | 119385 | Human | NM_133447 | | | | | ACCAGCCAGGAGGACCTTTAT | ATAAAGGTCCTCCTGGCTGGT | | 15378 | 6 |
| AGAP11_4 | TRCN0000253879 | CDS | AGAP11 | 119385 | Human | NM_133447 | | | | | CATACTTGATGAACAGTTATA | TATAACTGTTCATCAAGTATG | | 9256 | 7 |
| AGAP11_5 | TRCN0000265447 | CDS | AGAP11 | 119385 | Human | NM_133447 | | | | | TGGCCAAACGTGCCACTTTAA | TTAAAGTGGCACGTTTGGCCA | | 12210 | 7 |
| AGAP2_1 | TRCN0000072941 | CDS | AGAP2 | 116986 | Human | NM_014770 | Yes | A3 | SYBR | 0.75 | CTATGAGACTTGTGCAACCTA | TAGGTTGCACAAGTCTCATAG | | 13926 | 4 |
| AGAP3_1 | TRCN0000047999 | CDS | AGAP3 | 116988 | Human | NM_031946 | | | | | GCAGACATCTTGATCCAGCAT | ATGCTGGATCAAGATGTCTGC | | 9532 | 6 |
| AGAP3_2 | TRCN0000048000 | CDS | AGAP3 | 116988 | Human | NM_031946 | | | | | GCATCCCAATATCTACGCCAT | ATGGCGTAGATATTGGGATGC | | 11429 | 4 |
| AGAP3_3 | TRCN0000438364 | CDS | AGAP3 | 116988 | Human | NM_031946 | | | | | GGATACGGGCCAAGTATGAAC | GTTCATACTTGGCCCGTATCC | | 8746 | 3 |
| AGAP3_4 | TRCN0000438806 | CDS | AGAP3 | 116988 | Human | NM_031946 | | | | | ACATCAACCAGGCCACGAATG | CATTCGTGGCCTGGTTGATGT | | 9478 | 7 |
| AGAP3_5 | TRCN0000444944 | CDS | AGAP3 | 213990 | Mouse | NM_139153 | | | | | GTTCAGCCTGGAGGATGAAAT | ATTTCATCCTCCAGGCTGAAC | Y | 11257 | 6 |
| AGFG1_1 | TRCN0000060223 | CDS | AGFG1 | 3267 | Human | NM_004504 | Yes | A549 | SYBR | 0.99 | CCTGAGGTCAAACCACTGAAA | TTTCAGTGGTTTGACCTCAGG | | 11014 | 5 |
| AGFG2_1 | TRCN0000147372 | CDS | AGFG2 | 3268 | Human | NM_006076 | Yes | A549 | SYBR | 0.76 | GTCAATCTCCATGACAACTTT | AAAGTTGTCATGGAGATTGAC | | 15969 | 4 |
| AP1B1_1 | TRCN0000065133 | CDS | AP1B1 | 162 | Human | NM_001127 | Yes | PC3 | WB | | CCACACTGTTAGACGAGCTTA | TAAGCTCGTCTAACAGTGTGG | | 7253 | 6 |
| AP1B1_2 | TRCN0000065134 | CDS | AP1B1 | 162 | Human | NM_001127 | Yes | PC3 | WB | | CGTGAAGTACAACGACCCTAT | ATAGGGTCGTTGTACTTCACG | | 14582 | 6 |
| AP1B1_3 | TRCN0000065135 | CDS | AP1B1 | 162 | Human | NM_001127 | Yes | PC3 | WB | | GCCGTGAAGAACAACATCGAT | ATCGATGTTGTTCTTCACGGC | | 15705 | 3 |
| AP1B1_4 | TRCN0000065136 | CDS | AP1B1 | 162 | Human | NM_001127 | Yes | PC3 | WB | | CCATTGTGAAACTCTTTCTAA | TTAGAAAGAGTTTCACAATGG | | 10179 | 6 |
| AP1B1_5 | TRCN0000065137 | CDS | AP1B1 | 162 | Human | NM_001127 | Yes | PC3 | WB | | CGTTGACAAGATCACAGAGTA | TACTCTGTGATCTTGTCAACG | | 15518 | 6 |
| AP1G1_1 | TRCN0000293941 | CDS | AP1G1 | 164 | Human | NM_001128 | Yes | A549 | SYBR | 0.97 | TTGCGAGTCCTAGCCATAAAT | ATTTATGGCTAGGACTCGCAA | | 11551 | 4 |
| AP2M1_2 | TRCN0000060241 | CDS | AP2M1 | 1173 | Human | NM_004068 | Yes | 293T/17 | SYBR | 0.95 | GCTGGATGAGATTCTAGACTT | AAGTCTAGAATCTCATCCAGC | | 7298 | 1 |
| AP3B1_1 | TRCN0000286136 | CDS | AP3B1 | 8546 | Human | NM_003664 | Yes | A549 | SYBR | 0.92 | GCAAGTATTCTTTGGCTAATT | AATTAGCCAAAGAATACTTGC | | 9237 | 7 |
| AP3D1_2 | TRCN0000293891 | CDS | AP3D1 | 8943 | Human | NM_003938 | Yes | A549 | SYBR | 0.98 | AGTTCACCTTCAAGCGAATTG | CAATTCGCTTGAAGGTGAACT | | 8936 | 4 |
| AP4M1_1 | TRCN0000059718 | CDS | AP4M1 | 9179 | Human | NM_004722 | Yes | A3 | SYBR | 0.81 | GCTCCAGGTTTATCTAAAGTT | AACTTTAGATAAACCTGGAGC | | 9560 | 6 |
| ARAP1_1 | TRCN0000047835 | CDS | ARAP1 | 116985 | Human | NM_015242 | Yes | MCH58 | SYBR | 0.55 | GTGGCCTATTAAGAGTCTCAA | TTGAGACTCTTAATAGGCCAC | | 10248 | 1 |
| ARAP2_1 | TRCN0000149274 | 3UTR | ARAP2 | 116984 | Human | NM_015230 | | | | | GCACTTTGAGAAACCTTGTAT | ATACAAGGTTTCTCAAAGTGC | | 9635 | 4 |
| ARAP2_2 | TRCN0000149503 | CDS | ARAP2 | 116984 | Human | NM_015230 | | | | | GCTATCATTGAACACCTGTAT | ATACAGGTGTTCAATGATAGC | | 14015 | 6 |
| ARAP2_3 | TRCN0000149818 | 3UTR | ARAP2 | 116984 | Human | NM_015230 | | | | | CCAGAACTTGTCAATGAGAAA | TTTCTCATTGACAAGTTCTGG | | 16613 | 6 |
| ARAP2_4 | TRCN0000182989 | CDS | ARAP2 | 116984 | Human | NM_015230 | | | | | CATCTAAATCTAGGACTCAAA | TTTGAGTCCTAGATTTAGATG | | 10437 | 6 |
| ARAP2_5 | TRCN0000183746 | 3UTR | ARAP2 | 116984 | Human | NM_015230 | | | | | CCTTGTAAAGTTCCATTGTAT | ATACAATGGAACTTTACAAGG | | 12188 | 4 |
| ARAP3_1 | TRCN0000047344 | CDS | ARAP3 | 64411 | Human | NM_022481 | Yes | A549 | SYBR | 0.71 | GTCTGGAGTAATGAGATAGTA | TACTATCTCATTACTCCAGAC | | 9119 | 2 |
| ARF1_1 | TRCN0000039876 | CDS | ARF1 | 375 | Human | NM_001658 | Yes | MCF7 | SYBR | 0.97 | CAATGACAGAGAGCGTGTGAA | TTCACACGCTCTCTGTCATTG | | 9163 | 1 |
| ARF1_2 | TRCN0000039875 | CDS | ARF1 | 375 | Human | NM_001658 | Yes | MCF7 | SYBR | 0.97 | CCTCCTGGTGTTCGCCAACAA | TTGTTGGCGAACACCAGGAGG | | 7914 | 5 |
| ARF3_1 | TRCN0000047674 | CDS | ARF3 | 377 | Human | NM_001659 | Yes | A549 | SYBR | 0.7 | GATGCTGTACTCCTTGTCTTT | AAAGACAAGGAGTACAGCATC | | 8964 | 1 |
| ARF3_2 | TRCN0000047676 | CDS | ARF3 | 377 | Human | NM_001659 | Yes | A549 | SYBR | 0.62 | GAACACCCAAGGGTTGATATT | AATATCAACCCTTGGGTGTTC | | 8826 | 5 |
| ARF4_1 | TRCN0000047940 | CDS | ARF4 | 378 | Human | NM_001660 | Yes | A549 | SYBR | 0.98 | CATGGTATGTTCAAGCCACTT | AAGTGGCTTGAACATACCATG | | 15289 | 7 |
| ARF4_2 | TRCN0000293758 | 3UTR | ARF4 | 378 | Human | NM_001660 | Yes | A549 | SYBR | 0.92 | TTACTTCCCATAAGCGTAAAT | ATTTACGCTTATGGGAAGTAA | | 7615 | 7 |
| ARF5_1 | TRCN0000047991 | CDS | ARF5 | 381 | Human | NM_001662 | Yes | A549 | SYBR | 0.99 | GATGCAGTGCTGCTGGTATTT | AAATACCAGCAGCACTGCATC | | 11018 | 7 |
| ARF5_2 | TRCN0000294062 | 3UTR | ARF5 | 381 | Human | NM_001662 | Yes | A549 | SYBR | 0.9 | TGCATGTTCTCTCTGTTGTTG | CAACAACAGAGAGAACATGCA | | 9880 | 4 |
| ARF6_6 | TRCN0000294069 | 3UTR | ARF6 | 382 | Human | NM_001663 | Yes | A549 | SYBR | 0.74 | CTTGCTGTAGATGGCTTATTT | AAATAAGCCATCTACAGCAAG | | 10096 | 1 |
| ARFGAP1_2 | TRCN0000293964 | 3UTR | ARFGAP1 | 55738 | Human | NM_175609 | Yes | A549 | SYBR | 0.8 | GCCAGTTCACTACGCAGTATC | GATACTGCGTAGTGAACTGGC | | 9224 | 7 |
| ARFGAP2_1 | TRCN0000242598 | CDS | ARFGAP2 | 84364 | Human | NM_032389 | Yes | MCH58 | SYBR | 0.84 | TTTGACGATGTTGGTACTTTC | GAAAGTACCAACATCGTCAAA | | 8613 | 5 |

| | | | | | | | | | | | | | | | |
|---|---|---|---|---|---|---|---|---|---|---|---|---|---|---|---|
| ARFGAP3_2 | TRCN0000047337 | CDS | ARFGAP3 | 26286 | Human | NM_014570 | Yes | HeLa | SYBR | 0.94 | GCCAAGGTGGTATCTAAAGAA | TTCTTTAGATACCACCTTGGC | | 7781 | 1 |
| ARFIP1_1 | TRCN0000061914 | CDS | ARFIP1 | 27236 | Human | NM_014447 | Yes | A549 | SYBR | 0.98 | CCAGTTATTCTAGCAGATGAA | TTCATCTGCTAGAATAACTGG | | 10651 | 4 |
| ARFIP2_1 | TRCN0000061938 | CDS | ARFIP2 | 23647 | Human | NM_012402 | Yes | A549 | SYBR | 0.94 | GCAACTGTTATCAGAACGATT | AATCGTTCTGATAACAGTTGC | | 10478 | 5 |
| ARHGAP21_2 | TRCN0000156175 | CDS | ARHGAP21 | 57584 | Human | NM_020824 | Yes | A549 | SYBR | 0.92 | CCTCACAACTTCAGCTCCATT | AATGGAGCTGAAGTTGTGAGG | | 10519 | 4 |
| ARRB1_1 | TRCN0000230149 | CDS | ARRB1 | 408 | Human | NM_004041 | Yes | A549 | SYBR | 0.93 | GAACTGCCCTTCACCCTAATG | CATTAGGGTGAAGGGCAGTTC | | 12863 | 6 |
| ARRB2_1 | TRCN0000280685 | CDS | ARRB2 | 409 | Human | NM_004313 | Yes | A549 | SYBR | 0.95 | CTTCGTAGATCACCTGGACAA | TTGTCCAGGTGATCTACGAAG | | 10951 | 4 |
| ASAP1_2 | TRCN0000286562 | CDS | ASAP1 | 50807 | Human | NM_018482 | Yes | A549 | SYBR | 0.79 | CCAGGGATTTACTTGCACTAA | TTAGTGCAAGTAAATCCCTGG | | 11331 | 5 |
| ASAP2_1 | TRCN0000029749 | CDS | ASAP2 | 8853 | Human | NM_003887 | | | | | GCCTCAAACCTTCCATTGAAA | TTTCAATGGAAGGTTTGAGGC | | 9879 | 5 |
| ASAP2_2 | TRCN0000029750 | CDS | ASAP2 | 8853 | Human | NM_003887 | | | | | CCTGGATAAACAGACAGGGAA | TTCCCTGTCTGTTTATCCAGG | | 9666 | 7 |
| ASAP2_3 | TRCN0000029752 | CDS | ASAP2 | 8853 | Human | NM_003887 | | | | | CCCTTTGGACAGTTTGCTGAA | TTCAGCAAACTGTCCAAAGGG | | 10190 | 7 |
| ASAP2_4 | TRCN0000029753 | CDS | ASAP2 | 8853 | Human | NM_003887 | | | | | CCGGTGTCATTTGTGCACTTT | AAAGTGCACAAATGACACCGG | | 8649 | 2 |
| ASAP3_1 | TRCN0000151669 | CDS | ASAP3 | 55616 | Human | NM_017707 | Yes | MCH58 | SYBR | 0.75 | GACTACATTATGGCCAAGTAT | ATACTTGGCCATAATGTAGTC | | 7301 | 4 |
| ATP6V0C_2 | TRCN0000029561 | CDS | ATP6V0C | 527 | Human | NM_001694 | Yes | A549 | SYBR | 0.98 | GCTCGGCCTCTACGGTCTCAT | ATGAGACCGTAGAGGCCGAGC | | 8542 | 4 |
| BIG1_2 | TRCN0000048281 | CDS | ARFGEF1 | 10565 | Human | NM_006421 | Yes | A549 | SYBR | 0.91 | CCAGAACAACAGACAGAGAAA | TTTCTCTGTCTGTTGTTCTGG | | 7788 | 4 |
| BIG2_1 | TRCN0000251882 | CDS | ARFGEF2 | 99371 | Mouse | NM_001085495 | Yes | Hepa 1-6 | SYBR | 0.7 | CATTGAAGCTGACAAGTATTT | AAATACTTGTCAGCTTCAATG | Y | 15384 | 7 |
| BIG3_2 | TRCN0000144294 | CDS | ARFGEF3 | 57221 | Human | NM_020340 | Yes | A549 | SYBR | 0.89 | CCAACTCTGGATTTACTCTTT | AAAGAGTAAATCCAGAGTTGG | | 8431 | 1 |
| CERT_1 | TRCN0000382287 | CDS | COL4A3BP | 10087 | Human | NM_005713 | Yes | A549 | SYBR | 0.78 | GCTTCTCAGCGAGACGTATTA | TAATACGTCTCGCTGAGAAGC | | 12980 | 5 |
| COPB1_2 | TRCN0000380811 | CDS | COPB1 | 1315 | Human | NM_016451 | Yes | A549 | SYBR | 0.86 | TGCTGTTACCGGCCATATAAG | CTTATATGGCCGGTAACAGCA | | 8256 | 7 |
| COPG1_2 | TRCN0000380046 | CDS | COPG | 22820 | Human | NM_016128 | Yes | MCH58 | SYBR | 0.97 | GTCAGACAAAGTGCCGGATAA | TTATCCGGCACTTTGTCTGAC | | 10932 | 4 |
| COPG2_1 | TRCN0000154575 | CDS | COPG2 | 26958 | Human | NM_012133 | Yes | A3 | SYBR | 0.96 | GCCATTACTACACTCCTCAAA | TTTGAGGAGTGTAGTAATGGC | | 9247 | 7 |
| CYTH1_2 | TRCN0000230431 | CDS | CYTH1 | 9267 | Human | NM_004762 | Yes | A549 | SYBR | 0.86 | AGGCGTTTGCCCAGCGATATT | AATATCGCTGGGCAAACGCCT | | 7286 | 2 |
| CYTH2_1 | TRCN0000062099 | CDS | CYTH2 | 9266 | Human | NM_017457 | Yes | MCH58 | SYBR | 0.76 | GCGGATTTCAGTCAAGAAGAA | TTCTTCTTGACTGAAATCCGC | | 9407 | 4 |
| CYTH3_1 | TRCN0000179183 | CDS | CYTH3 | 9265 | Human | NM_004227 | Yes | A549 | SYBR | 0.89 | GATGACATTGAGAGGCTGAAA | TTTCAGCCTCTCAATGTCATC | | 9885 | 7 |
| CYTH4_1 | TRCN0000180672 | 3UTR | CYTH4 | 27128 | Human | NM_013385 | | | | | GCCACAGACATCATTGCTGTT | AACAGCAATGATGTCTGTGGC | | 6901 | 5 |
| CYTH4_2 | TRCN0000180673 | 3UTR | CYTH4 | 27128 | Human | NM_013385 | | | | | GAAGCCAAGAAGGGAGAGTTT | AAACTCTCCCTTCTTGGCTTC | | 9046 | 7 |
| CYTH4_3 | TRCN0000242586 | CDS | CYTH4 | 27128 | Human | NM_013385 | | | | | TTGCACGGTTCCTGTATAAAG | CTTTATACAGGAACCGTGCAA | | 12274 | 4 |
| CYTH4_4 | TRCN0000242587 | CDS | CYTH4 | 27128 | Human | NM_013385 | | | | | CCGCCAAGGGTATCCAGTATT | AATACTGGATACCCTTGGCGG | | 7916 | 7 |
| CYTH4_5 | TRCN0000242588 | 3UTR | CYTH4 | 27128 | Human | NM_013385 | | | | | CAAGCAGGATGGGTGCTATAT | ATATAGCACCCATCCTGCTTG | | 10668 | 4 |
| ELMOD1_1 | TRCN0000122881 | CDS | ELMOD1 | 55531 | Human | NM_018712 | Yes | 293T/17 | SYBR | 0.88 | GCTTGCCTTCTGCAAATCGTT | AACGATTTGCAGAAGGCAAGC | | 12697 | 1 |
| ELMOD2_1 | TRCN0000128010 | CDS | ELMOD2 | 255520 | Human | NM_153702 | Yes | A549 | SYBR | 0.91 | GAAGTGTGAATTGCAGCGAAT | ATTCGCTGCAATTCACACTTC | | 15799 | 7 |
| ELMOD3_1 | TRCN0000135161 | 3UTR | ELMOD3 | 84173 | Human | NM_032213 | | | | | CGAGATGAATGGAGGCTTAAA | TTTAAGCCTCCATTCATCTCG | | 12497 | 4 |
| ELMOD3_2 | TRCN0000135916 | CDS | ELMOD3 | 84173 | Human | NM_032213 | | | | | GAGTTGAAGAACCATGGCATT | AATGCCATGGTTCTTCAACTC | | 18107 | 4 |
| ELMOD3_3 | TRCN0000137628 | CDS | ELMOD3 | 84173 | Human | NM_032213 | | | | | GCAGCTAATCTCCTTCAGTGA | TCACTGAAGGAGATTAGCTGC | | 10585 | 7 |
| ELMOD3_4 | TRCN0000138508 | 3UTR | ELMOD3 | 84173 | Human | NM_032213 | | | | | CCGAGATGAATGGAGGCTTAA | TTAAGCCTCCATTCATCTCGG | | 13353 | 4 |
| ELMOD3_5 | TRCN0000138843 | 3UTR | ELMOD3 | 84173 | Human | NM_032213 | | | | | GCACCAGTGTCTGTGTTGTTA | TAACAACACAGACACTGGTGC | | 9706 | 4 |
| EXOC5_1 | TRCN0000061964 | CDS | EXOC5 | 10640 | Human | NM_006544 | Yes | 293T/17 | SYBR | 0.89 | GCCAGCTGATTCAGGAGTTTA | TAAACTCCTGAATCAGCTGGC | | 8921 | 4 |
| FAPP1_1 | TRCN0000147627 | CDS | PLEKHA3 | 65977 | Human | NM_019091 | Yes | A549 | SYBR | 0.91 | GAATGTGTGAAGATAGCCAAT | ATTGGCTATCTTCACACATTC | | 10204 | 4 |
| FBX8_2 | TRCN0000034316 | CDS | FBXO8 | 26269 | Human | NM_012180 | Yes | MCH58 | SYBR | 0.82 | GAGAGTATCTTGAAACTCTTA | TAAGAGTTTCAAGATACTCTC | | 13889 | 6 |
| GBF1_1 | TRCN0000154341 | CDS | GBF1 | 8729 | Human | NM_004193 | Yes | 293T/17 | SYBR | 0.84 | CTGCATAGTTTCGGTCATCTA | TAGATGACCGAAACTATGCAG | | 9173 | 1 |
| GGA1_1 | TRCN0000381215 | CDS | GGA1 | 26088 | Human | NM_001001560 | Yes | PC3 | WB | | GGACATCGGAGAAGGTGAAGA | TCTTCACCTTCTCCGATGTCC | | 10805 | 5 |
| GGA1_2 | TRCN0000381323 | CDS | GGA1 | 26088 | Human | NM_001001560 | | | | | AGATCTCGAAGAGGGTGAATG | CATTCACCCTCTTCGAGATCT | | 10877 | 7 |
| GGA1_3 | TRCN0000381482 | 3UTR | GGA1 | 26088 | Human | NM_001001560 | Yes | PC3 | WB | | TGTAGGCCTCAAGTGACTCTT | AAGAGTCACTTGAGGCCTACA | | 13018 | 6 |
| GGA1_4 | TRCN0000381503 | CDS | GGA1 | 26088 | Human | NM_001001560 | Yes | PC3 | WB | | AGAGGCCTACCAGATGCTAAA | TTTAGCATCTGGTAGGCCTCT | | 9854 | 5 |
| GGA1_5 | TRCN0000382063 | CDS | GGA1 | 26088 | Human | NM_001001560 | | | | | AGTGGGCAAGTTCCGCTTTCT | AGAAAGCGGAACTTGCCCACT | | 8547 | 7 |
| GGA2_2 | TRCN0000065006 | CDS | GGA2 | 23062 | Human | NM_015044 | Yes | A549 | SYBR | 0.8 | CCGTGCCCTCTGAATTATGTT | AACATAATTCAGAGGGCACGG | | 8153 | 4 |
| GGA3_1 | TRCN0000232888 | CDS | GGA3 | 23163 | Human | NM_014001 | Yes | A549 | SYBR | 0.66 | GAAGGTGCGGCTTCGGTATAA | TTATACCGAAGCCGCACCTTC | | 11254 | 4 |
| GIT1_1 | TRCN0000346581 | CDS | GIT1 | 216963 | Mouse | NM_001004144 | Yes | Hepa 1-6 | SYBR | 0.79 | CCACCTTGATCATCGACATTC | GAATGTCGATGATCAAGGTGG | Y | 10197 | 7 |
| GIT2_1 | TRCN0000364545 | CDS | GIT2 | 9815 | Human | NM_057169 | Yes | A549 | SYBR | 0.91 | ATAACGGTGCTAACTCTATAT | ATATAGAGTTAGCACCGTTAT | | 13481 | 7 |
| GOSR2_2 | TRCN0000115095 | CDS | GOSR2 | 56494 | Mouse | NM_019650 | Yes | Hepa 1-6 | SYBR | 0.93 | TCAGAAGAAGATCCTTGACAT | ATGTCAAGGATCTTCTTCTGA | Y | 12093 | 7 |
| HERC1_1 | TRCN0000235499 | CDS | HERC1 | 8925 | Human | NM_003922 | Yes | A549 | SYBR | 0.78 | GTTCGAACTGGTCTAAGTTTA | TAAACTTAGACCAGTTCGAAC | | 7996 | 7 |
| IKBKG_1 | TRCN0000022148 | CDS | IKBKG | 8517 | Human | NM_003639 | Yes | A3 | SYBR | 0.59 | GAGGGAGTACAGCAAACTGAA | TTCAGTTTGCTGTACTCCCTC | | 11425 | 5 |
| IL6_1 | TRCN0000059205 | CDS | IL6 | 3569 | Human | NM_000600 | | | | | CATCTCATTCTGCGCAGCTTT | AAAGCTGCGCAGAATGAGATG | | 12652 | 4 |
| IL6_2 | TRCN0000059206 | CDS | IL6 | 3569 | Human | NM_000600 | | | | | GACATGTAACAAGAGTAACAT | ATGTTACTCTTGTTACATGTC | | 10831 | 5 |
| IL6_3 | TRCN0000059207 | CDS | IL6 | 3569 | Human | NM_000600 | | | | | CAGAACGAATTGACAAACAAA | TTTGTTTGTCAATTCGTTCTG | | 8076 | 4 |
| IL6_4 | TRCN0000372667 | 3UTR | IL6 | 3569 | Human | NM_000600 | | | | | ATGAGCGTTAGGACACTATTT | AAATAGTGTCCTAACGCTCAT | | 7701 | 5 |

| | | | | | | | | | | | | | | | |
|---|---|---|---|---|---|---|---|---|---|---|---|---|---|---|---|
| IL6_5 | TRCN0000372668 | 3UTR | IL6 | 3569 | Human | NM_000600 | | | | | ATGTGAAGCTGAGTTAATTTA | TAAATTAACTCAGCTTCACAT | | 12864 | 7 |
| IL6R_1 | TRCN0000058779 | CDS | IL6R | 3570 | Human | NM_000565 | | | | | CCACTCCTGGAACTCATCTTT | AAAGATGAGTTCCAGGAGTGG | | 9840 | 5 |
| IL6R_2 | TRCN0000058780 | CDS | IL6R | 3570 | Human | NM_000565 | | | | | CTCTGGAAACTATTCATGCTA | TAGCATGAATAGTTTCCAGAG | | 11663 | 5 |
| IL6R_3 | TRCN0000058781 | CDS | IL6R | 3570 | Human | NM_000565 | | | | | AGCCCTTATGACATCAGCAAT | ATTGCTGATGTCATAAGGGCT | | 9085 | 4 |
| IL6R_4 | TRCN0000058782 | CDS | IL6R | 3570 | Human | NM_000565 | | | | | CTCCTCTGCATTGCCATTGTT | AACAATGGCAATGCAGAGGAG | | 12006 | 5 |
| IL6ST_1 | TRCN0000058287 | CDS | IL6ST | 3572 | Human | NM_002184 | Yes | A549 | SYBR | 0.96 | CGGCCAGAAGATCTACAATTA | TAATTGTAGATCTTCTGGCCG | | 14708 | 3 |
| IQSEC1_3 | TRCN0000149750 | CDS | IQSEC1 | 9922 | Human | NM_014869 | Yes | PC3 | WB | | GATCTATGAACGGATCCGTAA | TTACGGATCCGTTCATAGATC | | 11780 | 5 |
| IQSEC2_1 | TRCN0000144705 | CDS | IQSEC2 | 23096 | Human | NM_015075 | Yes | A549 | SYBR | 0.81 | GCTGAACGAAAGATGAAACTA | TAGTTTCATCTTTCGTTCAGC | | 7971 | 5 |
| IQSEC3_1 | TRCN0000147813 | CDS | IQSEC3 | 440073 | Human | NM_015232 | | | | | GCTCTTTGAGAACGAGTATTA | TAATACTCGTTCTCAAAGAGC | | 9816 | 4 |
| IQSEC3_2 | TRCN0000179835 | CDS | IQSEC3 | 440073 | Human | NM_015232 | | | | | CGAGTATTACTCTCATGGCAT | ATGCCATGAGAGTAATACTCG | | 9730 | 7 |
| IQSEC3_3 | TRCN0000180343 | CDS | IQSEC3 | 440073 | Human | NM_015232 | | | | | GAAGTCCATTGTGGGCATGAA | TTCATGCCCACAATGGACTTC | | 10311 | 7 |
| IQSEC3_4 | TRCN0000180674 | CDS | IQSEC3 | 440073 | Human | NM_015232 | | | | | GCTCAGCAAGAACTTCGAGAA | TTCTCGAAGTTCTTGCTGAGC | | 9791 | 7 |
| IQSEC3_5 | TRCN0000180949 | CDS | IQSEC3 | 440073 | Human | NM_015232 | | | | | GAGGACTTCATCCGAAACCTT | AAGGTTTCGGATGAAGTCCTC | | 16208 | 4 |
| ITSN1_1 | TRCN0000002009 | CDS | ITSN1 | 6453 | Human | NM_003024 | Yes | A549 | SYBR | 0.64 | GCACTAGCTGACATGAATAAT | ATTATTCATGTCAGCTAGTGC | | 16401 | 4 |
| JIP4_1 | TRCN0000281124 | CDS | SPAG9 | 9043 | Human | NM_003971 | Yes | A549 | SYBR | 0.94 | CCCGAGTGGAATCTTTAGAAT | ATTCTAAAGATTCCACTCGGG | | 10283 | 2 |
| KDELR1_2 | TRCN0000054458 | CDS | KDELR1 | 68137 | Mouse | NM_133950 | Yes | Hepa 1-6 | SYBR | 0.98 | CTGCGATTTCTTCTACCTCTA | TAGAGGTAGAAGAAATCGCAG | Y | 12036 | 3 |
| MAPK8IP3_1 | TRCN0000037984 | CDS | MAPK8IP3 | 23162 | Human | NM_015133 | | | | | GCCTTGAATGTGGTGAAGAAT | ATTCTTCACCACATTCAAGGC | | 8785 | 7 |
| MAPK8IP3_2 | TRCN0000037985 | CDS | MAPK8IP3 | 23162 | Human | NM_015133 | | | | | CCATGACTCCACAGAAGTCAT | ATGACTTCTGTGGAGTCATGG | | 7062 | 4 |
| MAPK8IP3_3 | TRCN0000037986 | CDS | MAPK8IP3 | 23162 | Human | NM_015133 | | | | | GCACATTGAGAGGTCCAAGAT | ATCTTGGACCTCTCAATGTGC | | 15256 | 4 |
| MAPK8IP3_4 | TRCN0000037987 | CDS | MAPK8IP3 | 23162 | Human | NM_015133 | | | | | GCGGAGGATGTAAGCAGCTAT | ATAGCTGCTTACATCCTCCGC | | 13648 | 5 |
| MAPK8IP3_5 | TRCN0000037988 | CDS | MAPK8IP3 | 23162 | Human | NM_015133 | | | | | TCTGAGCAACTATCACCTAAT | ATTAGGTGATAGTTGCTCAGA | | 9943 | 7 |
| MICALL1_1 | TRCN0000062133 | 3UTR | MICALL1 | 85377 | Human | NM_033386 | | | | | CCAAAGAATCTCTTGTTTCTT | AAGAAACAAGAGATTCTTTGG | | 9401 | 4 |
| MICALL1_5 | TRCN0000414130 | CDS | MICALL1 | 85377 | Human | NM_033386 | | | | | AGGACAATGTCTTCGAGAATA | TATTCTCGAAGACATTGTCCT | | 13001 | 6 |
| NCS1_1 | TRCN0000427733 | 3UTR | NCS1 | 23413 | Human | NM_014286 | | | | | GGCTACAGCCCTCTGCATAAA | TTTATGCAGAGGGCTGTAGCC | | 9500 | 7 |
| NCS1_2 | TRCN0000055733 | CDS | NCS1 | 23413 | Human | NM_014286 | | | | | CCCACCAAGTTTGCCACATTT | AAATGTGGCAAACTTGGTGGG | | 11867 | 7 |
| NCS1_3 | TRCN0000055734 | CDS | NCS1 | 23413 | Human | NM_014286 | | | | | GCGAATTGAGTTCTCCGAGTT | AACTCGGAGAACTCAATTCGC | | 10709 | 6 |
| NCS1_5 | TRCN0000055736 | CDS | NCS1 | 23413 | Human | NM_014286 | | | | | GAAGACCTACTTTACCGAGAA | TTCTCGGTAAAGTAGGTCTTC | | 11592 | 6 |
| NME1_2 | TRCN0000010062 | CDS | NME1 | 4830 | Human | NM_000269 | Yes | MCF7 | TaqMan | 0.91 | CCGCCTTGTTGGTCTGAAATT | AATTTCAGACCAACAAGGCGG | | 9163 | 7 |
| NOA1_1 | TRCN0000149579 | CDS | C4ORF14 | 84273 | Human | NM_032313 | Yes | MCF7 | SYBR | 0.99 | GCCAGGTACTACATTAAACCT | AGGTTTAATGTAGTACCTGGC | | 14674 | 5 |
| PI4KB_1 | TRCN0000005695 | CDS | PI4KB | 5298 | Human | NM_002651 | Yes | A549 | SYBR | 0.94 | CGACATGTTCAACTACTATAA | TTATAGTAGTTGAACATGTCG | | 14118 | 5 |
| PICK1_1 | TRCN0000037907 | CDS | PICK1 | 9463 | Human | NM_012407 | Yes | MCF7 | SYBR | 0.8 | CTGAACACGTACCTCAACAAA | TTTGTTGAGGTACGTGTTCAG | | 13242 | 7 |
| PIP5K1A_1 | TRCN0000024518 | CDS | PIP5K1A | 18720 | Mouse | NM_008847 | Yes | Hepa 1-6 | SYBR | 0.72 | GCTTCCAGGATACTACATGAA | TTCATGTAGTATCCTGGAAGC | Y | 10957 | 4 |
| PIP5K1B_20 | TRCN0000230744 | CDS | PIP5K1B | 8395 | Human | NM_003558 | Yes | MCF7 | SYBR | 0.79 | ACGACAGGCCTACACTCTATT | AATAGAGTGTAGGCCTGTCGT | | 14613 | 4 |
| PIP5K1C_2 | TRCN0000378595 | CDS | PIP5K1C | 23396 | Human | NM_012398 | Yes | MCF7 | SYBR | 0.79 | CGGCGAGAGCGACACATAATT | AATTATGTGTCGCTCTCGCCG | | 10617 | 4 |
| PLD1_1 | TRCN0000076820 | CDS | PLD1 | 18805 | Mouse | NM_008875 | Yes | Hepa 1-6 | SYBR | 0.9 | CCCAATGATGAAGTACACAAT | ATTGTGTACTTCATCATTGGG | Y | 10422 | 2 |
| PLD2_1 | TRCN0000051149 | CDS | PLD2 | 5338 | Human | NM_002663 | Yes | A3 | SYBR | 0.58 | GCCAAGTACAAGACTCCCATA | TATGGGAGTCTTGTACTTGGC | | 12996 | 4 |
| PLD3_1 | TRCN0000052165 | CDS | PLD3 | 23646 | Human | NM_012268 | Yes | A549 | SYBR | 0.98 | CTACTCAACGTGGTGGACAAT | ATTGTCCACCACGTTGAGTAG | | 9388 | 4 |
| PLEKHA8_1 | TRCN0000146594 | 3UTR | PLEKHA8 | 84725 | Human | NM_032639 | | | | | CCAAGGTCAAACTGGAAATTA | TAATTTCCAGTTTGACCTTGG | | 10440 | 4 |
| PLEKHA8_2 | TRCN0000147794 | 3UTR | PLEKHA8 | 84725 | Human | NM_032639 | | | | | GCCTTCTTAAGGAAACCATTT | AAATGGTTTCCTTAAGAAGGC | | 11668 | 4 |
| PLEKHA8_3 | TRCN0000146652 | CDS | PLEKHA8 | 84725 | Human | NM_032639 | | | | | CAAATGGCAGTCTGTGAAATT | AATTTCACAGACTGCCATTTG | | 8116 | 4 |
| PLEKHA8_4 | TRCN0000146717 | CDS | PLEKHA8 | 84725 | Human | NM_032639 | | | | | CCTGTTAAGATGGATCTTGTT | AACAAGATCCATCTTAACAGG | | 14765 | 7 |
| PLEKHA8_5 | TRCN0000147293 | CDS | PLEKHA8 | 84725 | Human | NM_032639 | | | | | GTGGTTCCAGTATTAGACAAA | TTTGTCTAATACTGGAACCAC | | 11872 | 4 |
| PLIN2_1 | TRCN0000134012 | CDS | PLIN2 | 123 | Human | NM_001122 | Yes | A549 | SYBR | 0.77 | CTTTAGATGACGTGATGGATT | AATCCATCACGTCATCTAAAG | | 12644 | 5 |
| PREB_1 | TRCN0000107832 | CDS | PREB | 10113 | Human | NM_013388 | Yes | A549 | SYBR | 0.86 | GCTAGAACTCAGGGTAGAGAA | TTCTCTACCCTGAGTTCTAGC | | 8492 | 4 |
| PSD_1 | TRCN0000234810 | CDS | PSD | 5662 | Human | NM_002779 | Yes | PC3 | WB | | GGGAGTACCTCAAGTTCTTTG | CAAAGAACTTGAGGTACTCCC | | 10202 | 1 |
| PSD_2 | TRCN0000234811 | CDS | PSD | 5662 | Human | NM_002779 | Yes | PC3 | WB | | CCTGGATCACTCGCATCAATG | CATTGATGCGAGTGATCCAGG | | 9007 | 1 |
| PSD_3 | TRCN0000234812 | 3UTR | PSD | 5662 | Human | NM_002779 | Yes | PC3 | WB | | GCTTGGACAGAGACCAGGATT | AATCCTGGTCTCTGTCCAAGC | | 8868 | 1 |
| PSD_4 | TRCN0000238782 | CDS | PSD | 5662 | Human | NM_002779 | Yes | PC3 | WB | | GGTTGGAGCAGATGGCCTTTA | TAAAGGCCATCTGCTCCAACC | | 13302 | 6 |
| PSD_5 | TRCN0000238783 | CDS | PSD | 5662 | Human | NM_002779 | Yes | PC3 | WB | | GAGAATCGCTAGAGCCAAATG | CATTTGGCTCTAGCGATTCTC | | 10657 | 1 |
| PSD2_1 | TRCN0000160504 | 3UTR | PSD2 | 84249 | Human | NM_032289 | | | | | CCTGTTTATATTTGGGTCTTT | AAAGACCCAAATATAAACAGG | | 7742 | 1 |
| PSD2_2 | TRCN0000160769 | CDS | PSD2 | 84249 | Human | NM_032289 | | | | | CCCAATGGATTCCATGAAGAT | ATCTTCATGGAATCCATTGGG | | 10290 | 4 |
| PSD2_3 | TRCN0000161046 | CDS | PSD2 | 84249 | Human | NM_032289 | | | | | GACCCTTTACAACTCCATCAA | TTGATGGAGTTGTAAAGGGTC | | 10148 | 7 |
| PSD2_4 | TRCN0000162449 | 3UTR | PSD2 | 84249 | Human | NM_032289 | | | | | CCTGTGACTTATACTGCTCTT | AAGAGCAGTATAAGTCACAGG | | 12405 | 5 |
| PSD2_5 | TRCN0000165014 | CDS | PSD2 | 84249 | Human | NM_032289 | | | | | GCGTGGCTGGAAGAAATTCTA | TAGAATTTCTTCCAGCCACGC | | 9682 | 1 |
| PSD3_1 | TRCN0000123164 | 3UTR | PSD3 | 23362 | Human | NM_015310 | | | | | CGCCTTTATATGTGAAATCTT | AAGATTTCACATATAAAGGCG | | 7235 | 1 |

| | | | | | | | | | | | | | | | |
|---|---|---|---|---|---|---|---|---|---|---|---|---|---|---|---|
| PSD3_2 | TRCN0000123165 | CDS | PSD3 | 23362 | Human | NM_015310 | | | | | GCTCCCAGTTTGAAACCATTT | AAATGGTTTCAAACTGGGAGC | | 14042 | 4 |
| PSD3_3 | TRCN0000123166 | CDS | PSD3 | 23362 | Human | NM_015310 | | | | | CCGGAACATTACAGGCTACAA | TTGTAGCCTGTAATGTTCCGG | | 11182 | 5 |
| PSD3_4 | TRCN0000123167 | CDS | PSD3 | 23362 | Human | NM_015310 | | | | | CCTGTGCAATAATGCTTCTTA | TAAGAAGCATTATTGCACAGG | | 7349 | 5 |
| PSD3_5 | TRCN0000123168 | CDS | PSD3 | 23362 | Human | NM_015310 | | | | | CTGTGCAATAATGCTTCTTAA | TTAAGAAGCATTATTGCACAG | | 12401 | 7 |
| PSD4_1 | TRCN0000181027 | CDS | PSD4 | 23550 | Human | NM_012455 | Yes | PC3 | SYBR | | CCTCCTGAATCTACCAGACAA | TTGTCTGGTAGATTCAGGAGG | | 9112 | 6 |
| PSD4_2 | TRCN0000416818 | CDS | PSD4 | 23550 | Human | NM_012455 | Yes | PC3 | SYBR | | GCTGGAGACAAGCTAGCTAAT | ATTAGCTAGCTTGTCTCCAGC | | 10179 | 7 |
| PSD4_3 | TRCN0000424411 | CDS | PSD4 | 23550 | Human | NM_012455 | Yes | PC3 | SYBR | | TGCAGAAGAACAATGACTTTA | TAAAGTCATTGTTCTTCTGCA | | 10321 | 1 |
| PSD4_4 | TRCN0000427083 | 3UTR | PSD4 | 23550 | Human | NM_012455 | Yes | PC3 | SYBR | | TCCCTGGAGGAGACTTATTTC | GAAATAAGTCTCCTCCAGGGA | | 8404 | 2 |
| PSD4_5 | TRCN0000429097 | 3UTR | PSD4 | 23550 | Human | NM_012455 | Yes | PC3 | SYBR | | GAATCCAGAGTGGCCTCATTT | AAATGAGGCCACTCTGGATTC | | 12684 | 5 |
| RAB11FIP1_1 | TRCN0000145281 | CDS | RAB11FIP1 | 80223 | Human | NM_025151 | Yes | A549 | SYBR | 0.93 | GAAATCCAAACCAGGAAAGAA | TTCTTTCCTGGTTTGGATTTC | | 12773 | 5 |
| RAB11FIP2_1 | TRCN0000144978 | CDS | RAB11FIP2 | 22841 | Human | NM_014904 | Yes | A549 | SYBR | 0.78 | GAATTGTGTTTCGGAAGACAA | TTGTCTTCCGAAACACAATTC | | 12992 | 5 |
| RAB11FIP3_1 | TRCN0000187717 | CDS | RAB11FIP3 | 215445 | Mouse | XM_484616 | Yes | Hepa 1-6 | SYBR | 0.74 | GAAGAGCATTGAGATCGAGAA | TTCTCGATCTCAATGCTCTTC | Y | 9817 | 2 |
| RAB11FIP4_1 | TRCN0000413922 | 3UTR | RAB11FIP4 | 84440 | Human | NM_032932 | Yes | PC3 | WB | | TCTAGTACGATGGGCTCTTTC | GAAAGAGCCCATCGTACTAGA | | 11318 | 5 |
| RAB11FIP4_2 | TRCN0000056503 | CDS | RAB11FIP4 | 84440 | Human | NM_032932 | Yes | PC3 | WB | | CGACAATGACATCACAGAGAA | TTCTCTGTGATGTCATTGTCG | | 8569 | 5 |
| RAB11FIP4_3 | TRCN0000056504 | CDS | RAB11FIP4 | 84440 | Human | NM_032932 | | | | | GCACGTGTACAACAGCGAATT | AATTCGCTGTTGTACACGTGC | | 9689 | 4 |
| RAB11FIP4_4 | TRCN0000056505 | CDS | RAB11FIP4 | 84440 | Human | NM_032932 | | | | | GAGGCAGTACATGGACAAGAT | ATCTTGTCCATGTACTGCCTC | | 9267 | 2 |
| RAB11FIP4_5 | TRCN0000056506 | CDS | RAB11FIP4 | 84440 | Human | NM_032932 | Yes | PC3 | WB | | AGAATCAACTTCAAGGACTTT | AAAGTCCTTGAAGTTGATTCT | | 9554 | 5 |
| RAB11FIP5_1 | TRCN0000007155 | CDS | RAB11FIP5 | 26056 | Human | NM_015470 | Yes | A549 | SYBR | 0.76 | GCAAGCTGTCTCTTCCCGGTT | AACCGGGAAGAGACAGCTTGC | | 9031 | 6 |
| RALA_1 | TRCN0000315072 | CDS | RALA | 5898 | Human | NM_005402 | Yes | A549 | SYBR | 0.95 | GGAGGAAGTCCAGATCGATAT | ATATCGATCTGGACTTCCTCC | | 12250 | 7 |
| RALB_1 | TRCN0000072956 | CDS | RALB | 5899 | Human | NM_002881 | Yes | A549 | SYBR | 0.97 | CAAGGTGTTCTTTGACCTAAT | ATTAGGTCAAAGAACACCTTG | | 11671 | 4 |
| RCHY1_1 | TRCN0000010849 | CDS | RCHY1 | 25898 | Human | NM_015436 | Yes | 293T/17 | SYBR | 0.8 | AGTGAAGGAAGTGCAGTGCAT | ATGCACTGCACTTCCTTCACT | | 9973 | 6 |
| SMAP1_6 | TRCN0000151035 | 3UTR | SMAP1 | 60682 | Human | NM_021940 | Yes | A549 | SYBR | 0.88 | GCAGCACAAGTGTAATGAATA | TATTCATTACACTTGTGCTGC | | 11771 | 7 |
| SMAP2_1 | TRCN0000146695 | CDS | SMAP2 | 64744 | Human | NM_022733 | Yes | MCH58 | SYBR | 0.89 | CGACTTTATGAAGCCTATCTT | AAGATAGGCTTCATAAAGTCG | | 11150 | 6 |
| STAT1_1 | TRCN0000280021 | CDS | STAT1 | 6772 | Human | NM_007315 | Yes | A549 | SYBR | 0.93 | CTGGAAGATTTACAAGATGAA | TTCATCTTGTAAATCTTCCAG | | 11389 | 4 |
| STAT3_1 | TRCN0000329887 | CDS | STAT3 | 6774 | Human | NM_003150 | Yes | A549 | SYBR | 0.88 | GCACAATCTACGAAGAATCAA | TTGATTCTTCGTAGATTGTGC | | 12790 | 6 |
| STAT5A_1 | TRCN0000232134 | CDS | STAT5A | 6776 | Human | NM_003152 | Yes | MCH58 | SYBR | 0.95 | GGACCTTCTTGTTGCGCTTTA | TAAAGCGCAACAAGAAGGTCC | | 8093 | 4 |
| STAT5B_1 | TRCN0000232140 | CDS | STAT5B | 6777 | Human | NM_012448 | Yes | A549 | SYBR | 0.98 | TATGTCCCTGAAACGAATTAA | TTAATTCGTTTCAGGGACATA | | 13717 | 4 |
| TBC1D10A_1 | TRCN0000160822 | CDS | TBC1D10A | 83874 | Human | NM_031937 | Yes | MCH58 | SYBR | 0.81 | CAAGGTGAAGTTACAGCAGAA | TTCTGCTGTAACTTCACCTTG | | 12506 | 4 |
| TBC1D10B_1 | TRCN0000106258 | CDS | TBC1D10B | 68449 | Mouse | NM_144522 | Yes | Hepa 1-6 | SYBR | 0.92 | GCAGTACCTGTCTAATAGCAA | TTGCTATTAGACAGGTACTGC | Y | 10236 | 4 |
| TBC1D10C_1 | TRCN0000242973 | 3UTR | TBC1D10C | 374403 | Human | NM_198517 | | | | | GGCAAATAGGCACCGCACTTT | AAAGTGCGGTGCCTATTTGCC | | 11190 | 7 |
| TBC1D10C_2 | TRCN0000242972 | CDS | TBC1D10C | 374403 | Human | NM_198517 | | | | | CGGCGGTACAAGAAGGTAAAG | CTTTACCTTCTTGTACCGCCG | | 10964 | 4 |
| TBC1D10C_3 | TRCN0000242974 | CDS | TBC1D10C | 374403 | Human | NM_198517 | | | | | TCAAGGCCTACACCCTGTATC | GATACAGGGTGTAGGCCTTGA | | 10138 | 5 |
| TBC1D10C_4 | TRCN0000242975 | CDS | TBC1D10C | 374403 | Human | NM_198517 | | | | | CTGAGAGGACCATGGACTTAG | CTAAGTCCATGGTCCTCTCAG | | 8928 | 5 |
| TBC1D10C_5 | TRCN0000257000 | CDS | TBC1D10C | 374403 | Human | NM_198517 | | | | | CCGACCGCTATGGATTCATTG | CAATGAATCCATAGCGGTCGG | | 11789 | 4 |
| TBC1D24_1 | TRCN0000245854 | CDS | TBC1D24 | 57465 | Human | NM_020705 | Yes | A549 | SYBR | 0.76 | GGAGTGAGAGAAATAAGTTTG | CAAACTTATTTCTCTCACTCC | | 10009 | 4 |
| TBC1D3_1 | TRCN0000255682 | 3UTR | TBC1D3 | 729873 | Human | NM_001123391 | Yes | PC3 | WB | | GGATAATTTCCCTAGGCTTAA | TTAAGCCTAGGGAAATTATCC | | 9424 | 4 |
| TBC1D3_2 | TRCN0000255679 | CDS | TBC1D3 | 729873 | Human | NM_001123391 | | | | | TAGGCTGCCTCATCCGGATAT | ATATCCGGATGAGGCAGCCTA | | 14590 | 7 |
| TBC1D3_3 | TRCN0000255680 | CDS | TBC1D3 | 729873 | Human | NM_001123391 | Yes | PC3 | WB | | TTGCAACCGGTTCGTTGATAC | GTATCAACGAACCGGTTGCAA | | 16228 | 7 |
| TBC1D3_4 | TRCN0000255681 | CDS | TBC1D3 | 729873 | Human | NM_001123391 | Yes | PC3 | WB | | GGACATTAAGGAAGCATATAT | ATATATGCTTCCTTAATGTCC | | 8551 | 2 |
| TBC1D3_5 | TRCN0000265682 | CDS | TBC1D3 | 729873 | Human | NM_001123391 | | | | | GATAACAAGAATCGCCTTTAA | TTAAAGGCGATTCTTGTTATC | | 12497 | 5 |
| TMED10_1 | TRCN0000029148 | CDS | TMED10 | 10972 | Human | NM_006827 | Yes | A549 | SYBR | 0.99 | CGCTTCTTCAAGGCCAAGAAA | TTTCTTGGCCTTGAAGAAGCG | | 16825 | 4 |
| TMED2_1 | TRCN0000065305 | CDS | TMED2 | 10959 | Human | NM_006815 | Yes | A549 | SYBR | 0.99 | CTCGGGCTATTTCGTTAGCAT | ATGCTAACGAAATAGCCCGAG | | 14240 | 4 |
| UACA_1 | TRCN0000159821 | CDS | UACA | 55075 | Human | NM_018003 | Yes | MCH58 | SYBR | 0.86 | GCAGAGCATTTGCAGATTAAA | TTTAATCTGCAAATGCTCTGC | | 10842 | 4 |
| USP6_1 | TRCN0000218643 | CDS | USP6 | 9098 | Human | NM_004505 | Yes | MCF7 | SYBR | 0.83 | GCTTCTAGTCCAACACAAATA | TATTTGTGTTGGACTAGAAGC | | 9668 | 4 |
| USP9X_1 | TRCN0000007361 | 3UTR | USP9X | 8239 | Human | NM_004652 | Yes | A549 | SYBR | 0.93 | GAGAGTTTATTCACTGTCTTA | TAAGACAGTGAATAAACTCTC | | 10797 | 5 |
| VEZT_1 | TRCN0000236554 | CDS | VEZT | 55591 | Human | NM_017599 | Yes | MCH58 | SYBR | 0.87 | TCAGACGGTTAGCCCTATTAC | GTAATAGGGCTAACCGTCTGA | | 6873 | 2 |
