## Supplementary material for "The small GTPase ARF3 controls metastasis and invasion modality by regulating N-cadherin levels": Supplemenary Table S2

| shRNA Clone ID | TRC ID | Gene Symbol | Target Sequence (sense) |
|---|---|---|---|
| Scr | N/A | N/A | CCGCAGGTATGCACGCGT |
| ARF1 KD2 | TRCN0000039875 | ARF1 | CCTCCTGGTGTTCGCCAACAA |
| ARF1 KD4 | TRCN0000039873 | ARF1 | CCTAGCATAGATTTGCAGTTA |
| ARF3 KD1 | TRCN0000047674 | ARF3 | GATGCTGTACTCCTTGTCTTT |
| ARF3 KD2 | TRCN0000047676 | ARF3 | GAACACCCAAGGGTTGATATT |
| N-Cadherin KD1 | TRCN0000327706 | CDH2 | CGCATTATGCAAGACTGGATT |
| N-Cadherin KD2 | TRCN0000327707 | CDH2 | CCAGTGACTATTAAGAGAAAT |
| PSD KD1 | TRCN0000234810 | PSD | GGGAGTACCTCAAGTTCTTTG |
| PSD KD2 | TRCN0000234811 | PSD | CCTGGATCACTCGCATCAATG |
| RAB11FIP4 KD1 | TRCN0000413922 | RAB11FIP4 | TCTAGTACGATGGGCTCTTTC |
| RAB11FIP4 KD2 | TRCN0000056503 | RAB11FIP4 | CGACAATGACATCACAGAGAA |

| Target Sequence (anti-sense) |
| --- |
| ACGCGTGCATACCTGCGG |
| TTGTTGGCGAACACCAGGAGG |
| TAACTGCAAATCTATGCTAGG |
| AAAGACAAGGAGTACAGCATC |
| AATACAACCCTTGGGTGTTC |
| AATCCAGTCTTGCATAATGCG |
| ATTTCTCTTAATAGTCACTGG |
| CAAAGAACTTGAGGTACTCCC |
| CATTGATGCGAGTGATCCAGG |
| GAAAGAGCCCATCGTACTAGA |
| TTCTCTGTGATGTCATTGTCG |
