## Supplementary material for "The small GTPase ARF3 controls metastasis and invasion modality by regulating N-cadherin levels": Supplemenary Table S3

| Antibody | Company | Cat. No. | Application | Dilution |
|---|---|---|---|---|
| Alexa Fluor 488 anti-mouse IgG | Thermo Fisher Scientific | A21202 | IF | 1 in 200 |
| Alexa Fluor 568 Phalloidin | Thermo Fisher Scientific | A12380 | IF | 1 in 200 |
| Alexa Fluor 594 anti-rabbit IgG | Thermo Fisher Scientific | A11012 | IF | 1 in 200 |
| anti-ARF1 | Novus Biologicals | NB-110-85530 | WB | 1 in 1000 |
| anti-ARF3 | BD Biosciences | 610784 | IF | 1 in 100 |
| anti-ARF3 | BD Biosciences | 610784 | WB | 1 in 1000 |
| anti-ARF3 | BD Biosciences | 610784 | IP | 2ug |
| anti-ARF4 | Proteintech | 11673-1-AP | WB | 1 in 1000 |
| anti-ARF5 | Novus Biologicals | H00000381-M01 | WB | 1 in 1000 |
| anti-ARF6 | Sigma Aldrich | A5230 | WB | 1 in 1000 |
| anti-E-Cadherin | Cell Signalling Technology | 3195 | WB | 1 in 1000 |
| anti-E-Cadherin | BD Biosciences | 610181 | WB | 1 in 1000 |
| anti-GAPDH | Cell Signalling Technology | 2118 | WB | 1 in 5000 |
| anti-GST | Sigma Aldrich | 06-332 | WB | 1 in 1000 |
| HCS CellMask Deep Red Stain | Thermo Fisher Scientific | H32721 | IF | 1 in 10,000 |
| Hoechst 34580 | Thermo Fisher Scientific | H21486 | IF | 1 in 1000 |
| anti-mNeonGreen | Chromotek | 32F6 | WB | 1 in 1000 |
| anti-N-Cadherin | Cell Signalling Technology | 13116 | IF | 1 in 200 |
| anti-N-Cadherin | Cell Signalling Technology | 13116 | WB | 1 in 1000 |
| anti-PSD | Novus Biologicals | NBP2-15083 | WB | 1 in 1000 |
| anti-Rab11FIP4 | Abcam | ab74446 | WB | 1 in 1000 |
