## Supplementary material for "The small GTPase ARF3 controls metastasis and invasion modality by regulating N-cadherin levels": Supplemenary Table S4

| TumourType | Dataset_ID | Normal vs Tumour | Count (Sample_ID) |
|---|---|---|---|
| Adrenal_Gland | GSE68606 | Cancer | 6 |
| Adrenal_Gland | GSE75415 | Cancer | 23 |
| Adrenal_Gland | GSE75415 | Normal | 7 |
| Adrenal_Gland | GSE76021 | Cancer | 29 |
| Bladder | GSE12630 | Cancer | 11 |
| Bladder | GSE2361 | Normal | 1 |
| Bladder | GSE3167 | Cancer | 5 |
| Bladder | GSE3167 | Normal | 5 |
| Blood | E-AFMX-5 | Normal | 24 |
| Blood | E-MEXP-120 | Cancer | 31 |
| Blood | E-MEXP-313 | Cancer | 99 |
| Blood | E-TABM-117 | Cancer | 37 |
| Blood | E-TABM-125 | Cancer | 68 |
| Blood | GSE1010 | Normal | 12 |
| Blood | GSE10255 | Cancer | 161 |
| Blood | GSE10631 | Cancer | 30 |
| Blood | GSE10631 | Normal | 14 |
| Blood | GSE11038 | Cancer | 24 |
| Blood | GSE1124 | Normal | 5 |
| Blood | GSE1133 | Cancer | 12 |
| Blood | GSE1133 | Normal | 24 |
| Blood | GSE1140 | Normal | 15 |
| Blood | GSE11582 | Normal | 374 |
| Blood | GSE11907 | Normal | 10 |
| Blood | GSE12845 | Normal | 14 |
| Blood | GSE12995 | Cancer | 175 |
| Blood | GSE13280 | Cancer | 44 |
| Blood | GSE13591 | Cancer | 153 |
| Blood | GSE13591 | Normal | 4 |
| Blood | GSE13996 | Cancer | 67 |
| Blood | GSE1427 | Cancer | 99 |
| Blood | GSE14286 | Cancer | 13 |
| Blood | GSE14317 | Cancer | 19 |
| Blood | GSE14317 | Normal | 7 |
| Blood | GSE14577 | Normal | 7 |
| Blood | GSE1466 | Cancer | 41 |
| Blood | GSE1466 | Normal | 6 |
| Blood | GSE15777 | Cancer | 22 |
| Blood | GSE15777 | Normal | 3 |
| Blood | GSE1751 | Normal | 14 |
| Blood | GSE2113 | Cancer | 52 |
| Blood | GSE2351 | Cancer | 2 |
| Blood | GSE2779 | Normal | 9 |
| Blood | GSE3365 | Normal | 42 |
| Blood | GSE3912 | Cancer | 35 |

| | | | |
|---|---|---|---|
| Blood | GSE4119 | Cancer | 72 |
| Blood | GSE4475 | Cancer | 135 |
| Blood | GSE4698 | Cancer | 60 |
| Blood | GSE5122 | Cancer | 58 |
| Blood | GSE5580 | Normal | 21 |
| Blood | GSE5788 | Cancer | 6 |
| Blood | GSE5808 | Normal | 3 |
| Blood | GSE5820 | Cancer | 29 |
| Blood | GSE5967 | Normal | 7 |
| Blood | GSE6236 | Normal | 28 |
| Blood | GSE6269 | Cancer | 4 |
| Blood | GSE6269 | Normal | 6 |
| Blood | GSE635 | Cancer | 121 |
| Blood | GSE6365 | Cancer | 90 |
| Blood | GSE6401 | Cancer | 102 |
| Blood | GSE6477 | Cancer | 147 |
| Blood | GSE6477 | Normal | 15 |
| Blood | GSE6613 | Normal | 33 |
| Blood | GSE6691 | Cancer | 43 |
| Blood | GSE6740 | Normal | 10 |
| Blood | GSE7148 | Normal | 14 |
| Blood | GSE7429 | Normal | 20 |
| Blood | GSE7638 | Normal | 50 |
| Blood | GSE7893 | Normal | 11 |
| Blood | GSE8650 | Normal | 21 |
| Blood | GSE8970 | Cancer | 5 |
| Blood | GSE9006 | Normal | 24 |
| Blood | GSE9476 | Cancer | 26 |
| Blood | GSE9476 | Normal | 38 |
| Blood | GSE9874 | Normal | 30 |
| Bone_Marrow | GSE10139 | Cancer | 40 |
| Bone_Marrow | GSE10172 | Cancer | 36 |
| Bone_Marrow | GSE11038 | Cancer | 36 |
| Bone_Marrow | GSE1159 | Cancer | 293 |
| Bone_Marrow | GSE12417 | Cancer | 163 |
| Bone_Marrow | GSE13425 | Cancer | 131 |
| Bone_Marrow | GSE14618 | Cancer | 42 |
| Bone_Marrow | GSE15347 | Cancer | 89 |
| Bone_Marrow | GSE1577 | Cancer | 29 |
| Bone_Marrow | GSE16102 | Normal | 2 |
| Bone_Marrow | GSE16131 | Cancer | 17 |
| Bone_Marrow | GSE16334 | Normal | 11 |
| Bone_Marrow | GSE16746 | Cancer | 37 |
| Bone_Marrow | GSE17195 | Cancer | 44 |
| Bone_Marrow | GSE1729 | Cancer | 43 |
| Bone_Marrow | GSE18026 | Normal | 2 |

| | | | |
|---|---|---|---|
| Bone_Marrow | GSE19147 | Cancer | 25 |
| Bone_Marrow | GSE19147 | Normal | 8 |
| Bone_Marrow | GSE22529 | Cancer | 41 |
| Bone_Marrow | GSE22529 | Normal | 11 |
| Bone_Marrow | GSE2361 | Normal | 1 |
| Bone_Marrow | GSE28497 | Cancer | 284 |
| Bone_Marrow | GSE28497 | Normal | 4 |
| Bone_Marrow | GSE34171 | Cancer | 78 |
| Bone_Marrow | GSE34860 | Cancer | 13 |
| Bone_Marrow | GSE37088 | Cancer | 72 |
| Bone_Marrow | GSE37642 | Cancer | 291 |
| Bone_Marrow | GSE43176 | Normal | 4 |
| Bone_Marrow | GSE44164 | Cancer | 32 |
| Bone_Marrow | GSE4475 | Cancer | 86 |
| Bone_Marrow | GSE48184 | Cancer | 29 |
| Bone_Marrow | GSE51082 | Cancer | 139 |
| Bone_Marrow | GSE5788 | Normal | 8 |
| Bone_Marrow | GSE647 | Cancer | 8 |
| Bone_Marrow | GSE649 | Cancer | 31 |
| Bone_Marrow | GSE660 | Cancer | 13 |
| Bone_Marrow | GSE67684 | Cancer | 39 |
| Bone_Marrow | GSE68954 | Cancer | 9 |
| Bone_Marrow | GSE68954 | Normal | 6 |
| Bone_Marrow | GSE78132 | Cancer | 44 |
| Bone_Marrow | GSE83449 | Cancer | 15 |
| Bone_Marrow | GSE8510 | Cancer | 5 |
| Bone_Marrow | GSE8835 | Cancer | 42 |
| Bone_Marrow | GSE8835 | Normal | 24 |
| Bone_Marrow | GSE8879 | Cancer | 55 |
| Bone_Marrow | GSE8970 | Cancer | 29 |
| Bone_Marrow | GSE9429 | Cancer | 10 |
| Brain | E-AFMX-5 | Normal | 45 |
| Brain | E-MEXP-114 | Normal | 23 |
| Brain | E-MEXP-1690 | Cancer | 6 |
| Brain | E-MEXP-1690 | Normal | 6 |
| Brain | GSE1133 | Normal | 52 |
| Brain | GSE1147 | Normal | 23 |
| Brain | GSE12649 | Normal | 34 |
| Brain | GSE12685 | Normal | 8 |
| Brain | GSE12907 | Cancer | 21 |
| Brain | GSE12907 | Normal | 4 |
| Brain | GSE1297 | Cancer | 31 |
| Brain | GSE13041 | Cancer | 191 |
| Brain | GSE13471 | Normal | 5 |
| Brain | GSE1993 | Cancer | 65 |
| Brain | GSE20186 | Cancer | 18 |

| | | | |
|---|---|---|---|
| Brain | GSE20186 | Normal | 18 |
| Brain | GSE20295 | Cancer | 40 |
| Brain | GSE20295 | Normal | 53 |
| Brain | GSE2175 | Cancer | 3 |
| Brain | GSE2175 | Normal | 1 |
| Brain | GSE2361 | Normal | 2 |
| Brain | GSE24072 | Cancer | 32 |
| Brain | GSE24250 | Cancer | 8 |
| Brain | GSE24250 | Normal | 6 |
| Brain | GSE2485 | Cancer | 9 |
| Brain | GSE2719 | Normal | 1 |
| Brain | GSE2841 | Cancer | 76 |
| Brain | GSE3185 | Cancer | 3 |
| Brain | GSE3790 | Normal | 87 |
| Brain | GSE4271 | Cancer | 100 |
| Brain | GSE4412 | Cancer | 85 |
| Brain | GSE4780 | Cancer | 5 |
| Brain | GSE5390 | Normal | 8 |
| Brain | GSE5392 | Normal | 42 |
| Brain | GSE62600 | Cancer | 19 |
| Brain | GSE62600 | Normal | 3 |
| Brain | GSE6306 | Normal | 1218 |
| Brain | GSE6613 | Cancer | 50 |
| Brain | GSE6613 | Normal | 22 |
| Brain | GSE83294 | Cancer | 85 |
| Brain | GSE8397 | Cancer | 47 |
| Brain | GSE84422 | Cancer | 951 |
| Brain | GSE8692 | Cancer | 12 |
| Brain | GSE9335 | Normal | 17 |
| Brain | GSE9963 | Normal | 18 |
| Breast | GSE11121 | Cancer | 200 |
| Breast | GSE11965 | Cancer | 16 |
| Breast | GSE12093 | Cancer | 136 |
| Breast | GSE12237 | Cancer | 6 |
| Breast | GSE12630 | Cancer | 11 |
| Breast | GSE1456 | Cancer | 159 |
| Breast | GSE1561 | Cancer | 49 |
| Breast | GSE15852 | Cancer | 43 |
| Breast | GSE15852 | Normal | 43 |
| Breast | GSE16873 | Cancer | 12 |
| Breast | GSE16873 | Normal | 12 |
| Breast | GSE2034 | Cancer | 286 |
| Breast | GSE22093 | Cancer | 103 |
| Breast | GSE2361 | Normal | 1 |
| Breast | GSE23988 | Cancer | 61 |
| Breast | GSE24185 | Cancer | 103 |

| | | | |
|---|---|---|---|
| Breast | GSE24509 | Cancer | 2 |
| Breast | GSE25066 | Cancer | 508 |
| Breast | GSE2603 | Cancer | 99 |
| Breast | GSE31519 | Cancer | 9 |
| Breast | GSE32072 | Cancer | 8 |
| Breast | GSE3494 | Cancer | 251 |
| Breast | GSE36774 | Cancer | 149 |
| Breast | GSE3726 | Cancer | 62 |
| Breast | GSE45255 | Cancer | 10 |
| Breast | GSE4611 | Cancer | 216 |
| Breast | GSE46184 | Cancer | 74 |
| Breast | GSE48984 | Normal | 3 |
| Breast | GSE4922 | Cancer | 289 |
| Breast | GSE5327 | Cancer | 58 |
| Breast | GSE5364 | Cancer | 183 |
| Breast | GSE5364 | Normal | 13 |
| Breast | GSE5462 | Cancer | 116 |
| Breast | GSE5847 | Cancer | 95 |
| Breast | GSE6532 | Cancer | 327 |
| Breast | GSE6772 | Cancer | 24 |
| Breast | GSE6772 | Normal | 2 |
| Breast | GSE6883 | Cancer | 9 |
| Breast | GSE6883 | Normal | 3 |
| Breast | GSE68892 | Cancer | 99 |
| Breast | GSE7390 | Cancer | 198 |
| Breast | GSE83232 | Cancer | 258 |
| Breast | GSE92697 | Cancer | 26 |
| Breast | GSE9574 | Cancer | 14 |
| Breast | GSE9574 | Normal | 15 |
| Breast | GSE9662 | Cancer | 24 |
| Cartilage | GSE15757 | Cancer | 4 |
| Cartilage | GSE37372 | Cancer | 20 |
| Cartilage | GSE6565 | Normal | 2 |
| Cartilage | GSE7007 | Cancer | 27 |
| Cervix | GSE7803 | Cancer | 31 |
| Cervix | GSE7803 | Normal | 10 |
| Colon | E-MEXP-383 | Cancer | 36 |
| Colon | E-MEXP-833 | Cancer | 24 |
| Colon | E-MTAB-57 | Cancer | 18 |
| Colon | E-MTAB-57 | Normal | 13 |
| Colon | GSE1133 | Cancer | 2 |
| Colon | GSE1152 | Normal | 2 |
| Colon | GSE12630 | Cancer | 9 |
| Colon | GSE12945 | Cancer | 62 |
| Colon | GSE2138 | Cancer | 20 |
| Colon | GSE2361 | Normal | 1 |

| | | | |
|---|---|---|---|
| Colon | GSE24514 | Cancer | 34 |
| Colon | GSE24514 | Normal | 15 |
| Colon | GSE26682 | Cancer | 155 |
| Colon | GSE2719 | Normal | 1 |
| Colon | GSE3726 | Cancer | 42 |
| Colon | GSE4045 | Cancer | 37 |
| Colon | GSE41258 | Cancer | 186 |
| Colon | GSE41258 | Normal | 54 |
| Colon | GSE5364 | Cancer | 9 |
| Colon | GSE5364 | Normal | 9 |
| Colon | GSE62322 | Cancer | 20 |
| Colon | GSE62322 | Normal | 18 |
| Colon | GSE6272 | Normal | 1 |
| Colon | GSE68468 | Cancer | 365 |
| Colon | GSE7208 | Cancer | 59 |
| Colon | GSE77955 | Cancer | 34 |
| Colon | GSE77955 | Normal | 13 |
| Esophagus | GSE13083 | Normal | 7 |
| Esophagus | GSE1420 | Cancer | 8 |
| Esophagus | GSE1420 | Normal | 8 |
| Esophagus | GSE23400 | Cancer | 53 |
| Esophagus | GSE23400 | Normal | 53 |
| Esophagus | GSE37203 | Cancer | 15 |
| Esophagus | GSE5364 | Cancer | 16 |
| Esophagus | GSE5364 | Normal | 13 |
| Heart | E-AFMX-5 | Normal | 4 |
| Heart | GSE1133 | Normal | 4 |
| Heart | GSE12288 | Cancer | 222 |
| Heart | GSE1789 | Normal | 5 |
| Heart | GSE2240 | Normal | 5 |
| Heart | GSE2361 | Normal | 1 |
| Heart | GSE2719 | Normal | 1 |
| Heart | GSE3585 | Normal | 5 |
| Heart | GSE974 | Normal | 16 |
| Immune_System | GSE10325 | Cancer | 38 |
| Immune_System | GSE10325 | Normal | 28 |
| Immune_System | GSE11907 | Cancer | 108 |
| Immune_System | GSE38351 | Cancer | 14 |
| Immune_System | GSE46923 | Cancer | 5 |
| Immune_System | GSE46923 | Normal | 21 |
| Joint | GSE12021 | Cancer | 22 |
| Joint | GSE12021 | Normal | 9 |
| Joint | GSE38351 | Cancer | 8 |
| Joint | GSE55235 | Cancer | 20 |
| Joint | GSE55235 | Normal | 10 |
| Joint | GSE55457 | Cancer | 23 |

| | | | |
|---|---|---|---|
| Joint | GSE55457 | Normal | 10 |
| Joint | GSE55584 | Cancer | 16 |
| Joint | GSE63359 | Cancer | 46 |
| Joint | GSE63359 | Normal | 26 |
| Kidney | E-AFMX-5 | Normal | 2 |
| Kidney | E-TABM-53 | Cancer | 64 |
| Kidney | E-TABM-53 | Normal | 9 |
| Kidney | GSE10320 | Cancer | 144 |
| Kidney | GSE1133 | Normal | 2 |
| Kidney | GSE11482 | Cancer | 41 |
| Kidney | GSE11904 | Cancer | 21 |
| Kidney | GSE12630 | Cancer | 11 |
| Kidney | GSE14767 | Cancer | 39 |
| Kidney | GSE15641 | Cancer | 69 |
| Kidney | GSE15641 | Normal | 23 |
| Kidney | GSE16102 | Normal | 2 |
| Kidney | GSE2004 | Normal | 19 |
| Kidney | GSE2361 | Normal | 1 |
| Kidney | GSE2712 | Cancer | 35 |
| Kidney | GSE2719 | Normal | 1 |
| Kidney | GSE27556 | Normal | 1 |
| Kidney | GSE30946 | Cancer | 41 |
| Kidney | GSE31403 | Cancer | 224 |
| Kidney | GSE3297 | Normal | 6 |
| Kidney | GSE6280 | Cancer | 14 |
| Kidney | GSE6280 | Normal | 6 |
| Kidney | GSE6344 | Cancer | 10 |
| Kidney | GSE6344 | Normal | 10 |
| Kidney | GSE65162 | Normal | 3 |
| Kidney | GSE68606 | Cancer | 2 |
| Kidney | GSE781 | Cancer | 9 |
| Kidney | GSE781 | Normal | 8 |
| Larynx | GSE10935 | Cancer | 2 |
| Larynx | GSE10935 | Normal | 2 |
| Larynx | GSE27020 | Cancer | 109 |
| Liver | E-AFMX-5 | Normal | 4 |
| Liver | E-TABM-292 | Cancer | 32 |
| Liver | E-TABM-36 | Cancer | 60 |
| Liver | GSE1133 | Normal | 4 |
| Liver | GSE12630 | Cancer | 8 |
| Liver | GSE14323 | Cancer | 47 |
| Liver | GSE14323 | Normal | 19 |
| Liver | GSE16102 | Normal | 2 |
| Liver | GSE19281 | Normal | 3 |
| Liver | GSE2361 | Normal | 2 |
| Liver | GSE2719 | Normal | 1 |

| | | | |
|---|---|---|---|
| Liver | GSE41258 | Normal | 13 |
| Liver | GSE5364 | Cancer | 9 |
| Liver | GSE5364 | Normal | 8 |
| Liver | GSE60502 | Cancer | 18 |
| Liver | GSE60502 | Normal | 18 |
| Liver | GSE6272 | Normal | 4 |
| Liver | GSE7473 | Normal | 4 |
| Lung | E-AFMX-5 | Normal | 6 |
| Lung | E-MEXP-231 | Cancer | 49 |
| Lung | E-MEXP-231 | Normal | 9 |
| Lung | E-TABM-15 | Cancer | 23 |
| Lung | E-TABM-15 | Normal | 18 |
| Lung | GSE10072 | Cancer | 58 |
| Lung | GSE10072 | Normal | 49 |
| Lung | GSE1133 | Normal | 4 |
| Lung | GSE12630 | Cancer | 15 |
| Lung | GSE1650 | Normal | 12 |
| Lung | GSE17475 | Cancer | 28 |
| Lung | GSE19027 | Cancer | 21 |
| Lung | GSE19027 | Normal | 28 |
| Lung | GSE2361 | Normal | 2 |
| Lung | GSE2395 | Normal | 11 |
| Lung | GSE2549 | Cancer | 40 |
| Lung | GSE2549 | Normal | 9 |
| Lung | GSE2719 | Normal | 1 |
| Lung | GSE27556 | Normal | 1 |
| Lung | GSE31908 | Cancer | 30 |
| Lung | GSE31908 | Normal | 20 |
| Lung | GSE3593 | Cancer | 198 |
| Lung | GSE39262 | Normal | 1 |
| Lung | GSE40839 | Normal | 10 |
| Lung | GSE41258 | Normal | 7 |
| Lung | GSE4573 | Cancer | 130 |
| Lung | GSE5060 | Normal | 22 |
| Lung | GSE5364 | Cancer | 18 |
| Lung | GSE5364 | Normal | 12 |
| Lung | GSE6253 | Cancer | 18 |
| Lung | GSE68465 | Cancer | 321 |
| Lung | GSE68465 | Normal | 19 |
| Lung | GSE68606 | Cancer | 26 |
| Lung | GSE75324 | Cancer | 80 |
| Lung | GSE75324 | Normal | 52 |
| Lung | GSE7670 | Cancer | 35 |
| Lung | GSE7670 | Normal | 30 |
| Lung | GSE7895 | Normal | 104 |
| Lung | GSE994 | Normal | 72 |

| | | | |
|---|---|---|---|
| Lung | GSE9971 | Cancer | 27 |
| Muscle | E-AFMX-5 | Normal | 4 |
| Muscle | E-CBIL-30 | Normal | 17 |
| Muscle | GSE10760 | Normal | 59 |
| Muscle | GSE1133 | Normal | 4 |
| Muscle | GSE11681 | Normal | 10 |
| Muscle | GSE11971 | Normal | 4 |
| Muscle | GSE12648 | Normal | 10 |
| Muscle | GSE1462 | Normal | 3 |
| Muscle | GSE1551 | Normal | 10 |
| Muscle | GSE1786 | Normal | 12 |
| Muscle | GSE2361 | Normal | 1 |
| Muscle | GSE2719 | Normal | 1 |
| Muscle | GSE3112 | Normal | 11 |
| Muscle | GSE3307 | Normal | 12 |
| Muscle | GSE362 | Normal | 15 |
| Muscle | GSE4667 | Normal | 35 |
| Muscle | GSE474 | Normal | 8 |
| Muscle | GSE5370 | Cancer | 2 |
| Muscle | GSE6011 | Normal | 14 |
| Muscle | GSE674 | Normal | 15 |
| Muscle | GSE8441 | Normal | 22 |
| Muscle | GSE9397 | Normal | 2 |
| Muscle | GSE9676 | Normal | 60 |
| Ovary | E-AFMX-5 | Normal | 2 |
| Ovary | GSE1133 | Normal | 2 |
| Ovary | GSE12630 | Cancer | 9 |
| Ovary | GSE14764 | Cancer | 80 |
| Ovary | GSE23603 | Cancer | 28 |
| Ovary | GSE2361 | Normal | 1 |
| Ovary | GSE26712 | Cancer | 185 |
| Ovary | GSE26712 | Normal | 10 |
| Ovary | GSE28015 | Cancer | 46 |
| Ovary | GSE3149 | Cancer | 153 |
| Ovary | GSE34405 | Normal | 3 |
| Ovary | GSE6008 | Cancer | 99 |
| Ovary | GSE6008 | Normal | 4 |
| Ovary | GSE68606 | Cancer | 5 |
| Pancreas | E-AFMX-5 | Normal | 4 |
| Pancreas | GSE1133 | Normal | 2 |
| Pancreas | GSE11907 | Cancer | 20 |
| Pancreas | GSE12630 | Cancer | 13 |
| Pancreas | GSE19281 | Cancer | 4 |
| Pancreas | GSE19281 | Normal | 3 |
| Pancreas | GSE2361 | Normal | 1 |
| Pancreas | GSE2719 | Normal | 1 |

| | | | |
|---|---|---|---|
| Pancreas | GSE43288 | Cancer | 4 |
| Pancreas | GSE43288 | Normal | 3 |
| Pharynx | GSE13597 | Cancer | 25 |
| Pharynx | GSE13597 | Normal | 3 |
| Prostate | E-AFMX-5 | Normal | 2 |
| Prostate | E-MEXP-1327 | Cancer | 25 |
| Prostate | E-MEXP-1327 | Normal | 60 |
| Prostate | E-TABM-26 | Cancer | 44 |
| Prostate | E-TABM-26 | Normal | 13 |
| Prostate | GSE1133 | Normal | 2 |
| Prostate | GSE12348 | Normal | 3 |
| Prostate | GSE12630 | Cancer | 11 |
| Prostate | GSE2361 | Normal | 1 |
| Prostate | GSE2443 | Cancer | 20 |
| Prostate | GSE25136 | Cancer | 79 |
| Prostate | GSE2719 | Normal | 1 |
| Prostate | GSE8218 | Cancer | 144 |
| Prostate | GSE8218 | Normal | 4 |
| Skin | E-AFMX-5 | Normal | 2 |
| Skin | GSE1133 | Normal | 2 |
| Skin | GSE11907 | Cancer | 22 |
| Skin | GSE12627 | Cancer | 42 |
| Skin | GSE12630 | Cancer | 17 |
| Skin | GSE1317 | Normal | 1 |
| Skin | GSE2361 | Normal | 1 |
| Skin | GSE2503 | Cancer | 5 |
| Skin | GSE2503 | Normal | 6 |
| Skin | GSE2719 | Normal | 1 |
| Skin | GSE3189 | Cancer | 45 |
| Skin | GSE3189 | Normal | 7 |
| Skin | GSE46517 | Cancer | 31 |
| Skin | GSE46517 | Normal | 7 |
| Skin | GSE4845 | Cancer | 42 |
| Skin | GSE4845 | Normal | 3 |
| Skin | GSE5667 | Normal | 5 |
| Skin | GSE6012 | Normal | 10 |
| Skin | GSE6710 | Cancer | 26 |
| Skin | GSE68606 | Normal | 1 |
| Skin | GSE8401 | Cancer | 83 |
| Skin | GSE8440 | Normal | 9 |
| Skin | GSE9118 | Normal | 1 |
| Skin | GSE9782 | Cancer | 264 |
| Small_Intestine | GSE13083 | Normal | 5 |
| Small_Intestine | GSE2361 | Normal | 1 |
| Small_Intestine | GSE2719 | Normal | 1 |
| Small_Intestine | GSE6272 | Normal | 15 |

| | | | |
|---|---|---|---|
| Soft_Tissue | GSE21124 | Cancer | 149 |
| Soft_Tissue | GSE21124 | Normal | 9 |
| Stomach | GSE12630 | Cancer | 15 |
| Stomach | GSE15460 | Cancer | 31 |
| Stomach | GSE2361 | Normal | 1 |
| Stomach | GSE2719 | Normal | 1 |
| Stomach | GSE29272 | Cancer | 134 |
| Stomach | GSE29272 | Normal | 134 |
| Stomach | GSE37023 | Cancer | 112 |
| Stomach | GSE37023 | Normal | 36 |
| Stomach | GSE6272 | Normal | 16 |
| Stomach | GSE68606 | Cancer | 10 |
| Testis | E-AFMX-5 | Normal | 10 |
| Testis | GSE10615 | Cancer | 27 |
| Testis | GSE10783 | Cancer | 34 |
| Testis | GSE1133 | Normal | 2 |
| Testis | GSE12630 | Cancer | 16 |
| Testis | GSE2361 | Normal | 1 |
| Testis | GSE3218 | Cancer | 107 |
| Thyroid | E-AFMX-5 | Normal | 4 |
| Thyroid | GSE1133 | Normal | 4 |
| Thyroid | GSE12630 | Cancer | 9 |
| Thyroid | GSE2361 | Normal | 1 |
| Thyroid | GSE27155 | Cancer | 12 |
| Thyroid | GSE27155 | Normal | 3 |
| Thyroid | GSE5054 | Cancer | 3 |
| Thyroid | GSE5364 | Cancer | 35 |
| Thyroid | GSE5364 | Normal | 16 |
| Tongue | E-AFMX-5 | Normal | 2 |
| Tongue | GSE1133 | Normal | 2 |
| Tongue | GSE31853 | Normal | 1 |
| Tongue | GSE3524 | Cancer | 16 |
| Tongue | GSE3524 | Normal | 4 |
| Urothelium | GSE3167 | Cancer | 41 |
| Urothelium | GSE3167 | Normal | 9 |
| Urothelium | GSE37317 | Cancer | 19 |
| Urothelium | GSE5287 | Cancer | 30 |
| Uterus | E-AFMX-5 | Normal | 4 |
| Uterus | GSE1133 | Normal | 4 |
| Uterus | GSE11855 | Cancer | 8 |
| Uterus | GSE2152 | Normal | 15 |
| Uterus | GSE2361 | Normal | 1 |
| Uterus | GSE36389 | Cancer | 13 |
| Uterus | GSE36389 | Normal | 7 |
| Uterus | GSE9750 | Cancer | 33 |
| Uterus | GSE9750 | Normal | 24 |
